# Interaction molecular QTL mapping discovers cellular and environmental modifiers of genetic regulatory effects

**DOI:** 10.1101/2023.06.26.546528

**Authors:** Silva Kasela, François Aguet, Sarah Kim-Hellmuth, Brielin C. Brown, Daniel C. Nachun, Russell P. Tracy, Peter Durda, Yongmei Liu, Kent D. Taylor, W. Craig Johnson, David Van Den Berg, Stacey Gabriel, Namrata Gupta, Joshua D. Smith, Thomas W. Blackwell, Jerome I. Rotter, Kristin G. Ardlie, Ani Manichaikul, Stephen S. Rich, R. Graham Barr, Tuuli Lappalainen

**Author notes:** Correspondence to (S.K.), (T.L.). Estonian Genome Centre, Institute of Genomics, University of Tartu, Tartu, Estonia (Silva Kasela).

## Abstract

Bulk tissue molecular quantitative trait loci (QTLs) have been the starting point for interpreting disease-associated variants, while context-specific QTLs show particular relevance for disease. Here, we present the results of mapping interaction QTLs (iQTLs) for cell type, age, and other phenotypic variables in multi-omic, longitudinal data from blood of individuals of diverse ancestries. By modeling the interaction between genotype and estimated cell type proportions, we demonstrate that cell type iQTLs could be considered as proxies for cell type-specific QTL effects. The interpretation of age iQTLs, however, warrants caution as the moderation effect of age on the genotype and molecular phenotype association may be mediated by changes in cell type composition. Finally, we show that cell type iQTLs contribute to cell type-specific enrichment of diseases that, in combination with additional functional data, may guide future functional studies. Overall, this study highlights iQTLs to gain insights into the context-specificity of regulatory effects.

## Introduction

Bulk tissue molecular quantitative trait loci (molQTLs) have been valuable in highlighting potential target genes and gene regulatory mechanisms of disease-associated genetic variants^1–3^. However, context-specific regulatory variants, such as cell type-specific or response QTLs, exhibit particular relevance for disease as compared to standard molQTLs from steady-state tissues^4^. Mapping cell type interaction expression QTLs by modeling the interaction effect between the genotype of a SNP and computationally inferred cell type estimates has shown to aid discovery of cell type-specific effects of expression QTLs^5–7^. Pinpointing the true mediating cell type with this approach may still be challenging due to the properties of the interaction model and correlations between cell type proportions. Thus, rigorous interpretation of cell type iQTLs is important for inferring insights about the true cell type specificity of these effects.

The etiology of most complex diseases is recognized to be influenced both by genetic and environmental factors and their interactions^8^. Detecting gene-environment (GxE) interactions in genome-wide association studies (GWAS) has proven difficult due to small effect sizes and computational challenges^9,10^. Mapping interaction molQTLs for physiological environments, such as age, sex, smoking, or inflammation, offers an opportunity to identify GxE interactions at the molecular level with improved statistical power attributed to stronger effects of regulatory variants. Recently, transcription-based framework has shown the potential to link genes with genetic variant-age interaction to age-associated diseases^11^, suggesting to focus on regulatory variants to study their complex interplay with other factors contributing jointly to variability in traits and diseases.

To comprehensively assess the utility of interaction molQTLs, we performed cell type interaction molQTL (iQTL) mapping from gene expression (RNA-seq) and DNA methylation (EPIC array) in 1,319 participants of diverse ancestries as part of the Trans-Omics for Precision Medicine (TOPMed) program Multi-Ethnic Study of Atherosclerosis (MESA) Multi-Omics pilot with data from two time points (exam 1 and exam 5, ten years apart) (**Figure 1A, Figure S1A**). This longitudinal design enabled us to assess the robustness of cell type iQTLs. Additionally, we characterize the sharing, replication and functional enrichment of cell type iQTLs with respect to their direction of effect. MESA phenotyping data allows us to map age, sex, and smoking iQTLs and study the mediation by cell type iQTLs. Finally, we highlight the informativeness of cell type iQTLs for proposing cell type-specific mechanisms underlying diseases.

**Figure 1.**
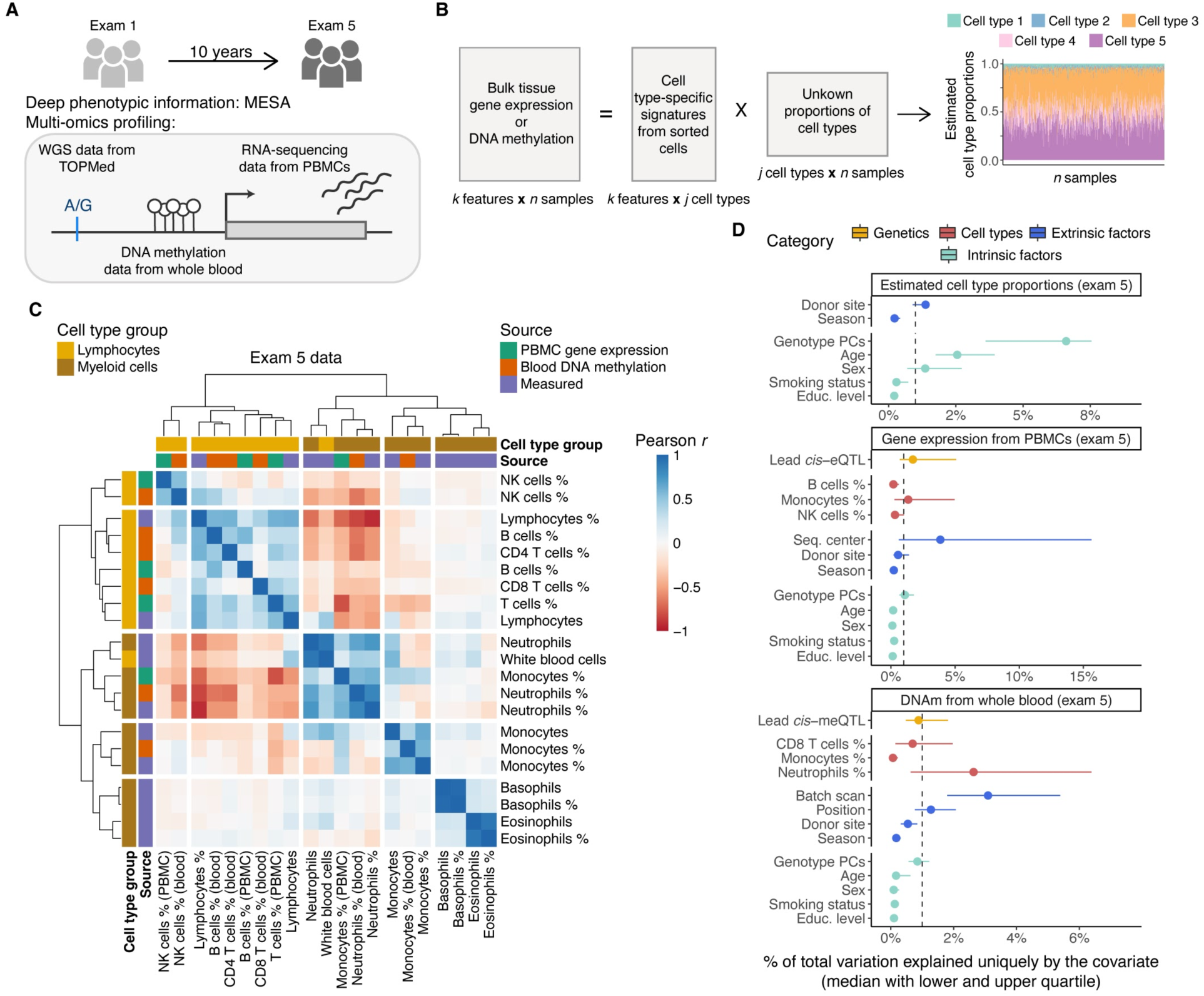
Study design and overview of the estimated cell type proportions. **A)** Illustration of the study design and data types profiled for n = 1,319 individuals. **B)** Graphical illustration of cell type deconvolution. **C)** Correlation of cell type proportions using exam 5 data from three sources: estimated with the CIBERSORT method from PBMC gene expression, estimated with the Houseman method from whole blood DNA methylation, and cell counts measured by flow cytometry. **D)** Sources of variability in estimated cell type proportions with CIBERSORT and the Houseman method, gene expression from PBMCs and DNA methylation from whole blood using exam 5 data. Median of the total variation explained is calculated across all the tested cell types, genes, CpG sites, respectively. Gray dashed line denotes 1% of total variance explained.

## Material and Methods

### MESA Multi-Omics pilot

The Multi-Ethnic Study of Atherosclerosis (MESA) is a prospective cohort study with the goals to identify progression of subclinical atherosclerosis^12^. MESA recruited 6,814 participants at six field centers, ages 45-84 years and free of clinical cardiovascular disease, during 2000-2002. MESA included multiple race/ethnic groups (38% non-Hispanic white, 28% African-American, 22% Hispanic and 12% Asian-Americans), is 53% female, and includes 49% ever-smokers (18% current). All MESA participants provided written informed consent, and the study was approved by the institutional review boards of collaborating institutions.

The MESA Multi-Omics pilot data includes 30x whole genome sequencing (WGS) of ~4,600 individuals through the Trans-Omics for Precision Medicine (TOPMed) Project^13^, with ~1,000 participants samples collected from two time points (exam 1 and exam 5, ten years apart). Whole blood and/or cell types (peripheral blood mononuclear cells (PBMCs), monocytes, T cells) were assayed for transcriptome (RNA-seq), Illumina EPIC methylomics data, plasma targeted and untargeted metabolomics data, and plasma proteomics data. The MESA Multi-Omics pilot biospecimen collection, molecular phenotype data production and quality control (QC) are described in detail in the **Supplemental Material and Methods**.

Here, we analyzed PBMC gene expression data for 19,699 genes from exam 1 (*n* = 931) and exam 5 (*n* = 864), and whole blood DNA methylation (DNAm) data for 747,868 CpG sites from exam 1 (*n* = 900, CpG sites passing QC - 740,291) and/or exam 5 (*n* = 899, CpG sites passing QC - 747,771) together with genotype data from TOPMed Freeze 8^14^.

### Cell type deconvolution

We estimated the cell type composition of PBMC expression and whole blood DNAm by applying two widely used methods - CIBERSORT^15^ and the Houseman method^16^, respectively.

We ran CIBERSORT with default settings, using the LM22 signature matrix provided with the software, on the TPM gene expression matrix containing 2,648 samples from the RNA-seq analysis freeze. We limited our analyses to broad cell types and added the proportions of cell subgroups for B cells, T cells, and NK cells.

We used the Houseman method implemented in the *meffil* R package^17^ together with the whole blood reference from Reinius *et al*.^18^ by using the `meffil.qc` function with “`blood gse35069 complete`” reference applied on the DNAm IDAT files. Importantly, in *meffil* each sample is individually normalized to the cell type reference dataset to avoid dependence between other samples and cell type composition estimates.

For downstream analysis of cell type estimates, we excluded data points per cell type that were more than ±3 standard deviations (SD) from the mean.

### Variability in cell type composition, gene expression, and DNAm

To estimate the unique contribution of different traits to variation in estimated cell type proportions, gene expression, and DNAm, we used fixed effects linear models with no interactions. We applied inverse normal transformation on the response variable (cell type proportions, gene expression levels of autosomal genes, DNA methylation levels of 100,000 randomly selected autosomal CpG sites). More specifically, we used the Type II test for computation of sums of squares (SS) to assess the significance of the main effects^19^ using the *car* package in R. To calculate the proportion of variation uniquely explained by a given trait, we used the eta squared metric by dividing the SS of each term by the total SS.

### Association between estimated cell type proportions and different traits

The effect of various traits on estimated cell type proportions was measured using a linear model. First, we applied inverse normal transformation on the estimated cell type proportions to justify the assumptions of linear regression. To avoid ties, we added random noise from normal distribution N(0, 10^−16^). Second, we leveraged the rich phenotype data available in MESA. We selected traits from 11 different categories, defined as baseline covariates (including age, sex, genotype PCs), anthropometric, smoking habits, alcohol consumption, physical activity, atherosclerosis, blood pressure, inflammation, kidney function, lipids, and lung function. Log-transformation with a pseudo-count of 1 was applied on the molecular traits, data points with >|3| SD from the mean were excluded, and numeric variables were scaled by dividing by two times its SD. This transformation results in comparable coefficients for untransformed binary traits and numeric traits^20^. Third, linear regression was fit with a cell type proportion as response variable and a trait as explanatory variable adjusted for age, sex, self-reported race/ethnicity, educational attainment, site, and month of the exam. If genotype PCs were the traits of interest, then self-reported race/ethnicity was excluded from the covariates list. To adjust for multiple correction, we applied Bonferroni correction and considered associations to be significant, if *P*-value < 0.05 / (# of traits groups × # cell type groups), where the count of cell type groups is equal to 5, corresponding to B cells, T cells/CD4 T cells/CD8 T cells, NK cells, monocytes, and neutrophils.

### Mapping of interaction QTLs (iQTLs)

We mapped interaction QTLs (iQTLs) using tensorQTL^21^. Namely, we fit a linear regression model *Y ~ G + E + GxE + C*, where *Y* is the molecular phenotype (gene expression or DNAm; inverse normal transformed), *G* is the genotype of the genetic variant with MAF > 0.01 in the MESA Multi-Omics pilot data, *E* is the environmental variable (estimated cell type proportions, age, sex, smoking phenotype; mean-centered), *GxE* is the interaction effect between the genotype and environmental variable, and *C* represents additional covariates that correspond to 11 genotype PCs from TOPMed Freeze 8, sex, and PEER factors^22^. Estimation of PEER factors are described in the **Supplemental Material and Methods**. Smoking phenotypes considered as environmental variables were current smoking (binary variable), smoking status (numeric variable with current smokers coded as 2, former smokers as 1, and never smokers as 0), and cotinine levels (inverse normal transformed with random noise added from normal distribution N(0, 10^−16^) to avoid ties).

As for regular QTL mapping in the MESA Multi-Omics data (**Supplemental Material and Methods**), iQTLs were tested for variants ±1Mb of the gene’s TSS or ±500kb of the CpG site. To avoid potential outlier effects in cell type iQTLs, only variants with MAF > 0.05 in the samples belonging to the top and bottom halves of the distribution of estimated cell type proportions were included to the analyses. For age, sex, and smoking iQTLs, we used more stringent MAF filter in the top and bottom halves of interaction values (MAF interaction > 0.1).

To identify genes with significant ieQTLs (ieGenes) or CpG sites with significant imeQTLs (imeSites), the top nominal *P*-values for each molecular phenotype were corrected for multiple testing at the phenotype level using eigenMT^23^, followed by Benjamini-Hochberg procedure across molecular phenotypes. As the significance threshold accounting for multiple testing, we used false discovery rate (FDR) < 0.05 for cell type iQTLs and FDR < 0.25 for trait iQTLs. We further combined significant iQTLs across exams by selecting the molecular phenotype-variant pair with lower interaction *P*-value.

We note that cell type iQTLs could be confounded by factors that affect both the cell type abundance and/or also modify the molQTL effect size, but correcting cell type abundances for these factors and using residualized cell type proportions in the iQTL model rather reduces the study power in most typical scenarios (**Supplemental Note**).

We noticed considerably lower number of monocyte imeQTLs as compared to other cell type imeQTLs, probably attributable to the lower variance in monocyte estimates (SD = 0.02, SD > 0.03 for other cell type estimates). Thus, we only show data related to monocyte imeQTLs on the supplemental figures and tables.

### Direction of iQTL effect

For continuous interaction variables, we grouped the direction of iQTL effect into three categories: 1) positive (increasing) - QTL effect size is positively correlated (increasing) with the interaction variable, 2) negative (decreasing) - QTL effect size is negatively correlated (decreasing) with the interaction variable, 3) uncertain. Assignment of iQTLs into these three categories was done based on the estimates from the linear model. iQTL with nominally non-significant genotype main effect (*P*_G_ > 0.05) was assigned to the “uncertain” group. For clarification, with mean-centered interaction variables, the genotype main effect corresponds to the QTL effect when the interaction variable is 0. Thus, the genotype effect crosses in the middle when plotting interaction variable against molecular phenotype and coloring data points according to the genotype of the iQTL variant. iQTL with nominally significant genotype main effect (*P*_G_ < 0.05) was assigned to the “positive” or “negative” group, if the product of genotype main effect and interaction effect (*β*_G_ x *β*_GxI_) was greater or smaller than 0, respectively.

For binary interaction variables, we fit QTL models separately for both groups. We assigned iQTL into one of the four categories: 1) no effect in one - nominally non-significant genotype effect if one of the groups, 2) magnitude difference - nominally significant genotype effect in both of the groups with the same sign of the estimate, 3) opposite effect - nominally significant genotype effect in both of the groups with the opposite sign of the estimate, 4) uncertain - nominally non-significant genotype effect in both of the groups.

### Sentinel CpG sites for imeQTLs

Using bisulfite DNA sequencing, significant correlation in DNAm between CpG sites (co-methylation) has been observed for short distances up to 1kb, which decreases to baseline after 2kb^24^. To investigate the extent of co-methylation in the EPIC array, we calculated pairwise Pearson correlation coefficients between CpG sites within 500kb on chromosome 22. We used inverse normal transformed DNAm data from exam 5 as an example. We observed that the degree of co-methylation dropped rapidly within 500bp and stayed on average around 0.19 after 1kb. Of note, a similar observation of shorter distances for stronger co-methylation has been previously made based on the Illumina 450K array^25^. Based on this, for the imeQTLs we defined sentinel CpG sites to be used in all the downstream analyses by keeping the CpG site-variant pair with the most significant interaction *P*-value in a 2kb window (±1kb from the most associated CpG site).

### Reproducibility of iQTLs

We leveraged data from two time points in MESA to estimate reproducibility of iQTLs by treating one of the time points as discovery and the other as validation. We calculated the proportion of true positives (*π*_1_)^26^ based on the interaction *P*-value observed in the validation data using the *qvalue* package in R. This metric was used, if more than 20 or more than 100 phenotype-variant pairs were found in the validation data (setting lambda = 0.5 or lambda = 0.85 in the `pi0est()` function, respectively). Additionally, we calculated the fraction of phenotype-variant pairs showing at least nominal significance of the interaction effect in the validation data.

### Sharing of cell type iQTLs

We estimated sharing among the same type of cell type iQTLs using the *π*_1_ statistic as in the reproducibility of iQTLs analysis.

To estimate sharing between cell type ieQTLs and sentinel cell type imeQTLs (FDR < 0.05 in exam 1 or exam 5), we focused on ieVariants that are in LD (*r^2^* ≥ 0.5 within 1Mb) with imeVariants, and vice versa. First, we calculated LD using MESA multi-omics pilot data between the cell type iVariants. Second, for a given query and validation set, we calculated the proportion of variants from the query set to be in LD with the variants from the validation set with the denominator set to the minimum of variants from the query and validation set, termed as the normalized overlap. Third, to estimate the significance of sharing between a cell type ieQTL (query set) and a cell type imeQTL (validation set), or vice versa, we asked whether the query set with positive direction is more likely to overlap with the validation set with positive direction as compared to the validation set with negative direction. For this, we calculated the odds ratio (OR) as the ratio of the odds of the two aforementioned events. To estimate the OR if any cell is equal to zero in the 2×2 table, we applied the Haldane-Anscombe correction^27^ by adding a fixed value of 0.5 to all cells.

### Sharing of cell type iQTLs across populations

To assess the sharing of cell type iQTLs across self-reported race/ethnicity groups in MESA, we leveraged eQTL data from purified cell types from MESA. Namely, expression data from monocytes and T cells were available for a subset of individuals from exam 5 (*n* = 355 and *n* = 362, respectively). We chose monocytes as the cell type of interest for this analysis, because of the high quality of the data. There was more variability among estimated cell type proportions from T cell data. eQTL mapping in monocytes was done following the standard pipeline (**Supplemental Material and Methods**). Monocyte eQTLs were fine-mapped to 95% credible sets of putative causal variants using SuSiE^28^ across all the individuals and by self-reported race/ethnicity groups. Then, we calculated the maximum LD between the monocyte ieQTLs and fine-mapped variants from the credible set across all individuals with expression data from monocytes or by self-reported race/ethnicity. For comparison between self-reported race/ethnicity groups, we focused on whether 1) the ieGene has been fine-mapped to putative causal variants in monocytes, 2) if yes, whether the maximum LD is above or below a specified threshold.

### Replication of cell type ieQTLs in the eQTL Catalogue

We performed replication analysis of cell type ieQTLs in 45 eQTL datasets from purified blood cell types (with and without stimulation) from the eQTL Catalogue^29^. For studies based on microarray technology, if multiple probes per gene existed, we chose the one with the lowest eQTL *P*-value.

We estimated replication using three different metrics: 1) the proportion of true positive (*π*_1_)^26^ using the *qvalue* R package, if more than 20 or more than 100 gene-variant pairs were found in the replication data (setting lambda = 0.5 or lambda = 0.85 in the pi0est() function, respectively); 2) effect size quantified as the absolute value of the median of the genotype effect in replication data; and 3) concordance in allelic direction defined as the proportion of gene-variant pairs having the same directionality of the genotype effect in the replication data and genotype main effect in the cell type iQTL data.

### Functional enrichment analysis

For functional enrichment analysis, we used the registry of candidate *cis*-regulatory elements (cCREs) produced by the ENCODE consortium^30^. The registry V2 consisted of 926,535 human cCREs covering 839 cell and tissue types. We downloaded 61 files representing unique samples with cCREs from various blood cell types, corresponding to 19 unique blood cell types. To maximize data about cCREs available per cell type, we combined data across different samples per cell type. For example, for a H3K27ac-high feature, we required that all samples with H3K27ac data available have an indication of high H3K27ac signal.

To test for the significance of overlap between cell type iQTLs and cCREs, we used the Genomic Annotation Shifter^31^ (GoShifter) method. GoShifter tests for enrichment by locally shifting annotations within the boundaries of associated loci. To generate a null distribution, the shifting process was repeated 10,000 times. As input, we only used independent (sentinel) cell type iQTLs with FDR < 0.05 in exam 1 or exam 5 that had positive or negative direction and provided a list of their LD proxies. To ensure independence of cell type iQTLs, we performed LD pruning with PLINK^32^ in a window of 1,000 variants, sliding by one variant at the time, and with a *r^2^* threshold of 0.1. LD proxies were defined as variants with *r^2^* ≥ 0.8 within 100kb of the cell type iQTLs. LD was calculated based on the unrelated 1,319 individuals from the MESA Multi-Omics pilot.

To quantify the observed enrichment, we used the delta-overlap parameter. Delta-overlap is defined as the difference between the observed proportion of loci overlapping a cCRE and the mean of the proportion of loci overlapping the cCRE under the null. Thus, larger delta-overlap values show stronger enrichment. To estimate the significance of the enrichment we calculated one-sided permutation *P*-value as the proportion of permuted loci overlapping a cCRE is equal to or greater than the observed overlap (adding a pseudo-count of 1 to numerator and denominator). To account for multiple testing, we applied Bonferroni correction method and accounted for the number of target cell types with the given cCRE data available, number of cell types tested for interaction effect, and the number of groups of direction of effect. This was applied separately for each of the tested cCRE and cell type iQTL combination.

### Colocalization analysis of cell type iQTLs

To investigate whether cell type iQTLs provide insights into cell type-specific mechanisms of diseases, we performed colocalization analysis with cell type iQTLs with positive or negative direction and selected diseases/traits. We focused on 7 immunological diseases (asthma^33^, hay fever^33^, Crohn’s disease^34^, inflammatory bowel disease^34^, rheumatoid arthritis^35^, systemic lupus erythematosus^36^, ulcerative colitis^34^) and 3 metabolic traits (HDL cholesterol^37^, LDL cholesterol^37^, triglycerides^37^). We used the harmonized and imputed GWAS summary statistics by the GTEx Consortium^38^. For this analysis we used autosomal cell type ieQTLs with FDR < 0.25 in exam 1 or exam 5 and autosomal sentinel cell type imeQTLs with FDR < 0.05 in exam 1 or exam 5 that had either positive or negative direction of effect.

We performed colocalization analysis with coloc^39^ assuming one causal variant. Coloc was run on a 400kb region centered on each cell type iQTL (±200kb from the iQTL) that had at least one GWAS variant with *P*-value < 10^−5^ within 100kb of the iQTL. Priors were set to *p_1_* = 10^−4^, *p_2_* = 10^−4^, *p_3_* = 5 × 10^−6^ as suggested^40^. As input for cell type iQTL data, we used regression beta and the variance of beta, and for GWAS data, we used the *P*-values. We excluded loci, where the molecular phenotype (TSS of a gene or CpG site) fell into the MHC region, due to complicated LD patterns in this region. Posterior probability for colocalization (PP4) > 0.5 was used as evidence for colocalization. For visualization of colocalized loci, we used locuszoom-like figures with LD calculated based on MESA individuals used for iQTL mapping.

Next, we tested whether we observe more colocalized loci for cell type iQTLs with positive direction and a given disease/trait as compared to height^33^. Height was used as a comparison to account for the enrichment of regulatory variants among trait-associated variants. We calculated the odds ratio (OR) as the ratio of the odds of cell type iQTL colocalizing with a trait of interest to the odds of cell type iQTL colocalizing with height. To estimate the OR if any cell is equal to zero in the 2×2 table, we applied the Haldane-Anscombe correction^27^ by adding a fixed value of 0.5 to all cells. For testing the significance of the OR, we required that at least 10 loci were tested for colocalization with the trait of interest. Bonferroni correction was applied separately for cell type ieQTLs and cell type imeQTLs to account for the number of cell type iQTL and disease/trait pairs used in enrichment testing.

### Mediated moderation

We hypothesized that trait iQTLs may be mediated by GxCell type effects. First, we assessed whether we observe enrichment of cell type iQTL effects among our trait iQTLs (age, sex, smoking iQTLs). For this we evaluated the interaction effect between the genotype of iQTL variant and cell type proportions. Enrichment was estimated using the inflation marker lambda (λ), which is calculated as the ratio of median observed *χ*^2^ test statistic to the median expected *χ*^2^ test statistic under the null.

Second, to formally assess mediation, we formulated the mediated moderation model^41^, where the effect of a moderator (*W*, *e.g.*, age) on the association between the independent variable (*G*, *e.g.*, genotype) and dependent variable (*Y*, *e.g.*, molecular phenotype) is transmitted through a mediator (*M*, *e.g.*, cell type proportion). Structural equation modeling (SEM) has been proposed for the analysis of mediated moderation^41^. As the mediated moderation effect is described by the path *XW* → *XM* → *Y*, we observed in a simulation analysis that similar results can be obtained by using mediation analysis techniques instead of SEM. Thus, we applied mediation analysis using the *mediate* package in R^42^ for added flexibility to account for additional covariates. More precisely, we defined the mediator and outcome models as follows:

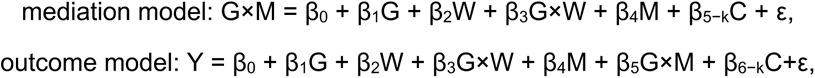

where *C* is the covariates matrix including 11 genotype PCs from TOPMed, sex, and PEER factors. *G*, *M*, and *W* were mean-centered for the mediation analysis.

We estimated the significance of the average causal mediation effect (ACME), average direct effect (ADE), total effect, and proportion of mediated effect by bootstrapping using *k* = 1000 Monte Carlo draws, and 95% confidence intervals were calculated using the bias-corrected and accelerated (BCa) method. *P*-value for ACME < 0.05 was used as an indicator for support for mediation.

### Cell type composition probes

To evaluate whether imeSites are likely associated with cell type composition, we used the “Cell Composition Association Table” from the *FlowSorted.Blood.450k* Bioconductor package^43^. This table summarizes the association between each autosomal probe on the Illumina 450k array that does not contain annotated SNPs and blood cell composition using ANOVA.

## Results

### Cell type composition of blood tissue

We used two methods to characterize the cellular composition of peripheral blood mononuclear cells (PBMCs) from RNA-seq and whole blood from DNA methylation (DNAm) data in MESA — CIBERSORT^15^ and the Houseman method^16^, respectively. These deconvolution methods leverage external purified leukocyte data to infer the proportions of white blood cells (WBC) in heterogeneous tissue samples by modeling bulk tissue data as the sum of weighted cell type-specific expression or DNAm signatures (**Figure 1B**).

Neutrophils were the most abundant cell type in whole blood samples, as expected, but were depleted in PBMC samples where monocytes and T cells constituted a majority of the cell populations (**Figure S1B**). We observed a moderate correlation between the CIBERSORT and Houseman estimates for the same cell type (Pearson correlation 0.42 < *r* < 0.57 in exam 5 data for B cell, NK cell, and T cell comparisons, **Figure S2**). Furthermore, clustering of the cell type abundances showed good concordance between the estimated proportions from different molecular datasets and measured cell type estimates available for a subset of individuals in exam 5 time point (**Figure S1C**). However, more rare cell types, such as eosinophils, were not estimated as accurately as more abundant cell types (**Figure S3)**. Of note, for more abundant cell types, correlation coefficients were similarly high across the different self-reported race/ethnicity groups (**Figure S4)**.

Next, we sought to identify factors that account for variability in cell type composition. Genotype principal components (PCs) that reflect genome-wide genetic effects and ancestry/population structure explained the highest median proportion of variance (~7%), followed by age, sex and donor site (**Figure 1D**). For a more detailed quantification of the unique contributions to total variation in gene expression and DNAm, we studied four categories of factors: 1) *cis* genetics - lead *cis*-molQTLs mapped in MESA (**Supplemental Material and Methods**), 2) cell type composition - estimated cell type proportions, 3) extrinsic technical and/or biological factors - batch variables, donor site, and season, 4) intrinsic biological variables - genotype PCs, age, sex, smoking status, and educational attainment as a proxy for socioeconomic status. The total amount of variability explained by all considered factors varied greatly from ~5% to 90% per gene or CpG site (median of 40% and 20%, respectively, **Figure S5**). Both in gene expression and DNAm data, the largest fraction of inter-sample variation was accounted for by batch variables, estimated cell type proportions, and lead *cis*-molQTLs after controlling for other variables (**Figure 1D**). While the median contribution of intrinsic biological variables was lower compared to other categories, the loci where a large proportion of variation was explained by age or smoking status identified known molecular biomarkers for aging (*e.g., CD248*^44^, *ELOVL2*, and *FHL2*^45^), or smoking (*e.g., AHRR*^46,47^ and *GPR15*^48^) (**Figure S5**). Discovery of age-related differences may be confounded, however, by relative changes in cell type composition due to the impact of age on cell type proportions^43^, which is generalizable for any outcome of interest that correlates with cell type composition. This highlights the importance of accounting for cell type composition as one of the largest sources of variability in studies analyzing gene expression or DNAm.

### Cell type interaction eQTLs and meQTLs in blood

Variability in cell type composition can be exploited to identify cell type interaction QTLs^7,49,50^, where the effect size of the regulatory variants increases (positive direction of effect) or decreases (negative direction of effect) depending on cell type abundance, thus serving as a proxy for cell type-specific QTLs (**Figure 2A**). Applying this framework, we identified cell type interaction *cis*-eQTLs (ieQTLs) for 2,130 genes (out of 19,699, ±1Mb of the transcription start site (TSS)) and cell type interaction *cis*-meQTLs (imeQTLs) for 22,141 CpG sites (out of 747,868, ±500kb of the CpG site) in at least one of the time points with false discovery rate (FDR) < 0.05 across ancestries (**Figure 2B**). Given the correlation in DNAm between proximal CpG sites^24,25^, we defined 20,099 sentinel CpG sites for imeQTLs to represent independent loci by keeping the most significant association in a 2kb window; these were used for further analyses described below.

**Figure 2.**
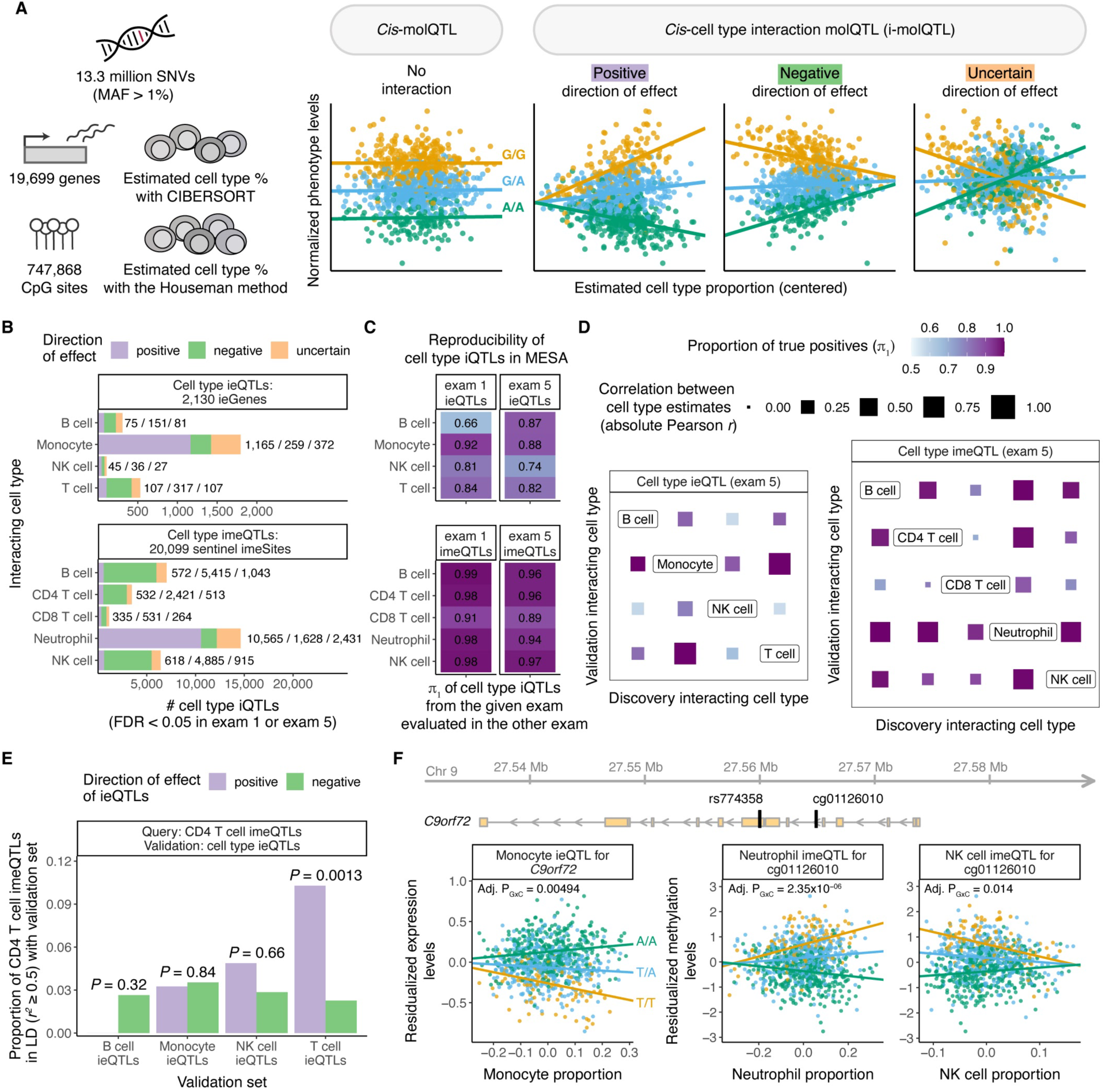
Discovery of cell type ieQTLs and imeQTLs. **A)** Illustration of the approach used to map cell type interaction molQTLs in MESA. **B)** Number of significant cell type ieQTLs and imeQTLs combined across exams (FDR < 0.05 in exam 1 or exam 5 data) stratified by direction of the iQTL effect. **C)** Reproducibility of cell type iQTLs with positive or negative direction of effect using one of the exams for discovery and the other for validation, and vice versa. The proportion of true positives (*π*_1_ statistic) is used as measure of reproducibility. **D)** Sharing among cell type ieQTLs and cell type imeQTLs with positive or negative direction of effect based on exam 5 data quantified as the proportion of true positives (*π*_1_). The size of the square represents the correlation between the two estimated cell type proportions measured using the absolute value of the Pearson correlation coefficient (r). **E)** Sharing between CD4 T cell imeQTLs (query set) and cell type ieQTLs (validation set) combined across exams, quantified as the proportion of CD4 T cell imeQTLs with positive direction in LD (r^2^ ≥ 0.5) with ieQTLs from the given validation set by direction of effect. P-value shows the significance of the odds of CD4 T cell imeQTL with positive direction to overlap with a cell type ieQTL with positive direction as compared to the odds of overlapping with a cell type ieQTL with negative direction. **F)** Example of a cell type iQTL (rs774358) affecting both the expression levels of a gene (C9orf72) and a nearby CpG site (cg01126010).

Discovery of both cell type ieQTLs and imeQTLs was dominated by the most abundant cell type, as previously observed^49^, with the majority of these iQTLs having positive direction of effect (**Figure 2B**). A relatively small percentage of all significant cell type iQTLs (on average, 16.8% across cell type iQTLs and exams) belonged to the ‘uncertain’ group enriched for variants with lower minor allele frequency (MAF) and higher association *P*-values of the interaction effect, indicative of likely false positive results^7^ (**Figure S6**). Using one of the time points as discovery and the other as validation, we observed high reproducibility rates for all cell type iQTLs with either positive or negative direction of effect as an internal quality measure (mean *π*_1_ of 0.84 and 0.96 for cell type ieQTLs and imeQTLs, respectively, **Figure 2C**). Cell type iQTLs with uncertain direction had considerably lower nominal reproducibility rates (**Figure S7**) and were excluded from subsequent analyses.

The MESA cohort design allowed us to investigate population-specific effects of cell type iQTLs. By comparing allele frequency estimates for lead monocyte ieQTLs with positive direction across self-reported race/ethnicity groups, we observed that 0%-14% of ieQTLs did not meet the MAF > 0.01 criteria in one of the specific populations (**Figure S8A**). To study whether the likely causal variants are the same across populations, we leveraged the fine-mapped eQTL data by self-reported race/ethnicity from purified monocytes from MESA exam 5 (**Supplemental Material and Methods**). First, we observed that 66.5%-74.8% of the monocyte ieGenes with positive direction of effect were fine-mapped to likely causal eQTLs in monocytes, with an overlap of 883 (93.1%) ieGenes fine-mapped in at least two self-reported race/ethnicity groups (**Figure S8B**). Second, we calculated LD between the lead ieQTL and fine-mapped variants by self-reported race/ethnicity. While there were considerable differences between the fraction of ieGenes with lead ieQTLs in strong LD (*r*^2^ > 0.5) with fine-mapped eQTLs by groups, these differences were less pronounced when using a more lenient *r*^2^ threshold (**Figure S8C**). This is consistent with the plausible scenario that cell type ieQTLs are largely shared across major ancestral groups when differences in LD and allele frequency are taken into account, as shown for eQTLs^1^.

### Sharing between cell type ieQTLs and imeQTLs

Next, we sought to analyze the extent of sharing of cell type ieQTLs and imeQTLs. We noticed that the iQTLs for highly abundant cell types - monocyte ieQTLs and neutrophil imeQTLs - with negative direction of effect can often be found as an iQTL for another cell type with positive direction of effect, and vice versa (**Figure S9**). In general, the high degree of sharing among cell type ieQTLs and imeQTLs reflected the magnitude of (anti)correlation between estimated cell type proportions (**Figure 2D**), suggesting that cell type iQTLs with specific genetic effects in one (or more) cell types often manifest in other (anti)correlated cell types. We also discovered indications of the same cell type iQTL affecting both expression levels of a gene and DNA methylation levels of a nearby CpG site (**Figure S10, Table S1**). For example, CD4^+^ T cell imeQTLs with positive direction overlapped significantly more often with T cell ieQTLs with positive direction (**Figure 2E**, *P* = 0.0013 as compared to T cell ieQTLs with negative direction). Across 500 unique gene-CpG site pairs associated with the same iQTL (or lead iQTLs in strong LD, *r*^2^ > 0.5), where both the ieQTL and imQTL effect was positive, we observed a discordant genotype main effect for majority of the pairs (64.4%), indicative of mostly negative correlation between gene expression and DNAm as described before for methylation-expression associations (eQTMs)^51^.

An example of a shared cell type iQTL is rs774358 - a variant associated with the expression of *C9orf72* gene and DNAm of the nearby CpG site cg01126010 with the molQTL effect increasing with monocyte and neutrophil abundance, respectively (**Figure 2F**). rs774358 is also a NK cell imeQTL with negative direction of effect, possibly due to negative correlation between the proportion of neutrophils and NK cells in blood. The *C9orf72* repeat expansion is one of the genetic hallmarks of amyotrophic lateral sclerosis (ALS). The expression of *C9orf72* is highest in myeloid cells^52,53^, indicative of myeloid cell-specific molQTL captured by our iQTL approach.

### Cell type specificity of cell type iQTLs

To analyze specificity of cell type ieQTLs by comparing their effects in purified cell types, we leveraged data from the eQTL Catalogue^29^. This resource includes 45 eQTL datasets from various blood cell types with and without stimulation from the lymphocyte and myeloid lineage. We observed, in general, high replication rates for the cell type ieQTLs with positive direction in eQTL data from the corresponding cell (sub)type (max *π*_1_ > 0.8 except for B cell ieQTLs, **Figure S11**), which further manifested as higher median effect size and concordant allelic direction (**Figure 3A, Table S2**). For instance, monocyte ieQTLs with positive direction replicated well in eQTL data from steady-state monocytes as compared to stimulated monocytes, reflecting the need to map response QTLs to discover novel genes with molQTL specific to a cell state. We observed the highest replication rates for T cell ieQTLs with positive direction in different CD4 memory T cell subsets, likely reflecting the shift from naïve to memory T cells with age^54^ in the elderly study subjects from MESA. Importantly, the broad replication patterns matched the corresponding cell type for ieQTLs with positive but not negative direction, with replication in other cell types mirroring the sharing of ieQTLs and (anti)correlation between cell type proportions (**Figure S11**).

**Figure 3.**
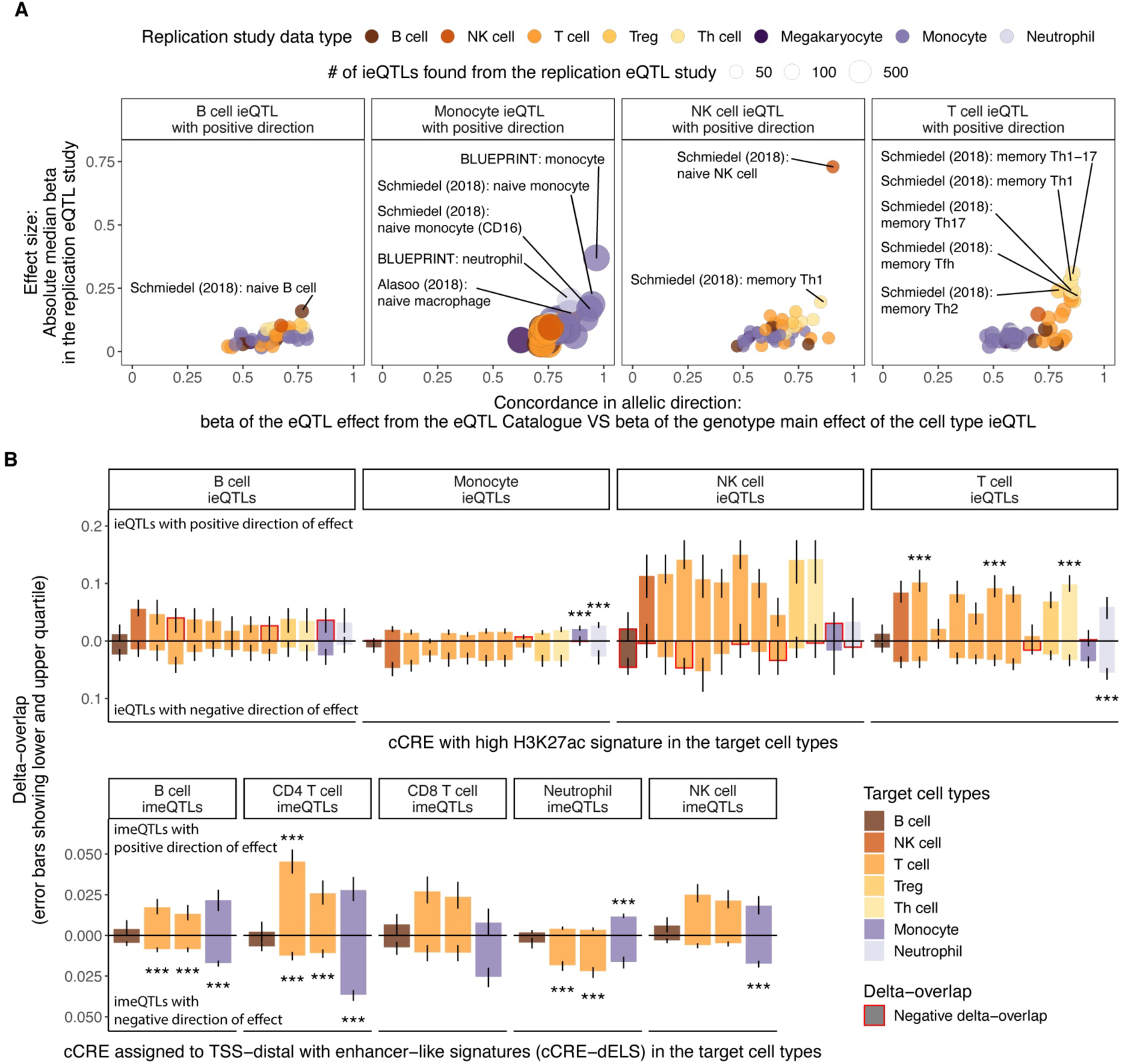
Replication and functional enrichment analysis of cell type iQTLs. **A)** Replication of ieQTLs with positive direction of effect in eQTL datasets from purified cell types from the eQTL Catalogue based on effect size in eQTL data and allelic concordance. Highlighted are up to five datasets with absolute median effect size (beta) > 0.15 in the eQTL dataset and proportion of QTLs with the same allelic direction > 0.75 for B cell ieQTLs or > 0.8 for other cell type ieQTLs. Numerical results for all reference cell types are reported in Table S2. **B)** Functional enrichment analysis with GoShifter showing the delta-overlap, which is the difference between the observed proportion of loci overlapping a cCRE and the null, for cell type ieQTLs (upper panel) overlapping cCRE with high H3K27ac and cell type imeQTLs overlapping cCRE-dELSs. *** - significant association (adjusted P < 0.05) after correcting for the number of target cell types with cCRE data, the number of cell types tested for interaction effect, and the number of groups of direction of effect. Numerical results for all reference cell types are reported in **Table S3**.

*Cis*-eQTLs and *cis*-meQTLs have been shown to be enriched in functional elements of the genome^1,51^. We analyzed the candidate *cis*-regulatory elements (cCREs) from various blood cell types produced by the ENCODE project^30^. After accounting for local genomic structure with GoShifter^31^, we observed highly cell type-specific enrichments of cell type iQTLs with positive direction in distal enhancer-like signatures (cCRE-dELS) and enhancer-associated H3K27ac marks (**Figure 3B, Figure S12, Table S3**), consistent with the tissue-specific nature of enhancers^55,56^. As an example, monocyte ieQTLs were characterized by high H3K27ac in monocytes and neutrophils (the cells of the myeloid phagocyte system^57^) and T cell ieQTLs and CD4^+^ T cell imeQTLs were enriched in T cell subtypes. When focusing on promoter-like signatures (cCRE-PLS), we observed evidence for enrichment of the best powered cell type iQTLs, monocyte ieQTLs and neutrophil imeQTLs with positive direction, in all the five assayed cell types (**Figure S12**). cCRE-PLS was also a highly shared feature in contrast to cCRE-dELS with 64.6% of cCRE-PLSs being present in all and 60.9% of cCRE-dELSs found only in one of the assayed blood cell types.

As exemplified by the results, cell type iQTLs can capture cell type specific effects rather than overall cell type dependence with a good resolution. The interpretation of cell type iQTLs, however, requires consideration of the direction of effect, correlation between cell types, and the quality of the deconvolution. Together, these results support mapping cell type iQTLs as proxies for cell type-specific QTL effects.

### Environmental modifiers of molQTL effect

Next, we leveraged the variation in age, sex, and three smoking phenotypes to quantify the impact of the selected higher-order phenotypes as modifiers of *cis*-QTL effects, *i.e.,* to discover trait iQTLs, where the regulatory variant has a context-specific effect. Compared with cell type iQTLs, trait iQTLs were less abundant. Using a relaxed FDR < 0.25, we identified 277 genes with either age, smoking, or sex interaction eQTLs, and 2,397 CpG sites with either age, smoking, or sex interaction meQTLs (**Figure 4A**). Reproducibility rates between exams added confidence to the robustness of these trait iQTLs (**Figure S13**), as independent replication data is scarce. As an example, we discovered an eQTL for *AHRR* that was significant only in current smokers (**Figure 4B**). Hypomethylation of *AHRR* is one of the most replicated biomarkers for active smoking^58^, and coordinated changes in both DNAm and gene expression across several tissues have been reported^47^.

**Figure 4.**
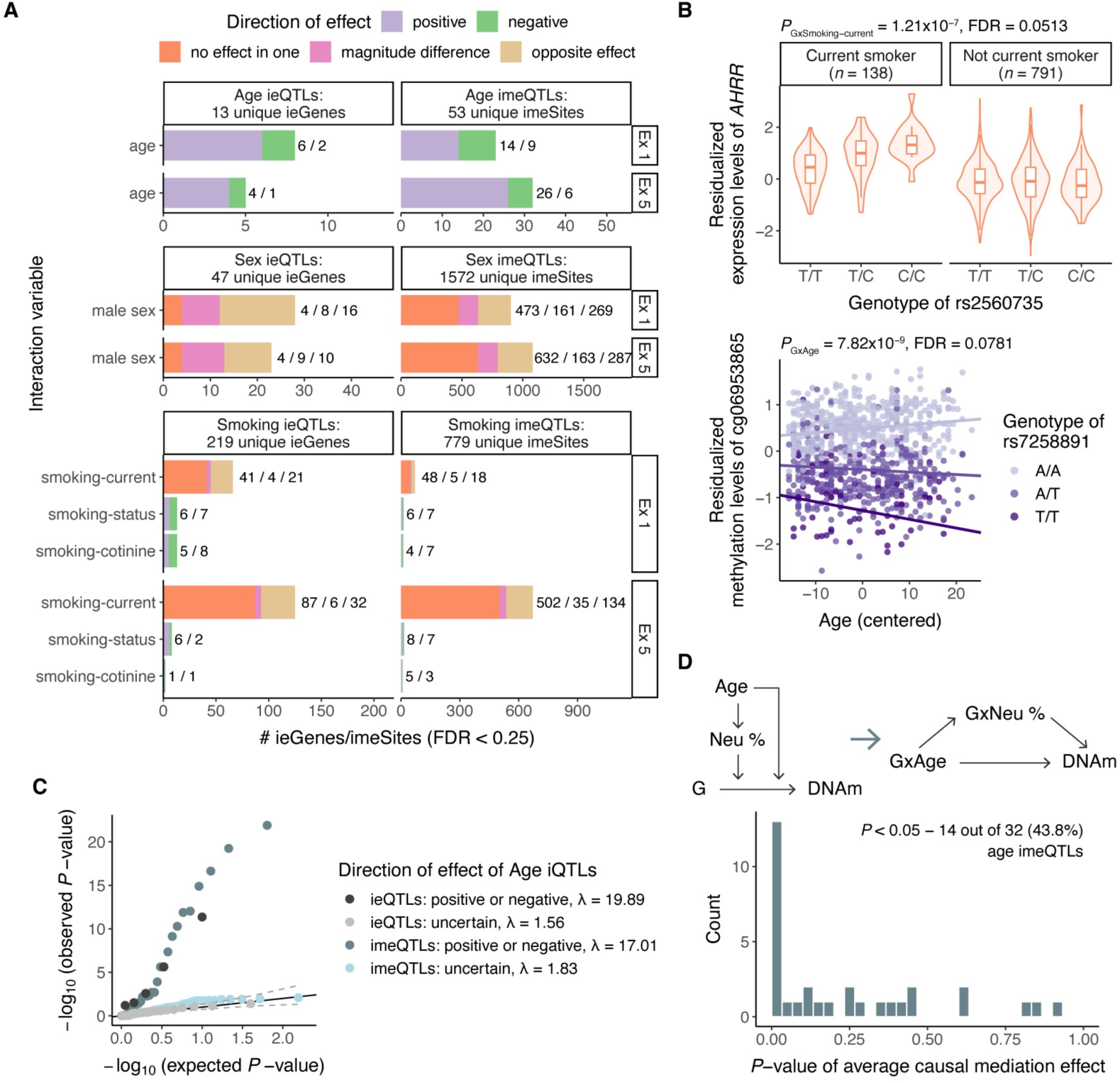
Trait iQTLs and mediated moderation. **A)** Number of significant trait ieQTLs and imeQTLs in exam 1 and exam 5 (FDR < 0.25) by direction of the iQTL effect. **B)** Example of smoking-current ieQTL for AHRR (upper plot) and age imeQTL for cg06953865 (lower plot). **C**) Inflation of GxMonocyte effect among age ieQTLs and GxNeutrophil effect among sentinel age imeQTLs in exam 5 data by direction of age iQTL effect. λ is the inflation factor. **D)** Schema of the mediated moderation approach, where the moderation effect of age on the genotype to DNAm association is mediated by changes in neutrophil proportions. The mediated moderation effect is described by the GxAge -> GxNeutrophil -> DNAm path. P-value histogram of average causal mediation effect (ACME) of GxNeutrophil meditating the GxAge effect on DNAm for 32 age imeQTLs with positive or negative direction.

As observed for cell type iQTLs, a significant GxE term from the interaction model is not specific to the environment tested and may capture effects related to factors correlated with the environment. Across different traits available in MESA, age, sex and smoking are the main non-genetic factors associated with cell type composition (**Figure 1D, Figure S14**), similarly to previous findings^59^. Indeed, we observed a strong enrichment of age iQTLs with positive or negative direction as cell type iQTLs when compared to age iQTLs with uncertain direction of effect as a background (λ = 19.89 vs 1.56 and 17.0 vs 1.83 for GxMonocyte and GxNeutrophil effect in exam 5, respectively, **Figure 4C, Figure S15A,F**), suggesting that some of the age iQTLs may be mediated by cell type iQTLs. While some sex and smoking iQTLs were very strong cell type iQTLs, the evidence for global inflation was weaker (median λ = 2.64 and 1.52 for GxMonocyte and GxNeutrophil interaction in exam 5, respectively, **Figure S15**). This is in line with the finding that the effects of age in DNAm were largely mediated by changes in immune cell proportions, while the effects of sex were typically independent of cellular composition^60^. However, as our cell type iQTL mapping is dominated by the most abundant cell type, we may be underpowered to detect global inflation of interaction with rarer immune cell types in blood.

To formally test for the effect of age iQTLs mediated by cell type iQTLs, we adapted the concept of mediated moderation^41^, where the effect of a moderator (age) on the association between genotype and molecular phenotype is transmitted through a mediator (cell type proportion) (**Figure S16A-B**). We evaluated this hypothesis using neutrophil proportion as the mediator for age imeQTLs in exam 5 as the observed inflation of GxNeutrophil effect was the strongest. As a basis for mediated moderation, neutrophil proportion was positively correlated with age (*r* = 0.14, *P* = 4.81×10^−5^, **Figure S16C**), in line with a reported continuous increase of neutrophils with age^61^. As a result, we observed support for GxNeutrophil mediating the GxAge effect on DNAm for 43.8% (14/32) of age imeQTLs with positive or negative direction of effect (*P*-value of average causal mediation effect (ACME) < 0.05, **Figure 4D, Table S4**) with, on average, 15.5% of the total effect explained by mediator (**Figure S16D**). Of note, the mediation signal was driven primarily by age imeQTLs with positive direction of effect, where 50% (13/26) showed nominal support for mediation. Interestingly, 71.4% (5 out of 7 CpG sites also present on the 450K array) of the imeSites with support for mediation have been identified as CpG sites associated with blood cell composition^43^ (**Figure S17**). As an example, rs7258891 is an age imeQTL for cg06953865 (**Figure 4B**), where we observed strong evidence for mediation (ACME *P* < 0.001). This variant has mainly been associated with various cell count phenotypes, including neutrophil percentage^62^, and the CpG site exhibits different average DNAm across cell types (*P* = 1.33×10^−7^), suggesting that the effect of age on the meQTL is likely mediated by changes in cell type composition differences.

Together, these results suggest that cell type composition changes may confound trait iQTLs by mediating the moderation effect of a trait on genotype and molecular phenotype association, as previously observed for differential expression and differential methylation analysis^63,64^. Thus, an apparent age iQTL effect may arise when a certain cell type proportion varies with age and the regulatory variant has a cell type-specific effect on a molecular phenotype. This warrants caution in interpreting GxE effects on molecular level.

### Cell type iQTLs contribute to immune-mediated inflammatory diseases

Genetic regulatory effects can aid elucidating the tissue specificity of heritable traits and diseases^65^. Given the observed cell type-specific nature of cell type iQTLs with positive direction, we analyzed whether cell type iQTLs provide insights into cell type-specific mechanisms of diseases. We performed colocalization analysis with coloc^39^ of cell type iQTLs (FDR < 0.25 for ieQTLs and FDR < 0.05 for imeQTLs) and a selection of immune diseases and cardiometabolic traits (**Figure S13, Table S5**). When compared with the number of cell type iQTLs colocalizing with height to account for widespread enrichment of QTLs among trait-associated variants^38^, our data confirmed several previously observed cell type-specific enrichments for traits and diseases (**Figure 5A**) - monocytes with lipid traits^66^, B cells with systemic lupus erythematosus^67^, and many different immune cell types, including NK cells, T cells and B cells, with inflammatory bowel disease^68^. Given the varying number of cell type iQTLs with positive direction, we had greater statistical power to detect significant associations involving cell type imeQTLs, particularly neutrophil imeQTLs. Emerging evidence also suggests the contribution of neutrophils in the pathogenesis of autoimmune and inflammatory diseases^69,70^.

**Figure 5.**
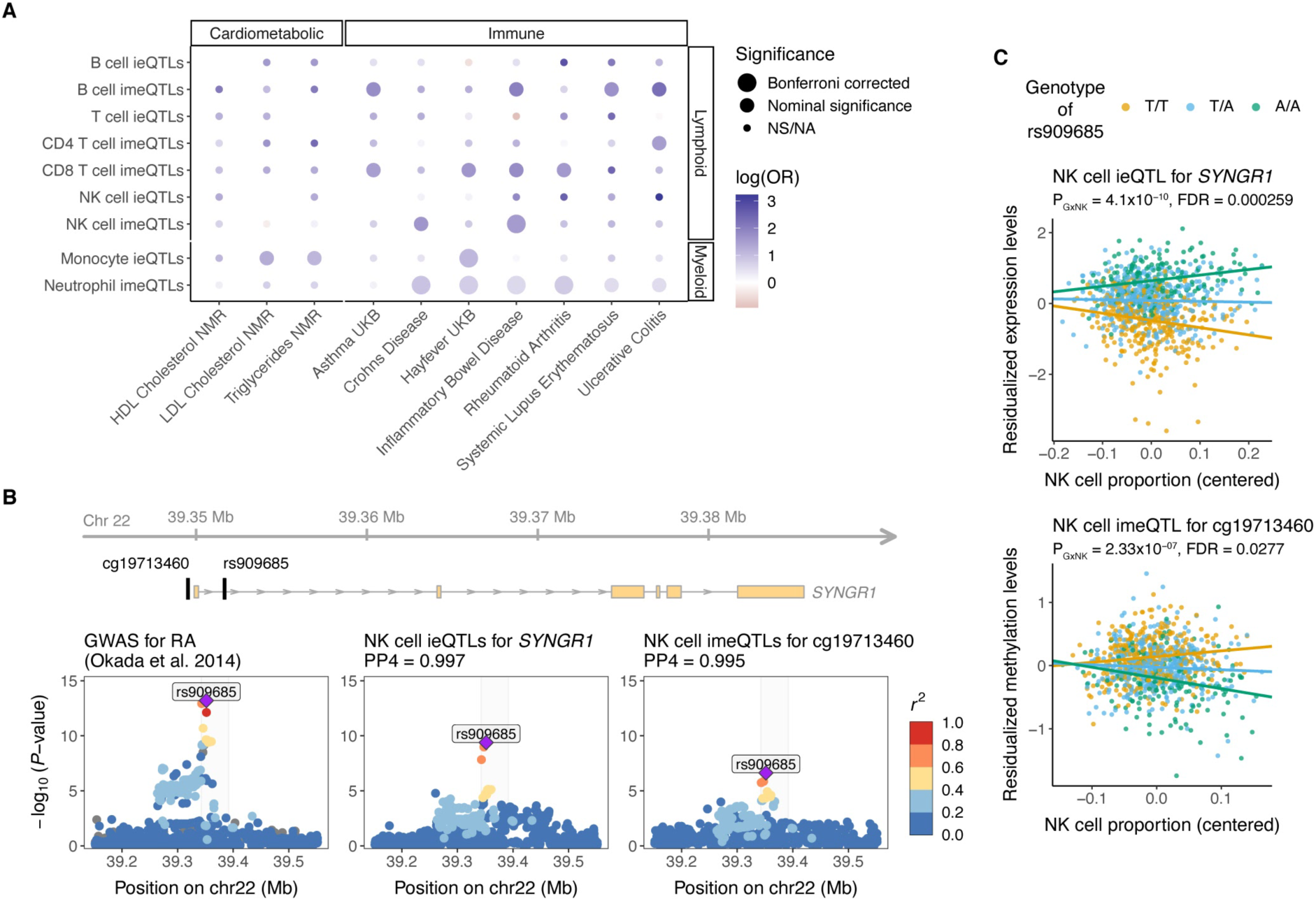
Cell type interaction QTLs and relevance for diseases. **A)** Relevance of cell type ieQTLs (FDR < 0.25) and cell type imeQTLs (FDR < 0.05) for selected cardiometabolic and immune diseases compared to height. For each of the cell type iQTLs, we calculated the odds ratio (OR) as the ratio of the odds for an iQTL to colocalize with cardiometabolic or immune disease to the odds of an iQTL to colocalize with height. For testing the significance of the OR, at least 10 loci tested for colocalization were required, otherwise noted as NA (not available). Bonferroni correction was applied separately for cell type ieQTLs and cell type imeQTLs. NS - not significant. **B)** Colocalization between GWAS for RA and NK cell ieQTLs for SYNGR1 and imeQTLs for a nearby CpG site cg19713460 shown as regional association plots. The highlighted region is depicted at the top and shows the location of the lead GWAS variant for RA, rs909685, and the CpG site relative to the SYNGR1 gene. **C)** Association plot for the NK cell ieQTL for SYNGR1 and the NK cell imeQTL for cg19713460. Dots are colored based on the genotype of rs909685. Data in B) and C) are from exam 1, where we observed the lowest interaction P-values.

In addition to studying disease-specific enrichment, cell type iQTLs can be used to understand the cell type specific mechanism of a disease-associated variant. For instance, the A allele of rs909685 (T/A), located in the intron of the synaptogyrin-1 (*SYNGR1*) gene, has been shown to increase the susceptibility to rheumatoid arthritis (RA) for individuals of European, Asian, and African ancestries^35,71,72^. In our data, rs909685 was associated with *SYNGR1* expression, with the effect size increasing with NK cell and T cell proportions, and decreasing with monocyte proportion (**Figure 5B, Figure S14A**). rs909685 was also associated with the methylation levels of cg19713460, located in the promoter region of *SYNGR1* (400bp from the transcription start site (TSS)), with the effect size increasing with NK cell proportion (**Figure 5B**). Of note, rs909685 A allele was associated with higher expression levels of *SYNGR1* and lower methylation levels of cg19713460 (**Figure 5C**, **Figure S14B**). For the cell type iQTLs, we observed very strong evidence for colocalization with the RA GWAS signal (PP4 > 0.99). Interestingly, rs909685 falls into the cCRE that is characterized by high DNase and H3K27ac in NK cells, CD8^+^ T cells, and B cells (**Figure S14C**). Furthermore, the *SYNGR1* knockdown lowered the release of pro-inflammatory cytokines or chemokines (*e.g.,* IFN-γ, TNF, and RANTES) by activated NK cells, suggesting a functional role of *SYNGR1* in NK cells^73^. Together, these data suggest that rs909685 influences susceptibility to RA via NK cell-specific action, captured by our cell type iQTLs integrated with functional annotation data. As a likely mechanism, a causal chain from methylation of the promoter of *SYNGR1* to affecting mRNA expression resulting in affected RA risk has been proposed^74^. This example highlights the usefulness of incorporating cell type iQTLs and functional data into investigations of cell type-specific mechanisms of disease-associated variants.

## Discussion

We performed interaction QTL mapping with cell type abundance, age, sex, and smoking as the environmental factors to identify regulatory variants with plasticity in effect size rather than constant molecular effects. While a sample size of ~900 individuals of multi-ethnic background at two time points was sufficient to map cell type iQTLs for a large number of genes and CpG sites, discovery of molQTLs interacting with higher order physiological traits were limited. Given the unique aspects of our study design, we were able to assess the reproducibility of the iQTLs between time points to demonstrate the robustness of the results, and highlight the sharing between cell type ieQTLs and imeQTLs characterized mostly by negative correlation between gene expression and DNAm and discordant genotype main effect. Importantly, the interpretation of cell type iQTLs depends on several factors - direction of effect, correlation between cell types within the tissue, and resolution of the cell type deconvolution. Our results suggest that biologically most informative results are obtained for molQTLs when the effect size is increasing (positive direction) with the most abundant cell type in the tissue.

Even though cell type iQTLs cannot be considered cell type-specific *per se*, cell type iQTLs with positive direction replicate well in eQTL datasets from purified cell types and show enrichment in cCREs from the interacting (or similar) cell type. We demonstrated this concept in whole blood, which had the necessary cell type-specific eQTL replication data. Our results show promise for interaction QTL approaches for identification of cell type-specific QTLs in other tissues where single-cell or cell type-specific data are not available or easily acquired. Moreover, cell type iQTLs combined with functional annotations of the genome can help prioritize cell types for functional follow-up studies.

molQTLs with GxE interactions at the molecular level hold the promise to guide discovery of GxE interactions in complex diseases^75–80^. These loci may mark the genetic component of inter-individual variation in response to different environments or physiological states, including disease, thus contributing to phenotypic variation in humans. Our results with age imeQTLs, however, suggest that cell type composition changes may partly mediate the moderation effect of age. Similar observations have previously been made for sex-biased *cis*-eQTLs^81^, yet the confounding effect that cell type composition has on molQTL effect size variation has not been appreciated to the same extent as in differential expression and methylation studies, particularly in epigenome-wide association studies^43,64^.

Based on our results, we propose that mediation by cell type composition is the primary starting hypothesis for molQTLs with GxE effects, and this should be explicitly ruled out before postulating other molecular moderation mechanisms. We further hypothesize that when trait iQTL and GWAS signal colocalize, only molQTLs with GxE not mediated by cell types would have a GxE interaction at the GWAS level - whilst molQTLs with support for mediation most likely are subject to confounding. Future studies with larger sample sizes will be needed to properly evaluate this hypothesis.

Overall, the integration of genomic data with functional multi-omic data in large and diverse longitudinal cohorts offers an opportunity to map genetic effects on molecular traits, and to study its complex interplay with other environmental factors. Our study shows the value of mapping interaction QTLs as a feasible computational approach to obtain insights into the context-specificity of regulatory effects.

## Acknowledgements

We thank all members of the Lappalainen laboratory for valuable discussions and support. We also thank Paul J. Hoffman, Grant T. Hiura, Kristina L. Buschur, Sailalitha Bollepalli, and Inga-Maria Launonen for their input and advice on earlier versions of this manuscript. S.K., R.G.B. and T.L. were supported by NIH/NHLBI grant R01HL142028. S.K.H. is supported by Marie-Skłodowska Curie fellowship H2020 grant 706636, Helmholtz Young Investigator grant VH-NG-1620 and DFG Emmy Noether Programme grant KI 2091/2-1. B.C.B. is supported by NIH/NHGRI grant K99HG012373. T.L. is supported by NIH/NIMH grant R01MH106842, NIH/NHGRI grant UM1HG008901 and NIH/NIGMS grant R01GM122924. Whole genome sequencing (WGS) for the Trans-Omics in Precision Medicine (TOPMed) program was supported by the National Heart, Lung and Blood Institute (NHLBI). WGS for “NHLBI TOPMed: Multi-Ethnic Study of Atherosclerosis (MESA)” (phs001416.v1.p1) was performed at the Broad Institute of MIT and Harvard (3U54HG003067-13S1). Centralized read mapping and genotype calling, along with variant quality metrics and filtering were provided by the TOPMed Informatics Research Center (3R01HL-117626-02S1). Phenotype harmonization, data management, sample-identity QC, and general study coordination, were provided by the TOPMed Data Coordinating Center (3R01HL-120393-02S1), and TOPMed MESA Multi-Omics (HHSN2682015000031/HSN26800004). The MESA projects are conducted and supported by the National Heart, Lung, and Blood Institute (NHLBI) in collaboration with MESA investigators. Support for the Multi-Ethnic Study of Atherosclerosis (MESA) projects are conducted and supported by the National Heart, Lung, and Blood Institute (NHLBI) in collaboration with MESA investigators. Support for MESA is provided by contracts 75N92020D00001, HHSN268201500003I, N01-HC-95159, 75N92020D00005, N01-HC-95160, 75N92020D00002, N01-HC-95161, 75N92020D00003, N01-HC-95162, 75N92020D00006, N01-HC-95163, 75N92020D00004, N01-HC-95164, 75N92020D00007, N01-HC-95165, N01-HC-95166, N01-HC-95167, N01-HC-95168, N01-HC-95169, UL1-TR-000040, UL1-TR-001079, UL1-TR-001420, UL1TR001881, DK063491, and R01HL105756. The MESA Lung Study is supported by R01-HL077612 and R01-HL093081. Harmonization of spirometry traits were supported by NIH/NHLBI R21-HL121457, R21-HL129924, and K23-HL130627. This publication was developed under the Science to Achieve Results (STAR) research assistance agreements, No. RD831697 (MESA Air) and RD-83830001 (MESA Air Next Stage), awarded by the U.S Environmental Protection Agency (EPA). It has not been formally reviewed by the EPA. The views expressed in this document are solely those of the authors and the EPA does not endorse any products or commercial services mentioned in this publication. The authors thank the other investigators, the staff, and the participants of the MESA study for their valuable contributions. A full list of participating MESA investigators and institutes can be found at http://www.mesa-nhlbi.org.

## Author Contributions

S.K. and T.L. designed the study. S.K. performed the analyses. F.A., S.K.H., B.C.B., and D.C.N. contributed to the analyses of the data. J.I.R., S.S.R., R.P.T., P.D., Y.L., K.D.T., W.C.J., D.V.D.B., S.G., N.G., J.D.S., and T.W.B. were involved in the acquisition and processing of data. T.L., R.G.B., S.S.R., A.M., and K.G.A. supervised the work. S.K. and T.L. wrote the first draft. F.A., S.S.R., A.M., S.K.H., and B.C.B. contributed to the editing of the manuscript. All authors approved the final version of the manuscript.

## Data and code availability

MESA data are available through dbGaP (accession number phs001416). The full summary statistics of interaction molQTLs generated during this study are available at xxx (*will be shared after publication of this manuscript*). Example code for mapping iQTLs with tensorQTL is available at https://github.com/broadinstitute/tensorqtl.

