## Supplemental information for "Interaction molecular QTL mapping discovers cellular and environmental modifiers of genetic regulatory effects"

### Supplemental Figures

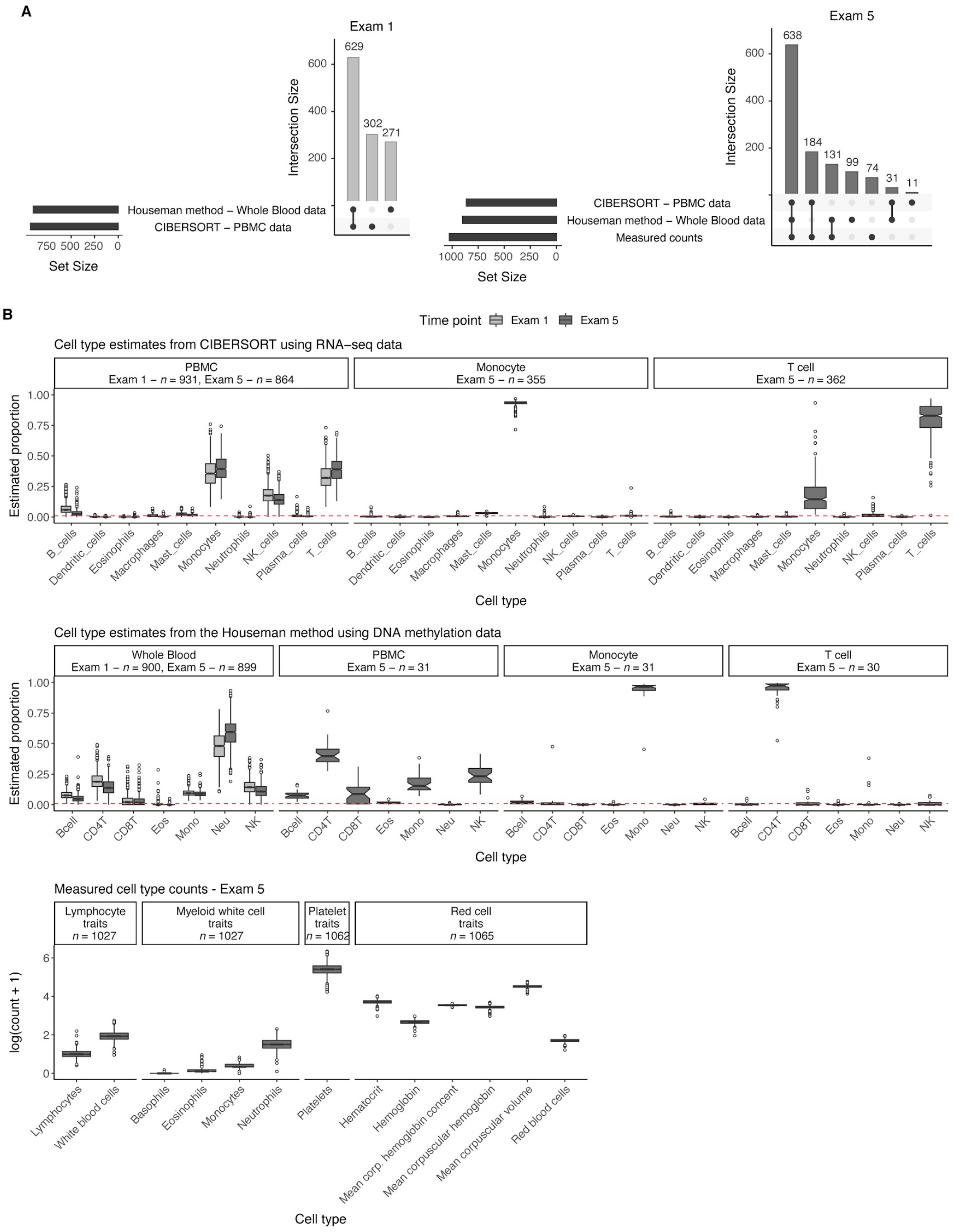


**Figure S1. Estimated and measured cell type proportions. A)** Sample overlap between cell type estimates by time point. **B)** Boxplots showing the proportion of estimated cell type proportions by the cell type deconvolution method or showing the measured counts available for a subset of the subjects in MESA multi-omics pilot from exam 5. Red dashed line denotes estimated proportion = 0.01.

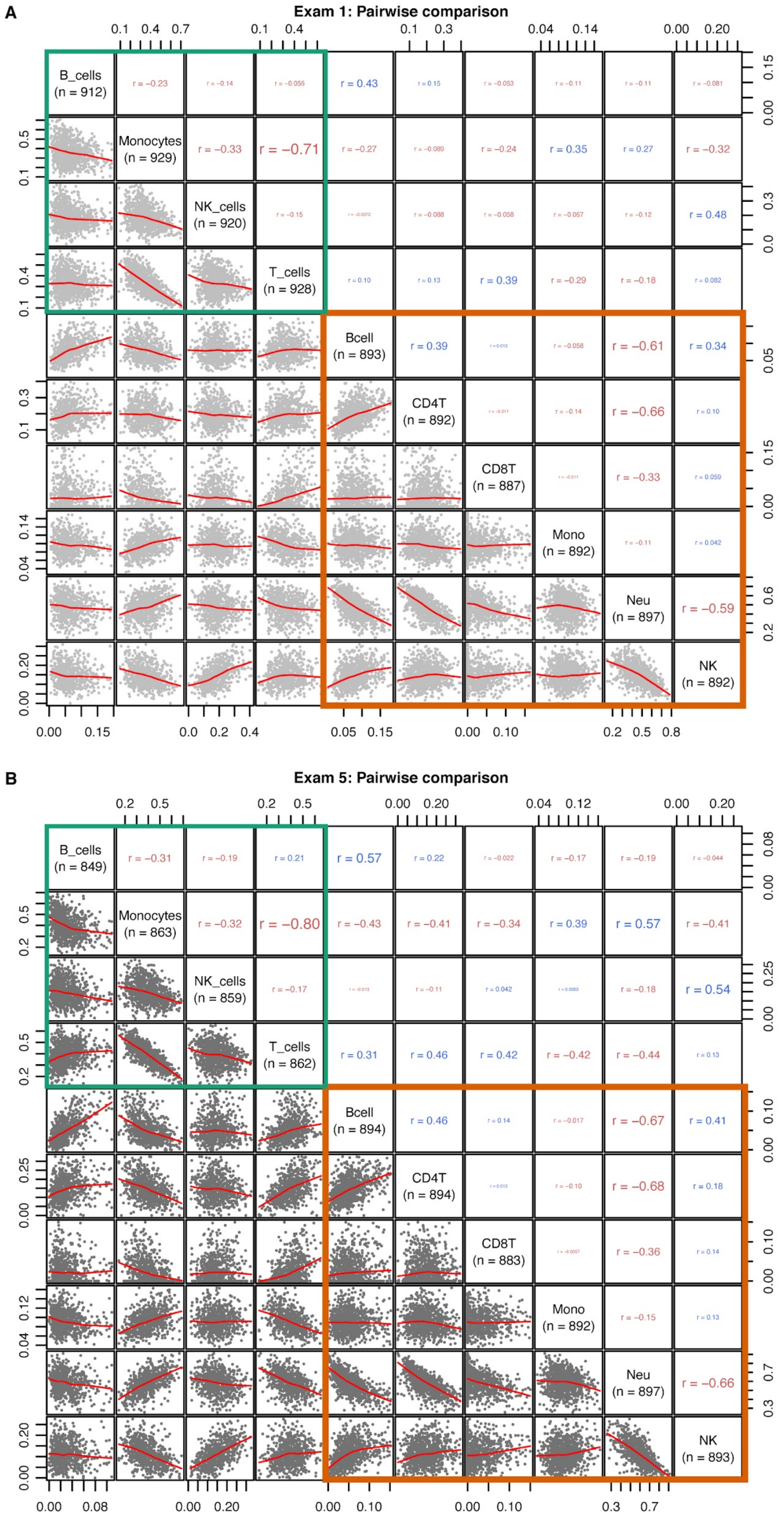


**Figure S2. Correlation between estimated cell type proportions. A)-B)** Pearson correlation coefficient between estimated cell type proportions within and across the two deconvolution methods for the two timepoints - **A)** Exam 1 and **B)** Exam 5. Green square denotes the correlations between estimated cell type proportions from PBMC RNA-seq with the CIBERSORT method and orange square denotes the correlations between estimated cell type proportions from whole blood DNA methylation with the Houseman method. Red line shows the LOWESS line using locally-weighted polynomial regression. Excluded data points per cell type that are more than ±3 SD from the mean.

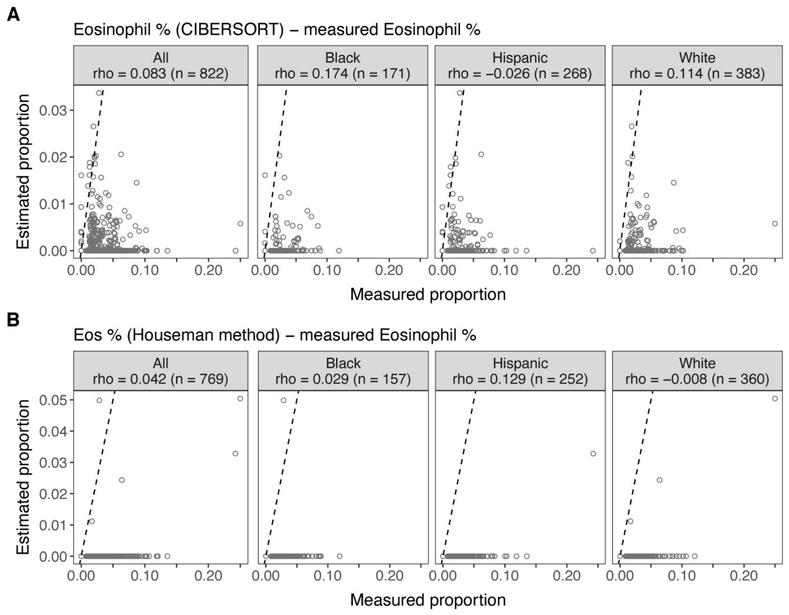


**Figure S3. Correlation between estimated and measured eosinophils proportions by self-reported race/ethnicity. A)-B)** Scatter plot of estimated and measured eosinophils proportions with the CIBERSORT method from PBMC RNA-seq (**A**) and with the Houseman method from whole blood DNA methylation (**B**) using Exam 5 data. Measured eosinophils proportions are calculated by dividing the given relative count by the total WBC count.

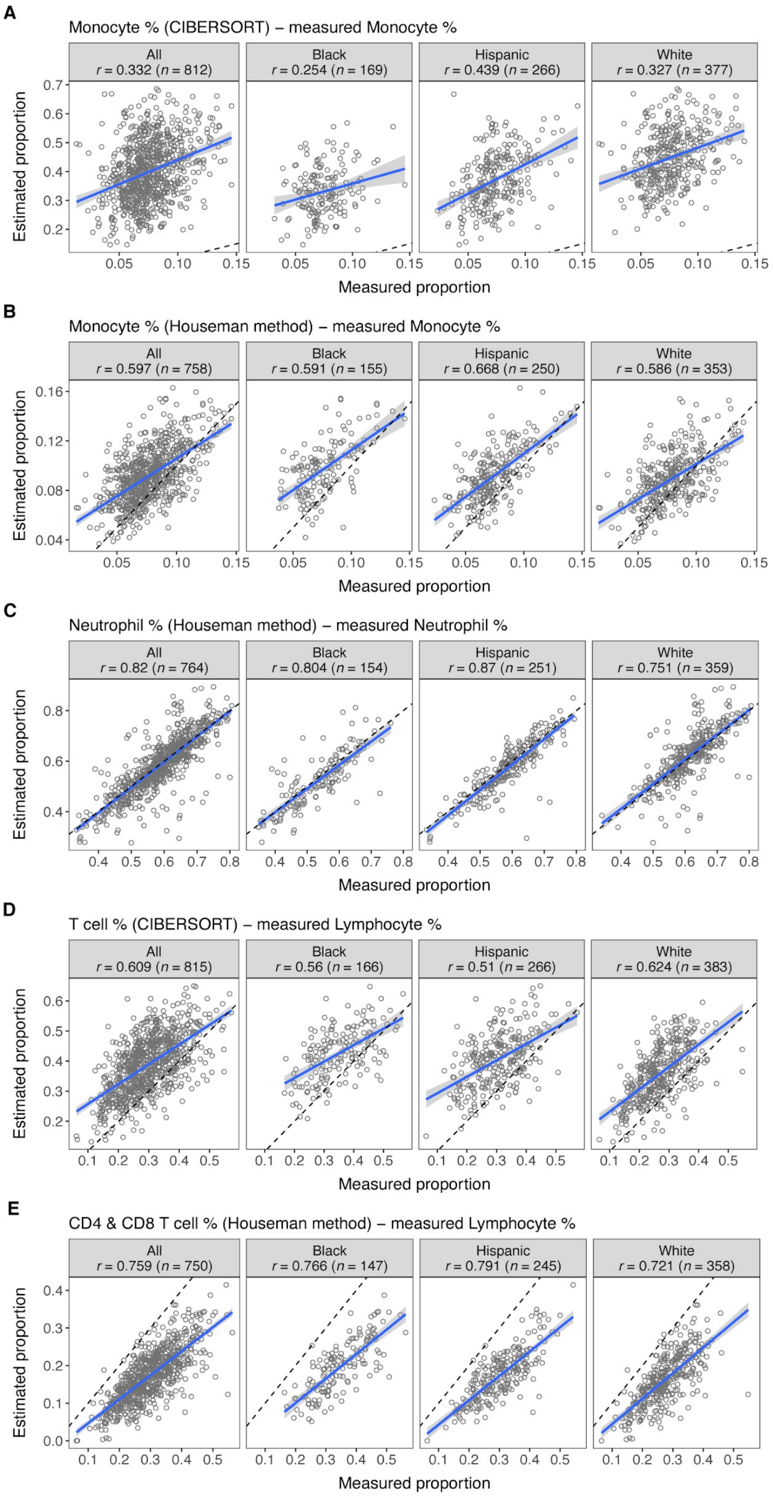


**Figure S4. Correlation between estimated and measured cell type proportions by self-reported race/ethnicity for the most abundant cell types. A)-E)** Scatter plot of estimated and measured cell type proportions using Exam 5 data. Measured cell type proportions of the white blood cell (WBC) subtypes are calculated by dividing the given relative count by the total WBC count. Blue line shows the linear regression fit. Excluded data points per cell type that are >3SD from the mean. The blue line shows the linear fit and shading shows the confidence interval.

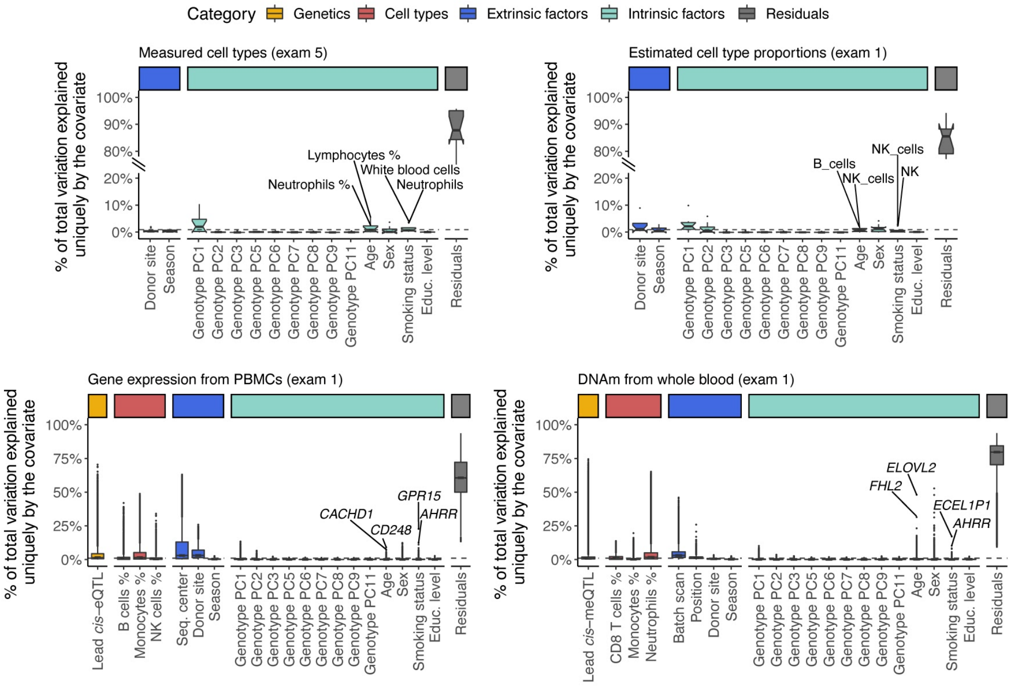


**Figure S5. Sources of variability in cell type proportions, gene expression and DNA methylation data**. Sources of variability in measured cell type proportions (exam 5), estimated cell type proportions (exam 1), gene expression from PBMCs (exam 1) and DNA methylation from whole blood (exam 1). Gray dashed line denotes 1% of total variation explained. Lead *cis*-eQTLs and *cis*-meQTLs are estimated from MESA.

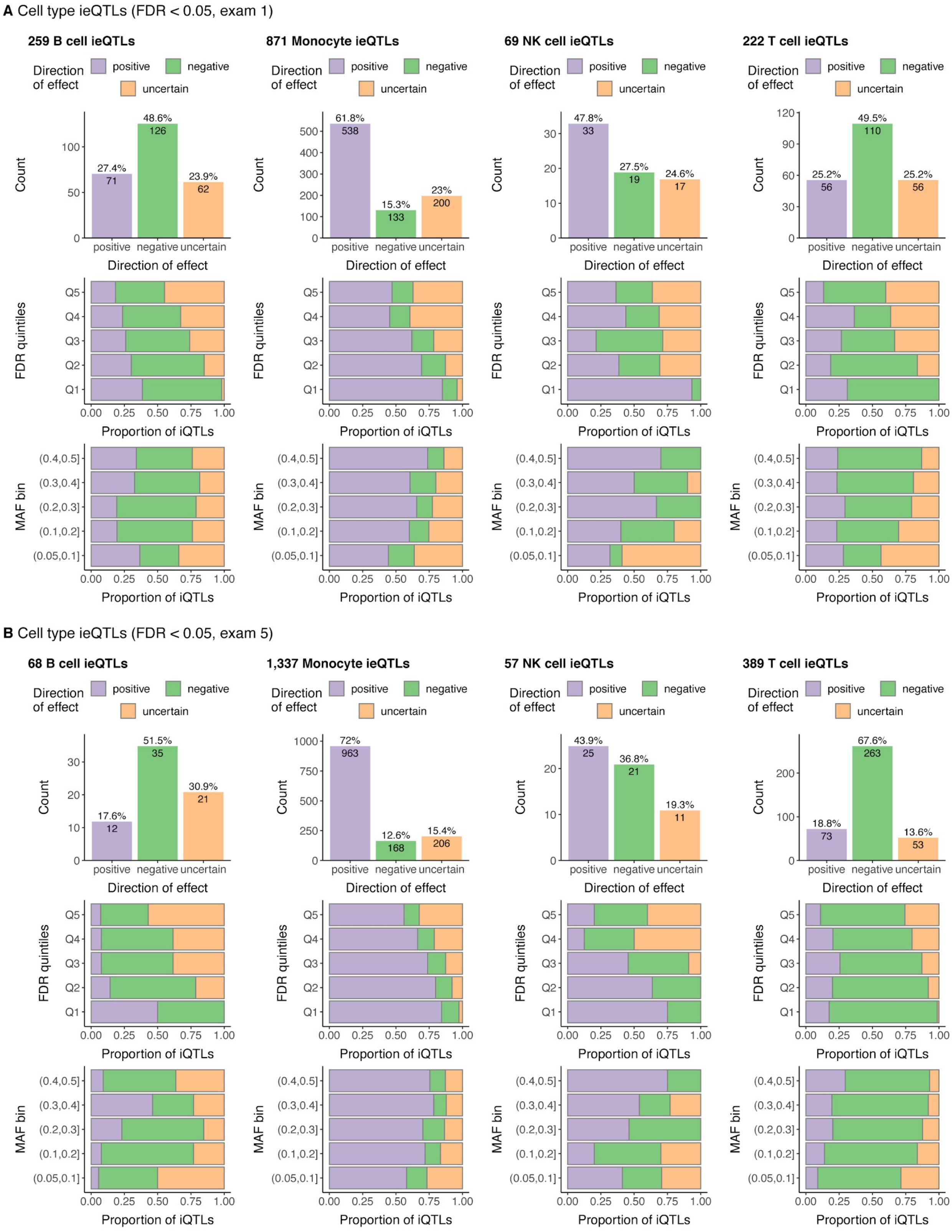
A Cell type ieQTLs (FDR < 0.05, exam 1)
259 B cell ieQTLs
871 Monocyte ieQTLs
69 NK cell ieQTLs
222 T cell ieQTLs
Direction of effect
positive
negative
uncertain
Count
27.4%
71
48.6%
126
23.9%
62
61.8%
538
15.3%
133
23%
200
47.8%
33
27.5%
19
24.6%
17
25.2%
56
49.5%
110
25.2%
56
FDR quintiles
Q5
Q4
Q3
Q2
Q1
Proportion of iQTLs
MAF bin
(0.4,0.5]
(0.3,0.4]
(0.2,0.3]
(0.1,0.2]
(0.05,0.1]
0.00
0.25
0.50
0.75
1.00
B Cell type ieQTLs (FDR < 0.05, exam 5)
68 B cell ieQTLs
1,337 Monocyte ieQTLs
57 NK cell ieQTLs
389 T cell ieQTLs
17.6%
12
51.5%
35
30.9%
21
72%
963
12.6%
168
15.4%
206
43.9%
25
36.8%
21
19.3%
11
18.8%
73
67.6%
263
13.6%
53

**C** Cell type imeQTLs (FDR < 0.05, exam 1)

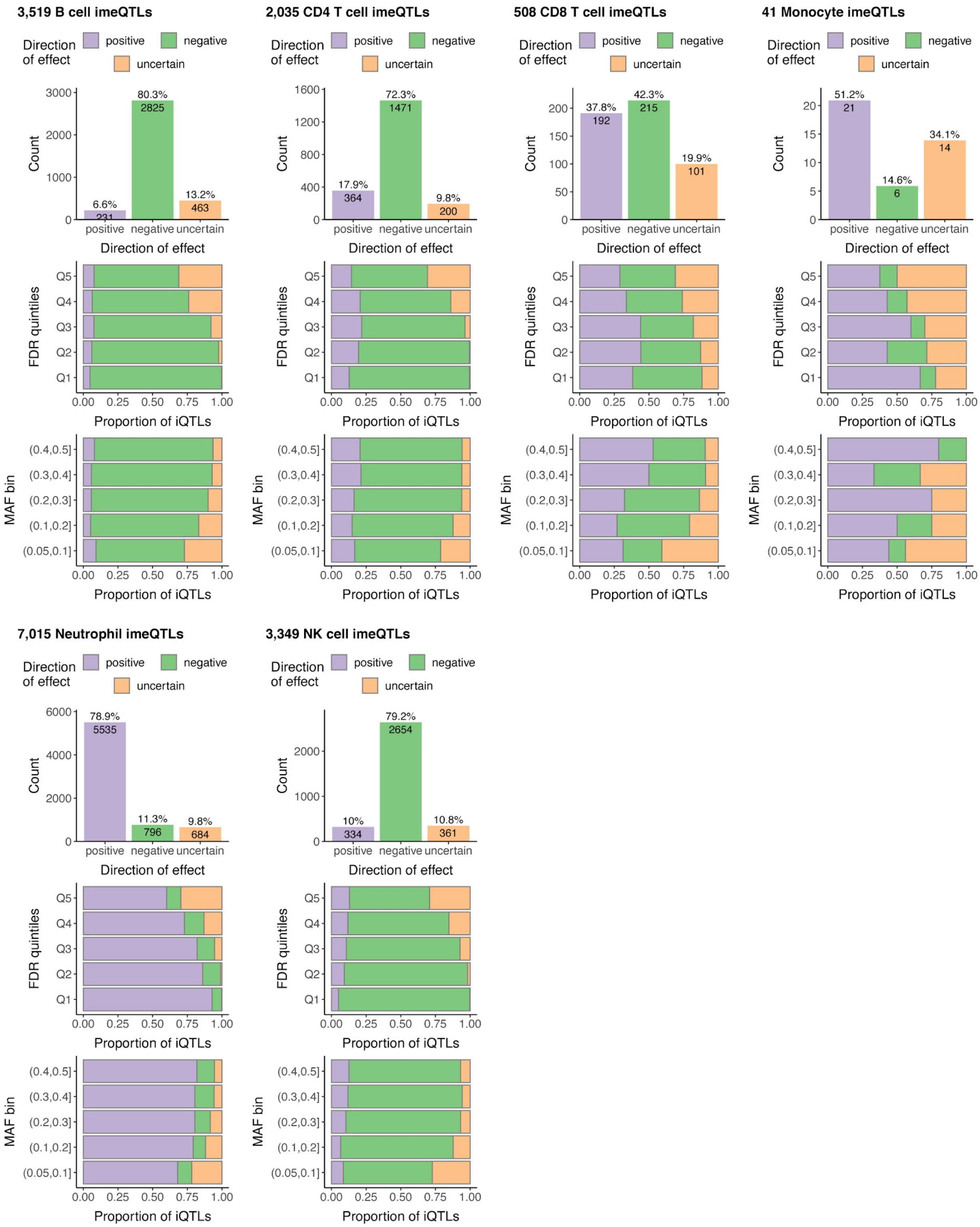

D Cell type imeQTLs (FDR < 0.05, exam 5)

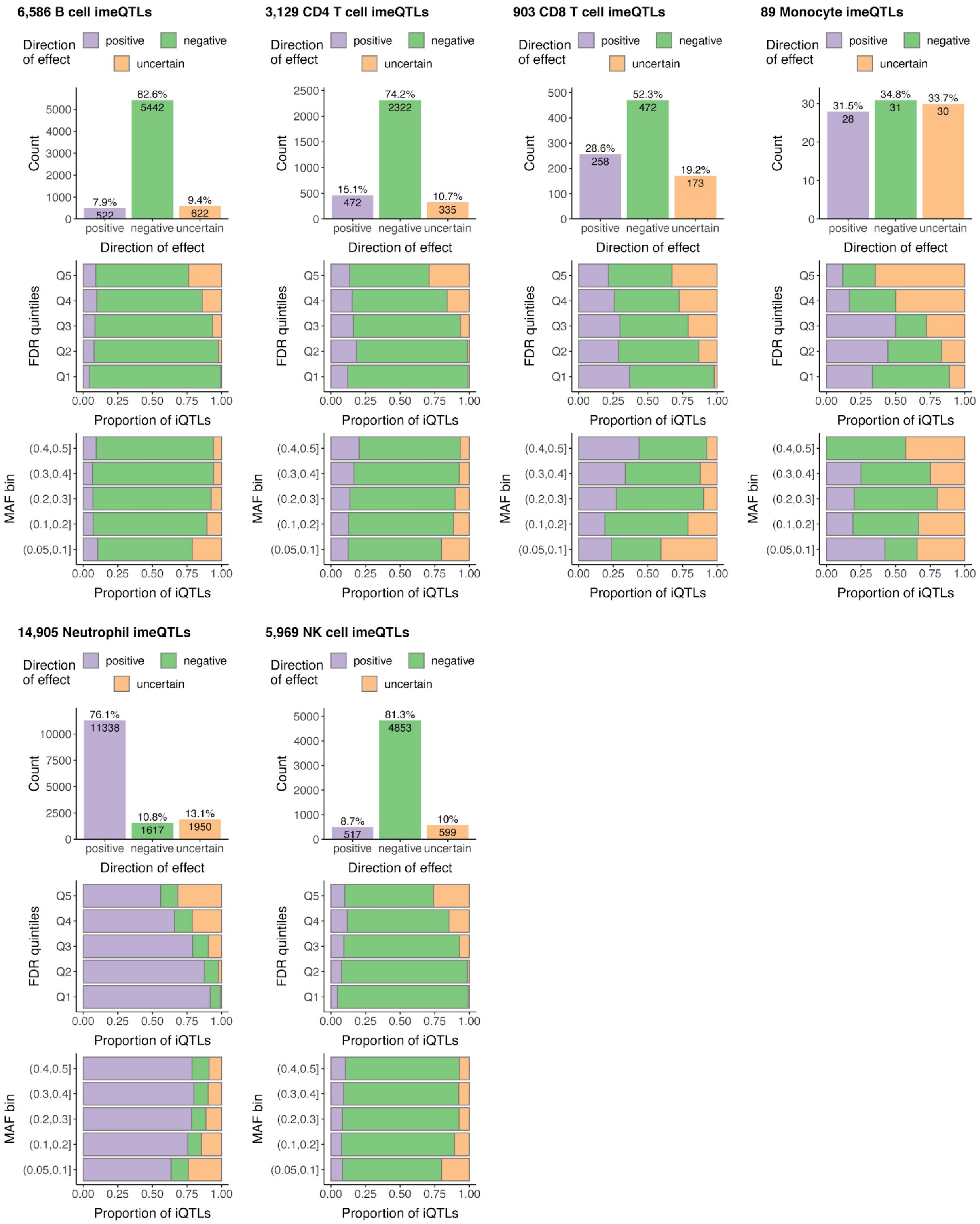


**Figure S6. Discovery of cell type interaction eQTLs (A-B) and cell type interaction meQTLs (C-D) in exam 1 and exam 5 data. A)-D)** Overview of cell type iQTLs (including also non-sentinel imeQTLs) with FDR < 0.05 by direction of effect (top panel), the proportion of iQTLs with different direction of effect by FDR quintiles (Q1 - the most significant bin, Q5 - the least significant bin) (middle panel), and the proportion of iQTLs with different direction of effect by MAF bins (bottom panel). Of the significant cell type iQTLs across exams and including Monocyte imeQTLs, 18.5% belong to the uncertain group enriched for variants with lower MAF and higher association *P*-values of the interaction effect.

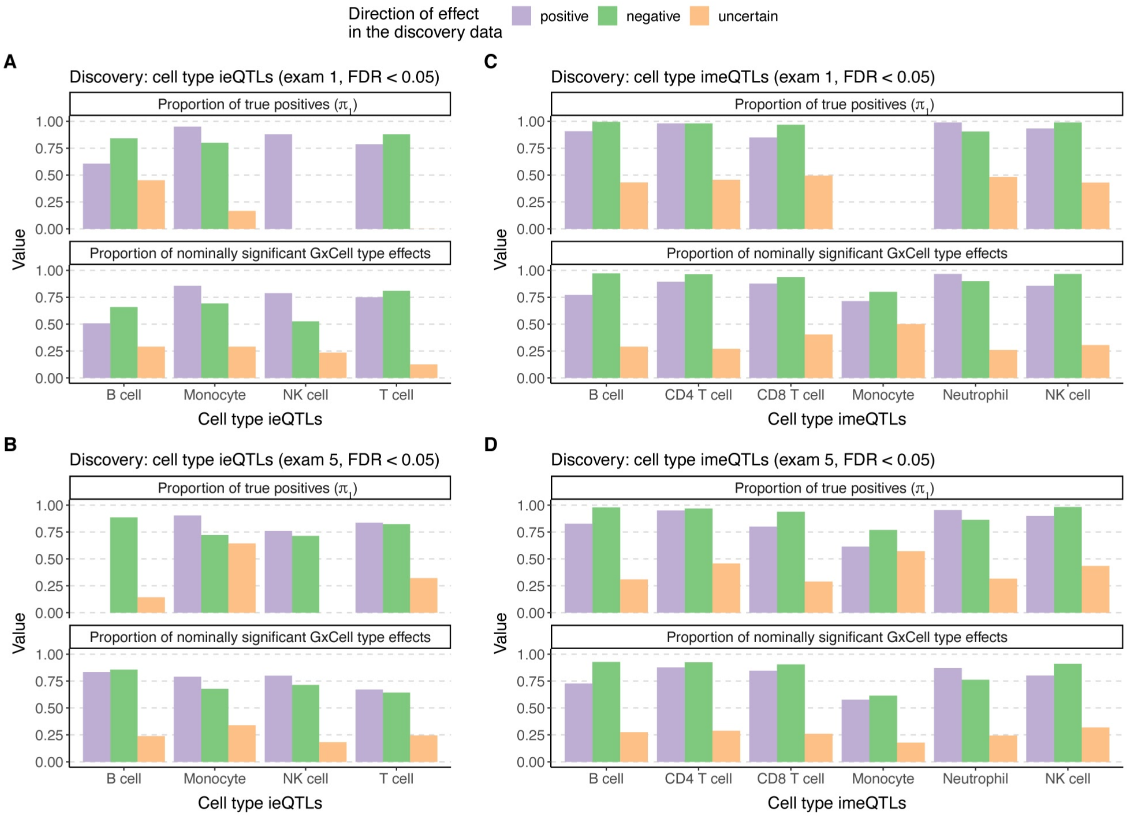


**Figure S7. Reproducibility of cell type iQTLs.** Reproducibility of cell type iQTLs by direction of effect using one of the exams as discovery and the other as validation, and vice versa, for **A)** cell type ieQTLs from exam 1, **B)** cell type ieQTLs from exam 5, **C)** sentinel cell type imeQTLs from exam 5, **D)** sentinel cell type imeQTLs from exam 5. The proportion of true positives ($\boldsymbol{\pi}_1$) and nominally significant GxCell type effects are used as the measure of reproducibility. On average, excluding monocyte imeQTLs, 27% of cell type iQTLs with uncertain direction compared to 81.2% of cell type iQTLs with positive/negative direction showed nominally significant interaction *P*-value in the validation exam. For calculating $\boldsymbol{\pi}_1$, lambda was set to 0.85, if more than 100 ieGenes or imeSites were found from the validation set, or lambda was set to 0.5, if more than 20 ieGenes or imeSites were found from the validation set.

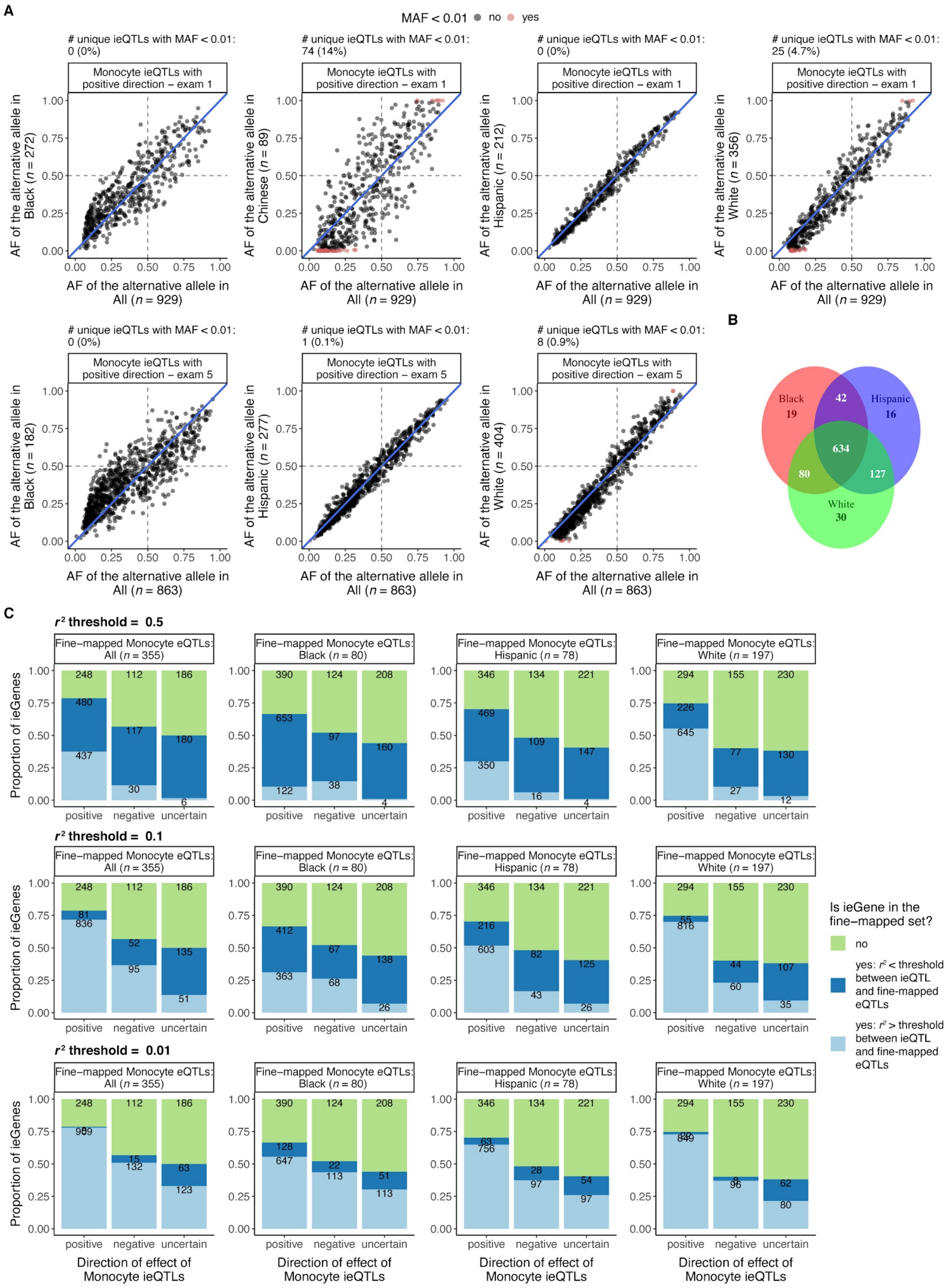

**Figure S8. Population-specificity of cell type iQTLs. A)** Comparison of allele frequencies of the alternative allele calculated from the whole set of individuals used for monocyte ieQTL mapping and a specific group in MESA (based on self-reported race/ethnicity) separately for unique monocyte ieQTLs with positive direction discovered in exam 1 (top row) and exam 1 (bottom row). Note that expression data is available for individuals of Chinese ancestries only in exam 1. **B)** Venn diagram of monocyte ieGenes that have been fine-mapped to likely causal eQTLs in monocytes by self-reported race/ethnicity. **C)** Comparison of monocyte ieQTLs by direction of effect and fine-mapped monocyte eQTLs from MESA using all the individuals with expression data from purified monocytes or by self-reported race/ethnicity. For each gene with a monocyte ieQTL (ieGene) with FDR < 0.05 in exam 1 or exam 5, we assessed whether 1) the ieGene has been fine-mapped to putative causal variants in monocytes, 2) if yes, then whether the maximum LD between the lead ieQTL and fine-mapped monocyte eQTLs from 95% credible set is above or below a specified threshold.

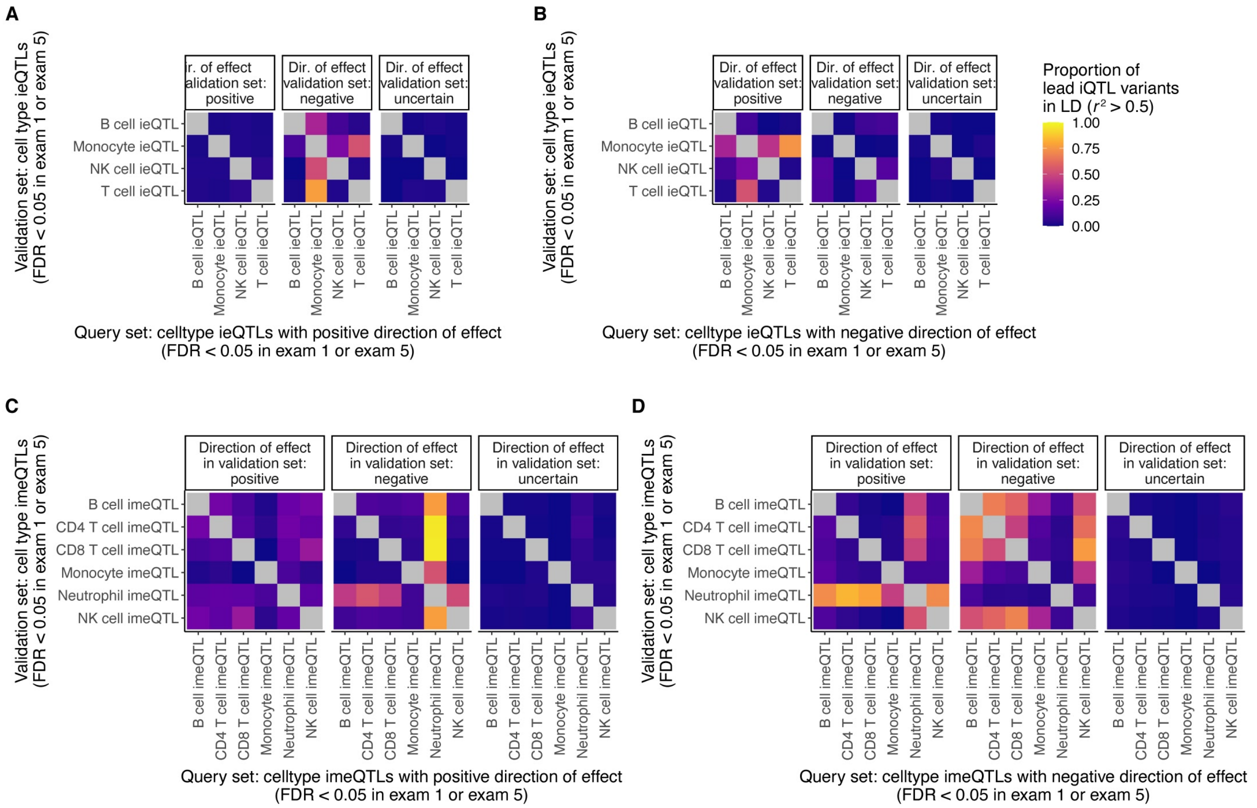


**Figure S9. Pairwise sharing of cell type iQTLs.** Sharing of **A)** cell type ieQTLs with positive direction, **B)** cell type ieQTLs with negative direction, **C)** sentinel cell type imeQTLs with positive direction, and **D)** sentinel cell type imeQTLs with negative direction among the same type of i-molQTLs quantified as the proportion of lead cell type iVariants in LD ($r^2$ > 0.5). To calculate the proportion of iQTLs in LD, the denominator is set to the minimum of the number of iQTLs in the query and validation set.

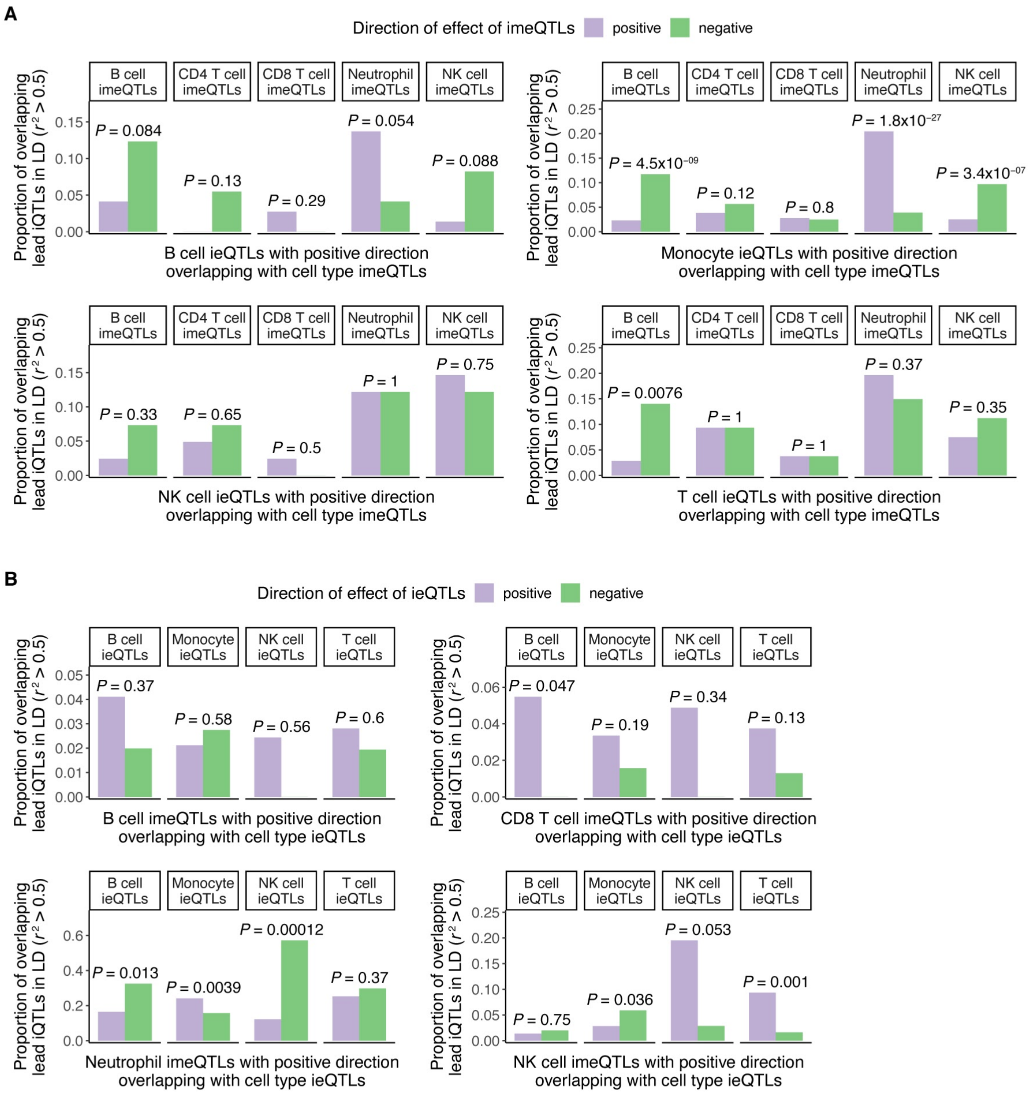


**Figure S10. Sharing between cell type ieQTLs and cell type imeQTLs. A)** Sharing between cell type ieQTLs with positive direction and sentinel cell type imeQTLs (FDR < 0.05 in exam 1 or exam 5) and **B)** sentinel cell type imeQTLs with positive direction and cell type ieQTLs (FDR < 0.05 in exam 1 or exam 5) quantified as the proportion of lead cell type iQTLs in LD ($r^2 > 0.5$). *P*-value shows, for example, the significance of the odds for Monocyte ieQTLs with positive direction to overlap with Neutrophil imeQTLs with positive direction than for Monocyte ieQTL with negative direction. To estimate the odds ratio if any cell is equal to zero in the 2x2 table, Haldane-Anscombe correction was used by adding a fixed value of 0.5 to all cells.

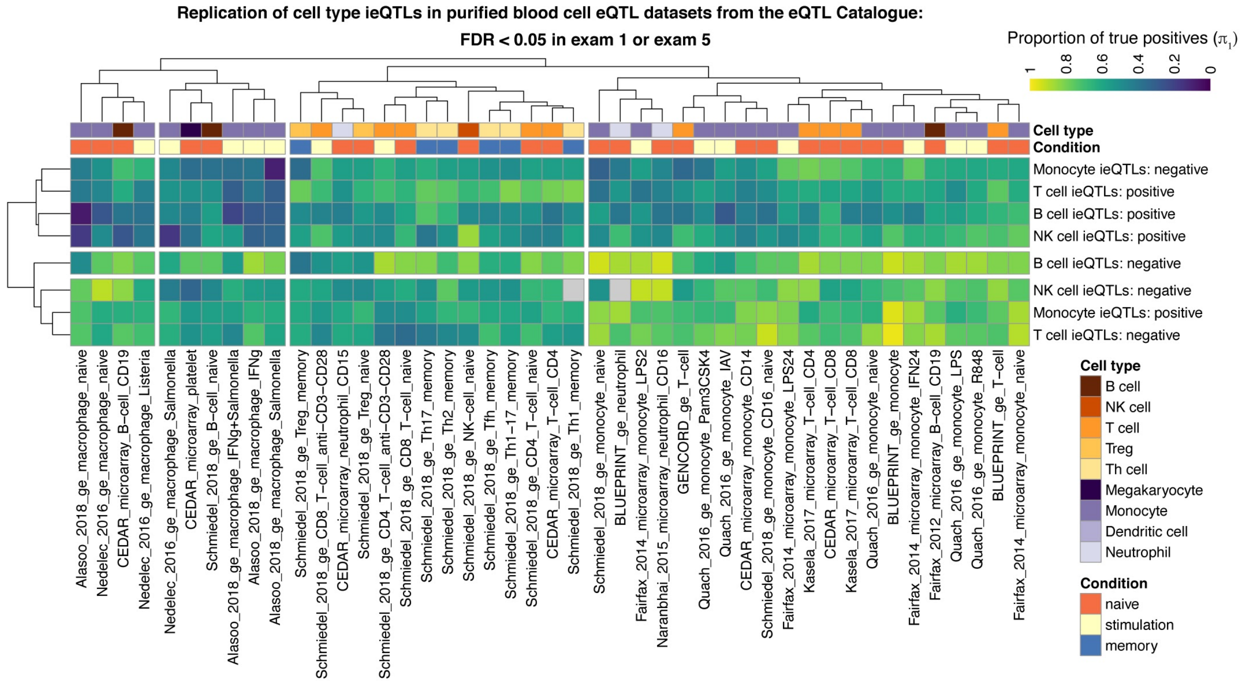


**Figure S11. Replication of cell type ieQTLs in the eQTL Catalogue.** Replication of cell type ieQTLs with FDR < 0.05 in exam 1 or exam 5 data using the eQTL data from various blood cell types in the eQTL Catalogue, 45 eQTL datasets in total. Replication is quantified as the proportion of true positives ($\pi_1$). Clustering of the rows and columns is based on the Euclidean distance and using the complete-linkage method.

**A** Cell type ieQTLs (FDR < 0.05 in exam 1 or exam 5)

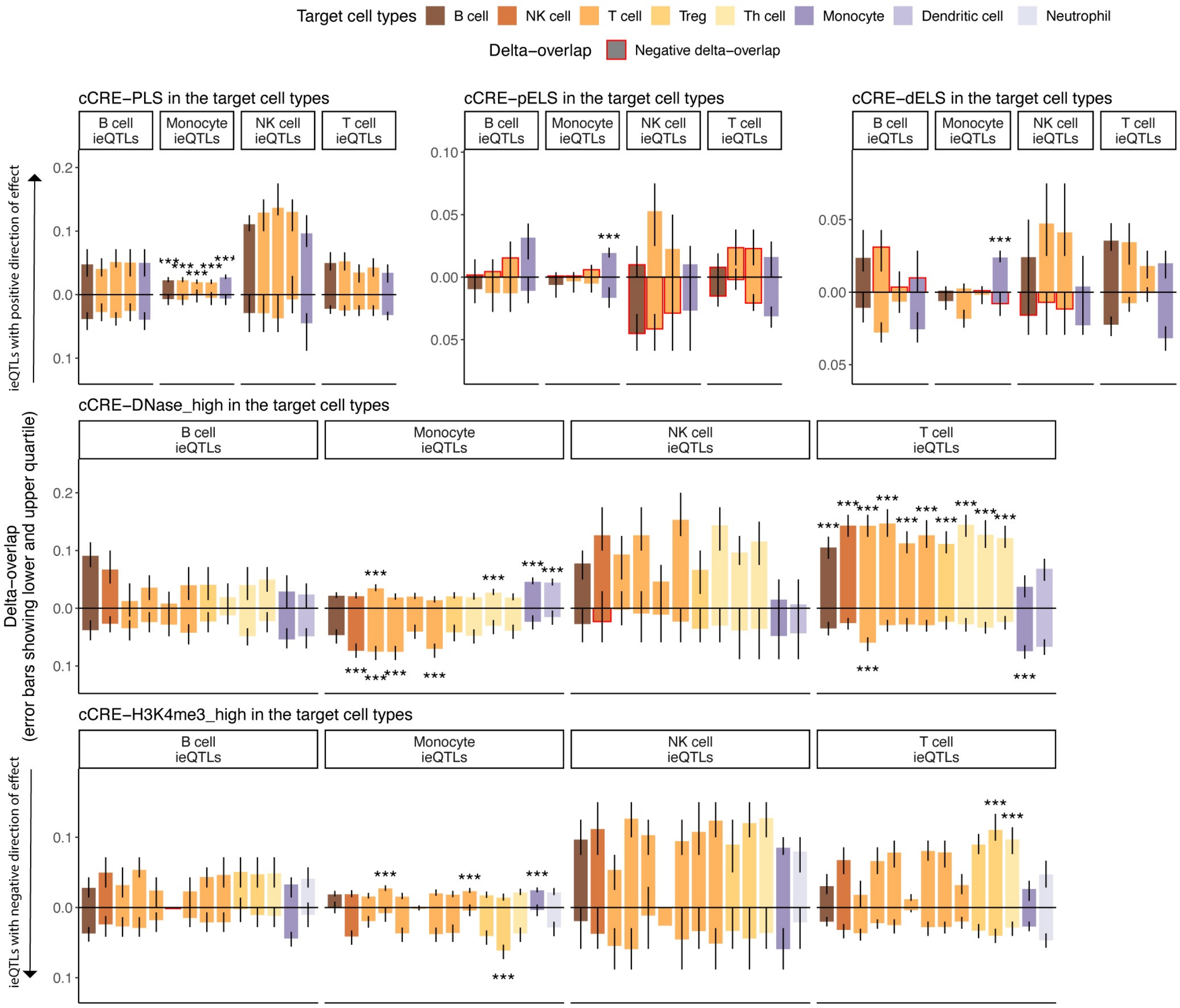

**B** Cell type imeQTLs (FDR < 0.05 in exam 1 or exam 5)

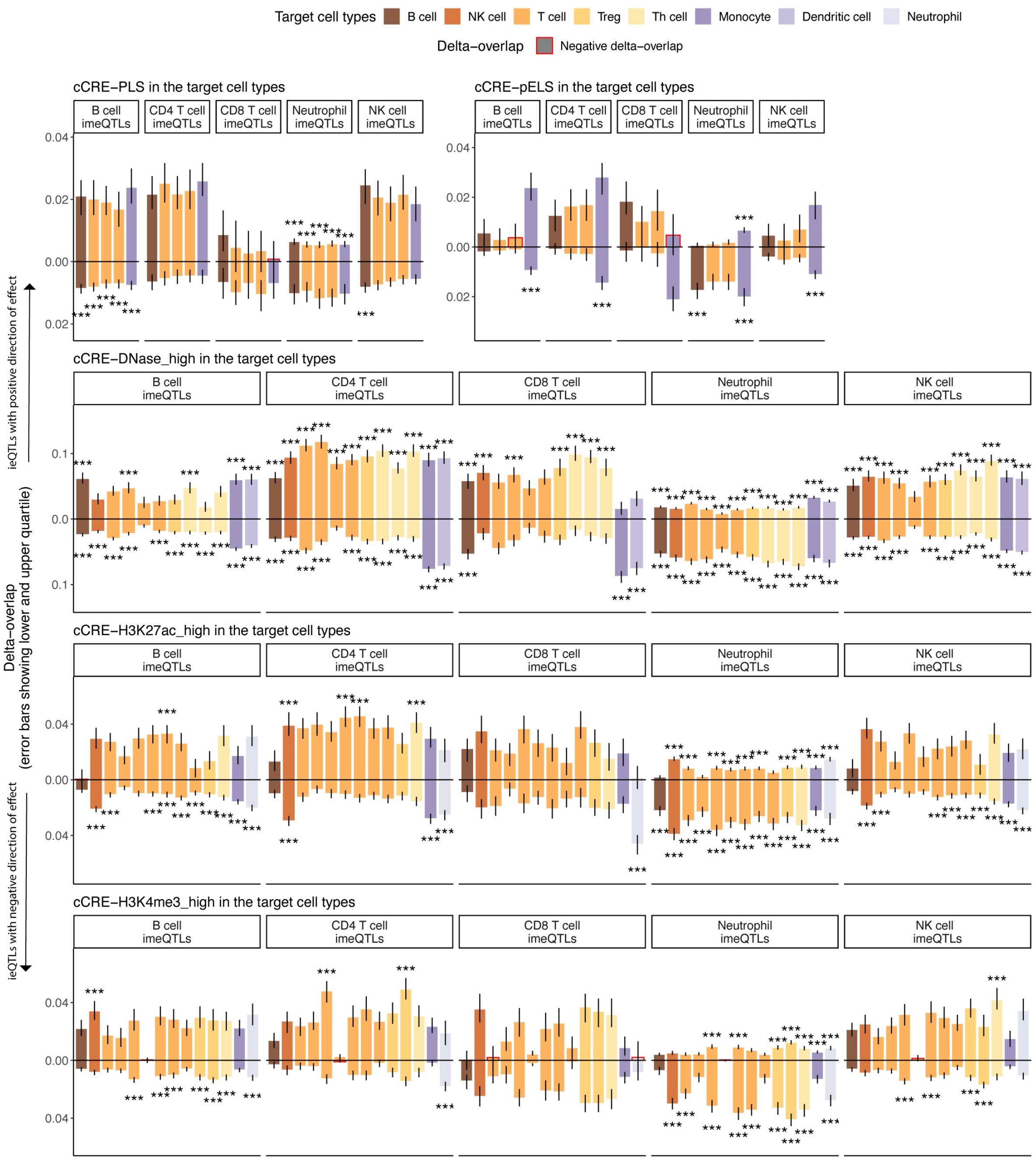


**Figure S12. Functional enrichment analysis of cell type iQTLs**. Functional enrichment analysis with GoShifter showing the delta-overlap, which is the difference between the observed proportion of loci overlapping a cCRE and the mean of the proportion of loci overlapping the cCRE under the null derived by local shifting, for **A**) cell type ieQTLs overlapping different cCRE from ENCODE and **B**) sentinel cell type imeQTLs overlapping different cCRE from ENCODE. Negative delta-overlap indicates cases where observed proportion of loci overlapping an annotation was less than under the null. *** - significant association after correcting for the number of target cell types with cCRE data, number of cell types tested for interaction effect, and the number of groups of direction of effect, applied separately for cell type ieQTLs and cell type imeQTLs.

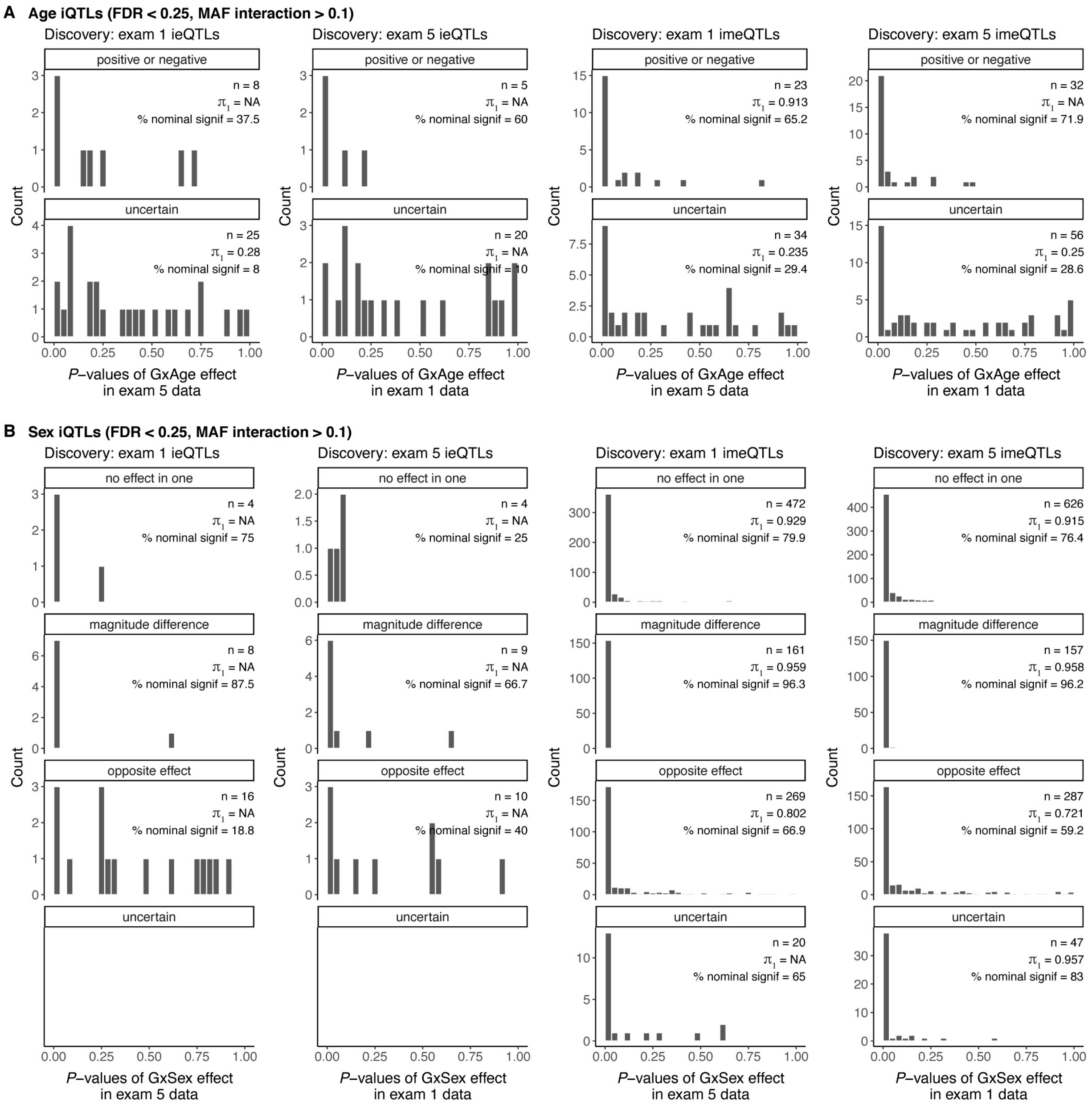
A Age iQTLs (FDR < 0.25, MAF interaction > 0.1)
Discovery: exam 1 ieQTLs
Discovery: exam 5 ieQTLs
Discovery: exam 1 imeQTLs
Discovery: exam 5 imeQTLs
positive or negative
n = 8
π1 = NA
% nominal signif = 37.5
n = 5
π1 = NA
% nominal signif = 60
n = 23
π1 = 0.913
% nominal signif = 65.2
n = 32
π1 = NA
% nominal signif = 71.9
uncertain
n = 25
π1 = 0.28
% nominal signif = 8
n = 20
π1 = NA
% nominal signif = 10
n = 34
π1 = 0.235
% nominal signif = 29.4
n = 56
π1 = 0.25
% nominal signif = 28.6
Count
P–values of GxAge effect in exam 5 data
P–values of GxAge effect in exam 1 data
B Sex iQTLs (FDR < 0.25, MAF interaction > 0.1)
Discovery: exam 1 ieQTLs
Discovery: exam 5 ieQTLs
Discovery: exam 1 imeQTLs
Discovery: exam 5 imeQTLs
no effect in one
n = 4
π1 = NA
% nominal signif = 75
n = 4
π1 = NA
% nominal signif = 25
n = 472
π1 = 0.929
% nominal signif = 79.9
n = 626
π1 = 0.915
% nominal signif = 76.4
magnitude difference
n = 8
π1 = NA
% nominal signif = 87.5
n = 9
π1 = NA
% nominal signif = 66.7
n = 161
π1 = 0.959
% nominal signif = 96.3
n = 157
π1 = 0.958
% nominal signif = 96.2
opposite effect
n = 16
π1 = NA
% nominal signif = 18.8
n = 10
π1 = NA
% nominal signif = 40
n = 269
π1 = 0.802
% nominal signif = 66.9
n = 287
π1 = 0.721
% nominal signif = 59.2
uncertain
n = 20
π1 = NA
% nominal signif = 65
n = 47
π1 = 0.957
% nominal signif = 83
Count
P–values of GxSex effect in exam 5 data
P–values of GxSex effect in exam 1 data

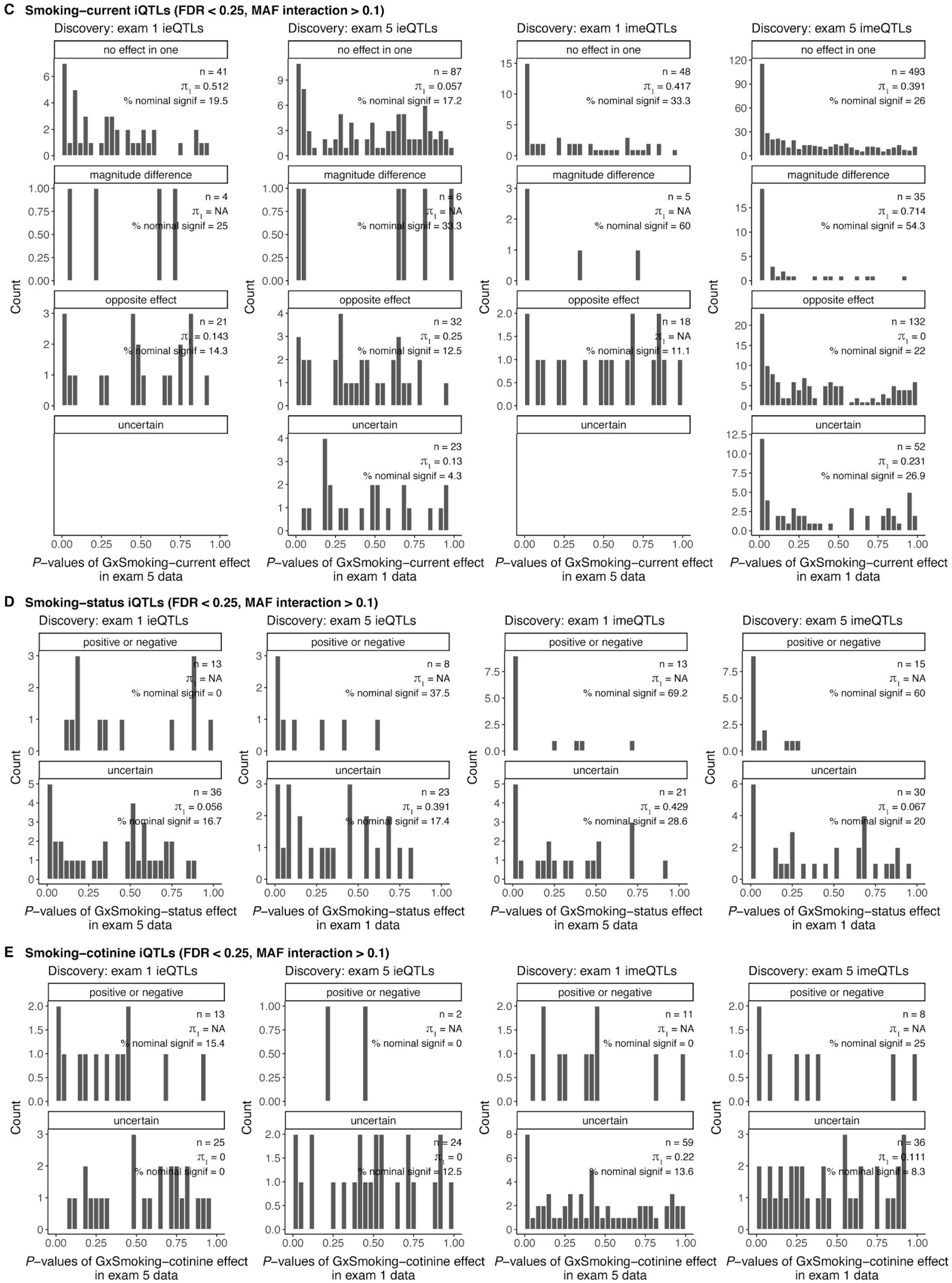


**Figure S13. Reproducibility of age, sex, and smoking.** Reproducibility of trait iQTLs by direction of effect using one of the exams as discovery and the other as validation, and vice versa, for **A)** age ieQTLs and sentinel age imeQTLs, **B)** sex ieQTLs and sentinel sex imeQTLs, **C)** smoking-current ieQTLs and sentinel smoking-current imeQTLs, **D)** smoking-status ieQTLs and sentinel smoking-status imeQTLs, **E)** smoking-cotinine ieQTLs and sentinel smoking-cotinine imeQTLs. The proportion of true positives ($\pi_1$) is used as the measure of reproducibility.

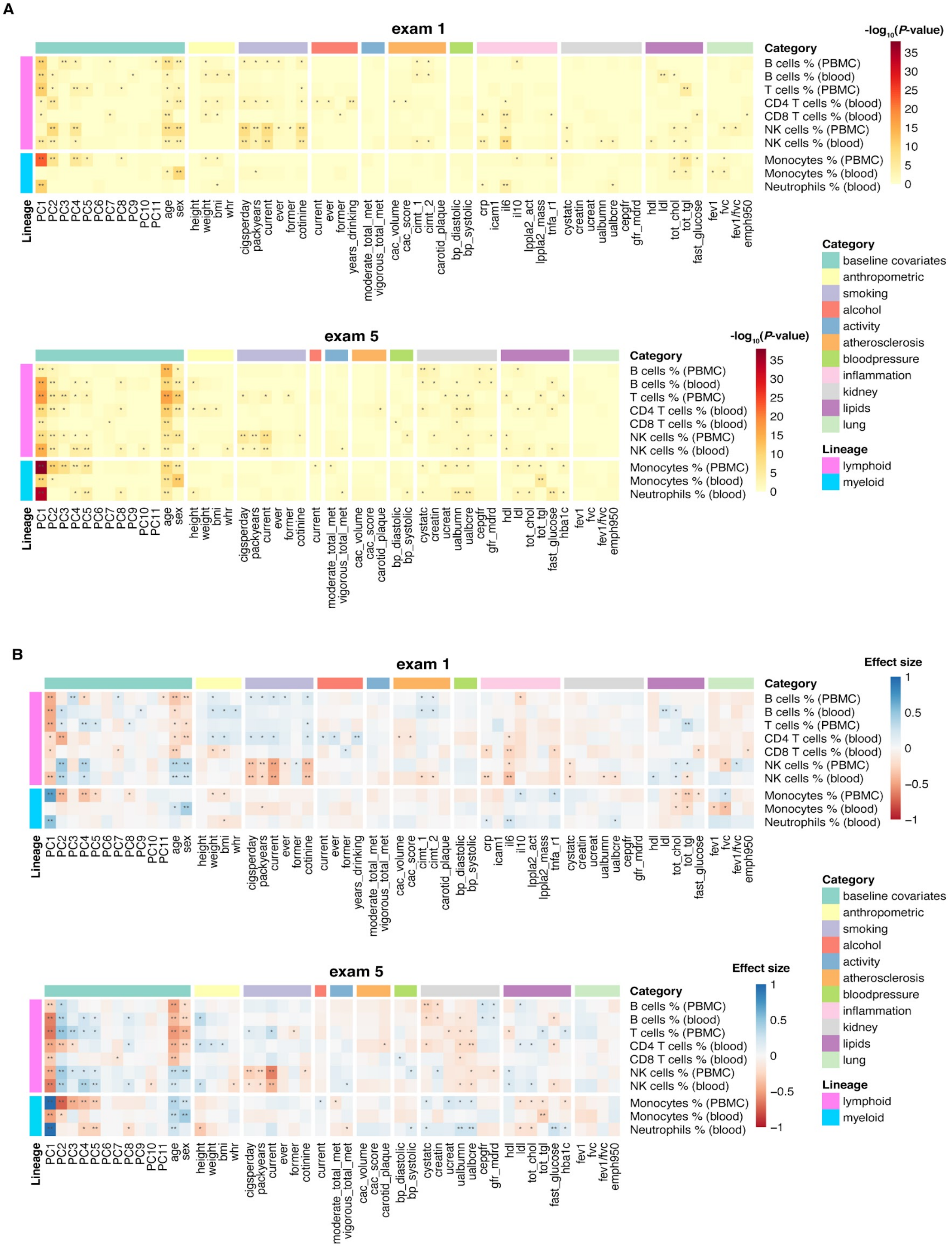


**Figure S14. Association between estimated cell type proportions and various traits in MESA.** Measuring the effect of various traits on estimated cell type proportions in MESA by time point (exam 1 or exam 5) shown as **A**) -log10(*P*-value) and **B**) effect size. The models were adjusted for age, sex, self-reported race/ethnicity, educational attainment, site, and season of the exam. If genotype PCs were the traits of interest, then self-reported race/ethnicity was not included as a covariate. For this analysis, cell type proportions were inverse normalized, and numeric traits were scaled by dividing by two times its standard deviations to make the effect size comparable between untransformed binary traits and numeric traits. ** - denotes Bonferroni significant associations, where P-value < 0.05 / (# of traits groups × # cell type groups), where the count of cell type group is equal to 5 (B cells, T cells/CD4 T cells/CD8 T cells, NK cells, monocytes, neutrophils), * - denotes nominally significant association.

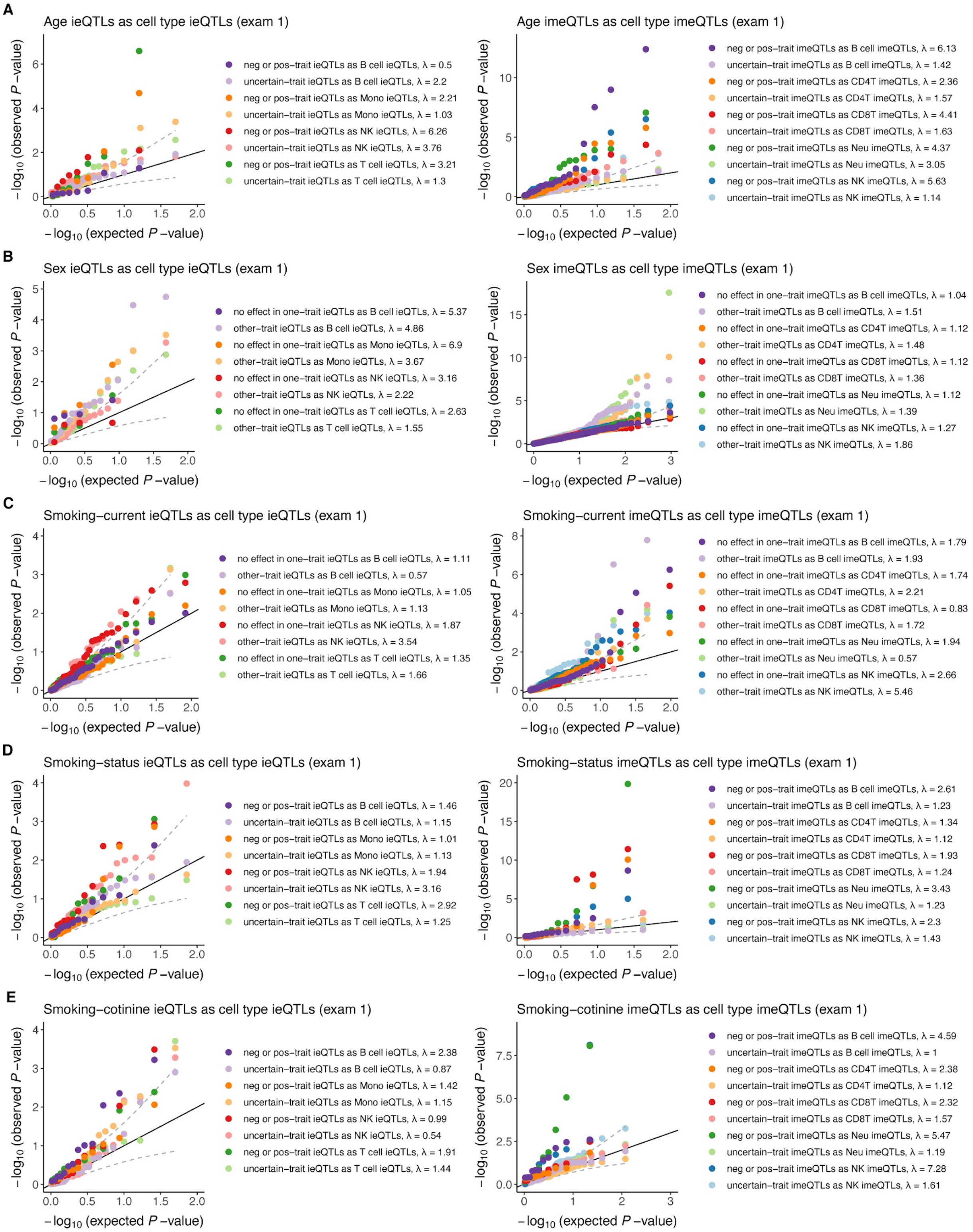

A
Age ieQTLs as cell type ieQTLs (exam 1)
neg or pos−trait ieQTLs as B cell ieQTLs, λ = 0.5
uncertain−trait ieQTLs as B cell ieQTLs, λ = 2.2
neg or pos−trait ieQTLs as Mono ieQTLs, λ = 2.21
uncertain−trait ieQTLs as Mono ieQTLs, λ = 1.03
neg or pos−trait ieQTLs as NK ieQTLs, λ = 6.26
uncertain−trait ieQTLs as NK ieQTLs, λ = 3.76
neg or pos−trait ieQTLs as T cell ieQTLs, λ = 3.21
uncertain−trait ieQTLs as T cell ieQTLs, λ = 1.3
Age imeQTLs as cell type imeQTLs (exam 1)
neg or pos−trait imeQTLs as B cell imeQTLs, λ = 6.13
uncertain−trait imeQTLs as B cell imeQTLs, λ = 1.42
neg or pos−trait imeQTLs as CD4T imeQTLs, λ = 2.36
uncertain−trait imeQTLs as CD4T imeQTLs, λ = 1.57
neg or pos−trait imeQTLs as CD8T imeQTLs, λ = 4.41
uncertain−trait imeQTLs as CD8T imeQTLs, λ = 1.63
neg or pos−trait imeQTLs as Neu imeQTLs, λ = 4.37
uncertain−trait imeQTLs as Neu imeQTLs, λ = 3.05
neg or pos−trait imeQTLs as NK imeQTLs, λ = 5.63
uncertain−trait imeQTLs as NK imeQTLs, λ = 1.14
B
Sex ieQTLs as cell type ieQTLs (exam 1)
no effect in one−trait ieQTLs as B cell ieQTLs, λ = 5.37
other−trait ieQTLs as B cell ieQTLs, λ = 4.86
no effect in one−trait ieQTLs as Mono ieQTLs, λ = 6.9
other−trait ieQTLs as Mono ieQTLs, λ = 3.67
no effect in one−trait ieQTLs as NK ieQTLs, λ = 3.16
other−trait ieQTLs as NK ieQTLs, λ = 2.22
no effect in one−trait ieQTLs as T cell ieQTLs, λ = 2.63
other−trait ieQTLs as T cell ieQTLs, λ = 1.55
Sex imeQTLs as cell type imeQTLs (exam 1)
no effect in one−trait imeQTLs as B cell imeQTLs, λ = 1.04
other−trait imeQTLs as B cell imeQTLs, λ = 1.51
no effect in one−trait imeQTLs as CD4T imeQTLs, λ = 1.12
other−trait imeQTLs as CD4T imeQTLs, λ = 1.48
no effect in one−trait imeQTLs as CD8T imeQTLs, λ = 1.12
other−trait imeQTLs as CD8T imeQTLs, λ = 1.36
no effect in one−trait imeQTLs as Neu imeQTLs, λ = 1.12
other−trait imeQTLs as Neu imeQTLs, λ = 1.39
no effect in one−trait imeQTLs as NK imeQTLs, λ = 1.27
other−trait imeQTLs as NK imeQTLs, λ = 1.86
C
Smoking−current ieQTLs as cell type ieQTLs (exam 1)
no effect in one−trait ieQTLs as B cell ieQTLs, λ = 1.11
other−trait ieQTLs as B cell ieQTLs, λ = 0.57
no effect in one−trait ieQTLs as Mono ieQTLs, λ = 1.05
other−trait ieQTLs as Mono ieQTLs, λ = 1.13
no effect in one−trait ieQTLs as NK ieQTLs, λ = 1.87
other−trait ieQTLs as NK ieQTLs, λ = 3.54
no effect in one−trait ieQTLs as T cell ieQTLs, λ = 1.35
other−trait ieQTLs as T cell ieQTLs, λ = 1.66
Smoking−current imeQTLs as cell type imeQTLs (exam 1)
no effect in one−trait imeQTLs as B cell imeQTLs, λ = 1.79
other−trait imeQTLs as B cell imeQTLs, λ = 1.93
no effect in one−trait imeQTLs as CD4T imeQTLs, λ = 1.74
other−trait imeQTLs as CD4T imeQTLs, λ = 2.21
no effect in one−trait imeQTLs as CD8T imeQTLs, λ = 0.83
other−trait imeQTLs as CD8T imeQTLs, λ = 1.72
no effect in one−trait imeQTLs as Neu imeQTLs, λ = 1.94
other−trait imeQTLs as Neu imeQTLs, λ = 0.57
no effect in one−trait imeQTLs as NK imeQTLs, λ = 2.66
other−trait imeQTLs as NK imeQTLs, λ = 5.46
D
Smoking−status ieQTLs as cell type ieQTLs (exam 1)
neg or pos−trait ieQTLs as B cell ieQTLs, λ = 1.46
uncertain−trait ieQTLs as B cell ieQTLs, λ = 1.15
neg or pos−trait ieQTLs as Mono ieQTLs, λ = 1.01
uncertain−trait ieQTLs as Mono ieQTLs, λ = 1.13
neg or pos−trait ieQTLs as NK ieQTLs, λ = 1.94
uncertain−trait ieQTLs as NK ieQTLs, λ = 3.16
neg or pos−trait ieQTLs as T cell ieQTLs, λ = 2.92
uncertain−trait ieQTLs as T cell ieQTLs, λ = 1.25
Smoking−status imeQTLs as cell type imeQTLs (exam 1)
neg or pos−trait imeQTLs as B cell imeQTLs, λ = 2.61
uncertain−trait imeQTLs as B cell imeQTLs, λ = 1.23
neg or pos−trait imeQTLs as CD4T imeQTLs, λ = 1.34
uncertain−trait imeQTLs as CD4T imeQTLs, λ = 1.12
neg or pos−trait imeQTLs as CD8T imeQTLs, λ = 1.93
uncertain−trait imeQTLs as CD8T imeQTLs, λ = 1.24
neg or pos−trait imeQTLs as Neu imeQTLs, λ = 3.43
uncertain−trait imeQTLs as Neu imeQTLs, λ = 1.23
neg or pos−trait imeQTLs as NK imeQTLs, λ = 2.3
uncertain−trait imeQTLs as NK imeQTLs, λ = 1.43
E
Smoking−cotinine ieQTLs as cell type ieQTLs (exam 1)
neg or pos−trait ieQTLs as B cell ieQTLs, λ = 2.38
uncertain−trait ieQTLs as B cell ieQTLs, λ = 0.87
neg or pos−trait ieQTLs as Mono ieQTLs, λ = 1.42
uncertain−trait ieQTLs as Mono ieQTLs, λ = 1.15
neg or pos−trait ieQTLs as NK ieQTLs, λ = 0.99
uncertain−trait ieQTLs as NK ieQTLs, λ = 0.54
neg or pos−trait ieQTLs as T cell ieQTLs, λ = 1.91
uncertain−trait ieQTLs as T cell ieQTLs, λ = 1.44
Smoking−cotinine imeQTLs as cell type imeQTLs (exam 1)
neg or pos−trait imeQTLs as B cell imeQTLs, λ = 4.59
uncertain−trait imeQTLs as B cell imeQTLs, λ = 1
neg or pos−trait imeQTLs as CD4T imeQTLs, λ = 2.38
uncertain−trait imeQTLs as CD4T imeQTLs, λ = 1.12
neg or pos−trait imeQTLs as CD8T imeQTLs, λ = 2.32
uncertain−trait imeQTLs as CD8T imeQTLs, λ = 1.57
neg or pos−trait imeQTLs as Neu imeQTLs, λ = 5.47
uncertain−trait imeQTLs as Neu imeQTLs, λ = 1.19
neg or pos−trait imeQTLs as NK imeQTLs, λ = 7.28
uncertain−trait imeQTLs as NK imeQTLs, λ = 1.61
−log10 (observed P −value)
−log10 (expected P −value)

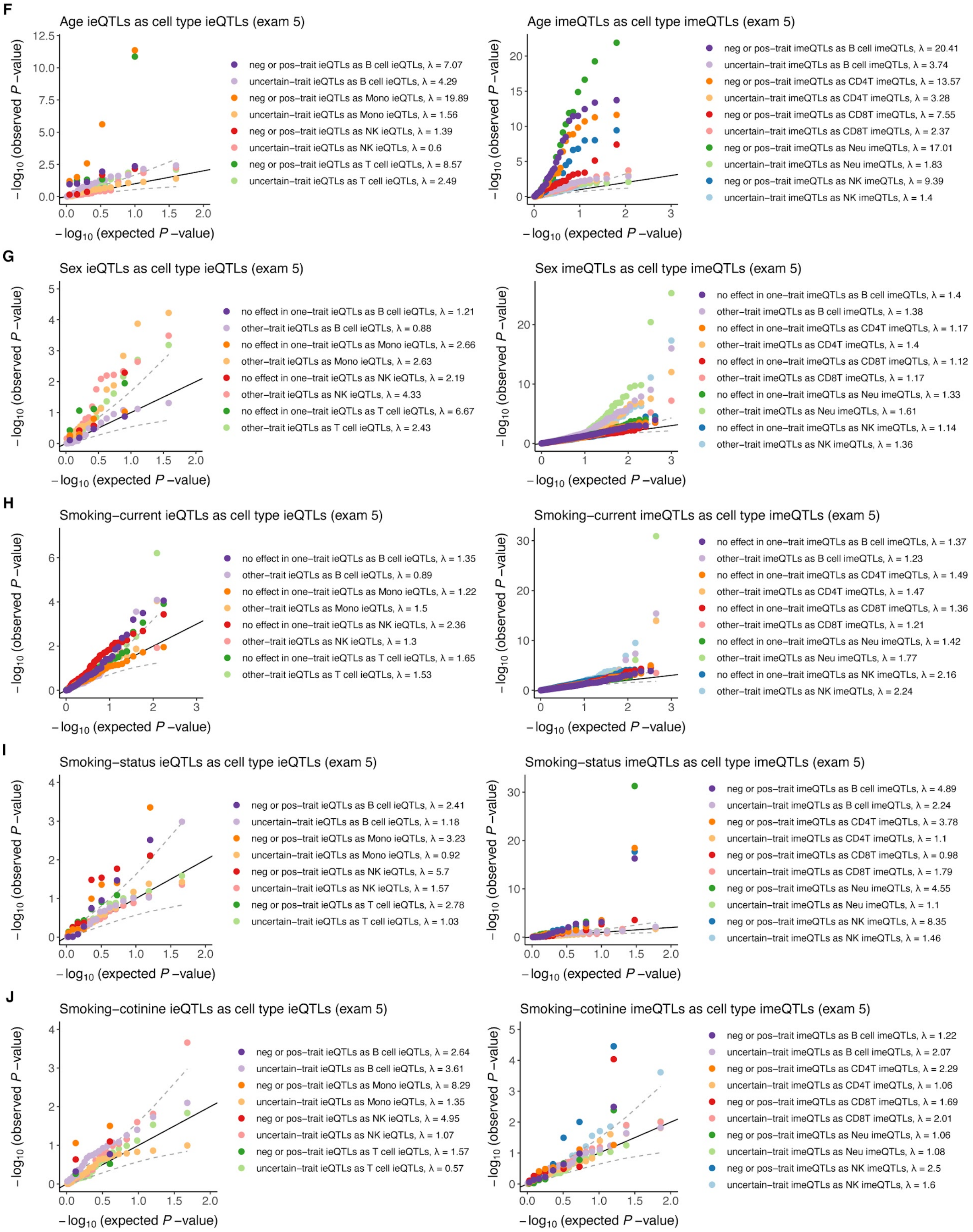


**Figure S15. Enrichment of trait iQTLs as cell type iQTLs.** QQ-plots showing the inflation of GxCell type interaction P-value for **A, F)** age ieQTLs and sentinel age imeQTLs, **B, G)** sex ieQTLs and sentinel sex imeQTLs, **C, H)** smoking-current ieQTLs and sentinel smoking-current imeQTLs, **D, I)** smoking-status ieQTLs and sentinel smoking-status imeQTLs, **E, J)** smoking-cotinine ieQTLs and sentinel smoking-cotinine imeQTLs based on exam 1 and exam 5 data, respectively. Trait iQTLs are classified into two groups based on the direction of effect - positive or negative and uncertain, if trait is continuous; or no effect in one group and other, if trait is binary. Lambda ($\lambda$) is the inflation factor to measure evidence for global inflation of GxCell type effects among trait iQTLs. Median inflation of GxMonocyte effect across sex and smoking ieQTLs (leaving out uncertain direction) was $\lambda = 1.28$ in exam 1 and $\lambda = 2.64$ in exam 5, respectively. Median inflation of GxNeutrophil effect across sentinel sex and smoking imeQTLs (leaving out uncertain direction) was $\lambda = 1.66$ in exam 1 and $\lambda = 1.52$ in exam 5, respectively.

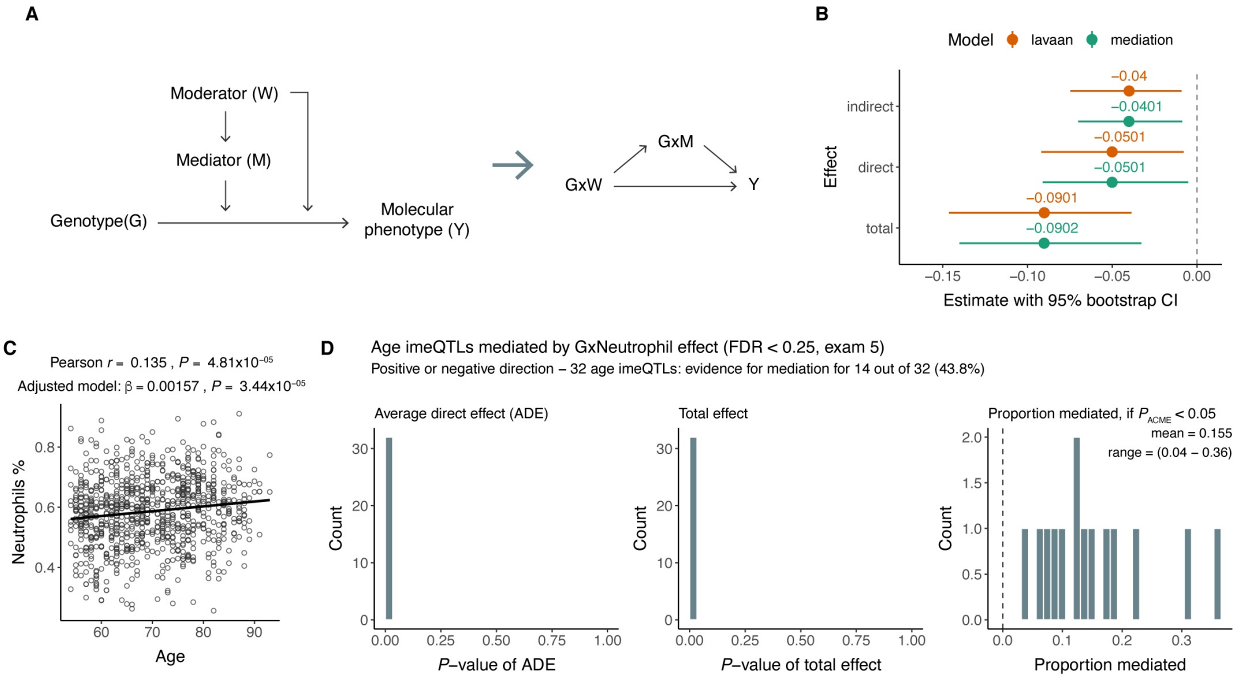


**Figure S16. Mediated moderation model. A)** Conceptual model of mediated moderation, where the mediated moderation effect is described by the path *GxW* → *GxM* → *Y*. **B)** Results of a simulation analysis comparing structural equation model (SEM) for analysis of mediated moderation and mediation analysis approach. We used the lavaan R package for SEM and the mediate R package for mediation analysis. We simulated data for $n$ = 1000 individuals mimicking the moderation effect of age on DNAm mediated by a neutrophil proportion. Genotype vector $G$ was simulated from binomial distribution - G ~ B(2, 0.35), moderator $W$ from normal distribution - $W$ ~ N(0, 9), mediator $M$ as a function of moderator - M ~ 0.0015*$W$ + N(0, 0.1). All the variables were mean-centered, and $W$ was also scaled. Molecular phenotype $Y$ was simulated as a function of $G$, $M$, and $W$ including interactions - $Y$ ~ -0.25*$G$ + 1.5*$M$ - 3*$GxM$ - 0.03*$GxW$ + N(0, 0.5). **C)** The effect of age on estimated neutrophil proportion in MESA exam 5. The adjusted model includes sex, self-reported race/ethnicity, educational attainment, site, and season as covariates. **D)** Results of mediation analysis assessing the mediation of GxAge effect on DNA methylation via GxNeutrophil effect. Barplots show the $P$-value distribution of average direct effect (ADE), total effect, and proportion of mediated effect.

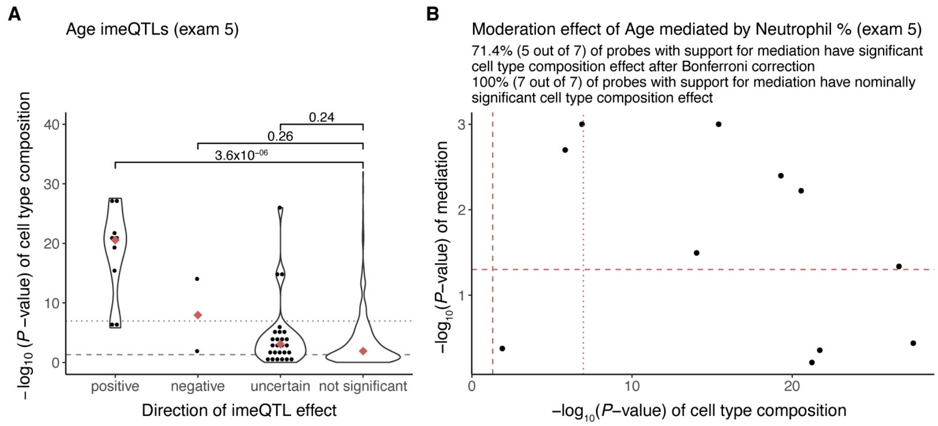


**Figure S17. Age imeQTLs and cell type composition probes. A)** Evidence for age imeSites from exam 5 to be associated with cell type composition by the direction of imeQTL effect. Group "not significant" includes imeQTL with adjusted *P*-value > 0.1. Dashed line denotes nominal significance (*P*-value of cell type composition < 0.05) and dotted line denotes significant association after Bonferroni correction (*P*-value of cell type composition < 0.05/456,655). Only autosomal probes present on the 450K array are included. Using Wilcoxon rank sum test to compare "not significant" group with others. **B)** Comparison of support for the moderation of effect of age to be mediated by neutrophil % and cell type composition effects of probes. Included are age imeQTLs from exam 5 with positive or negative direction that are also present on the 450K array. Dashed line denotes nominal significance (*P*-value of cell type composition < 0.05 or *P*-value of ACME < 0.05) and dotted line denotes significant association after Bonferroni correction (*P*-value of cell type composition < 0.05/456,655).

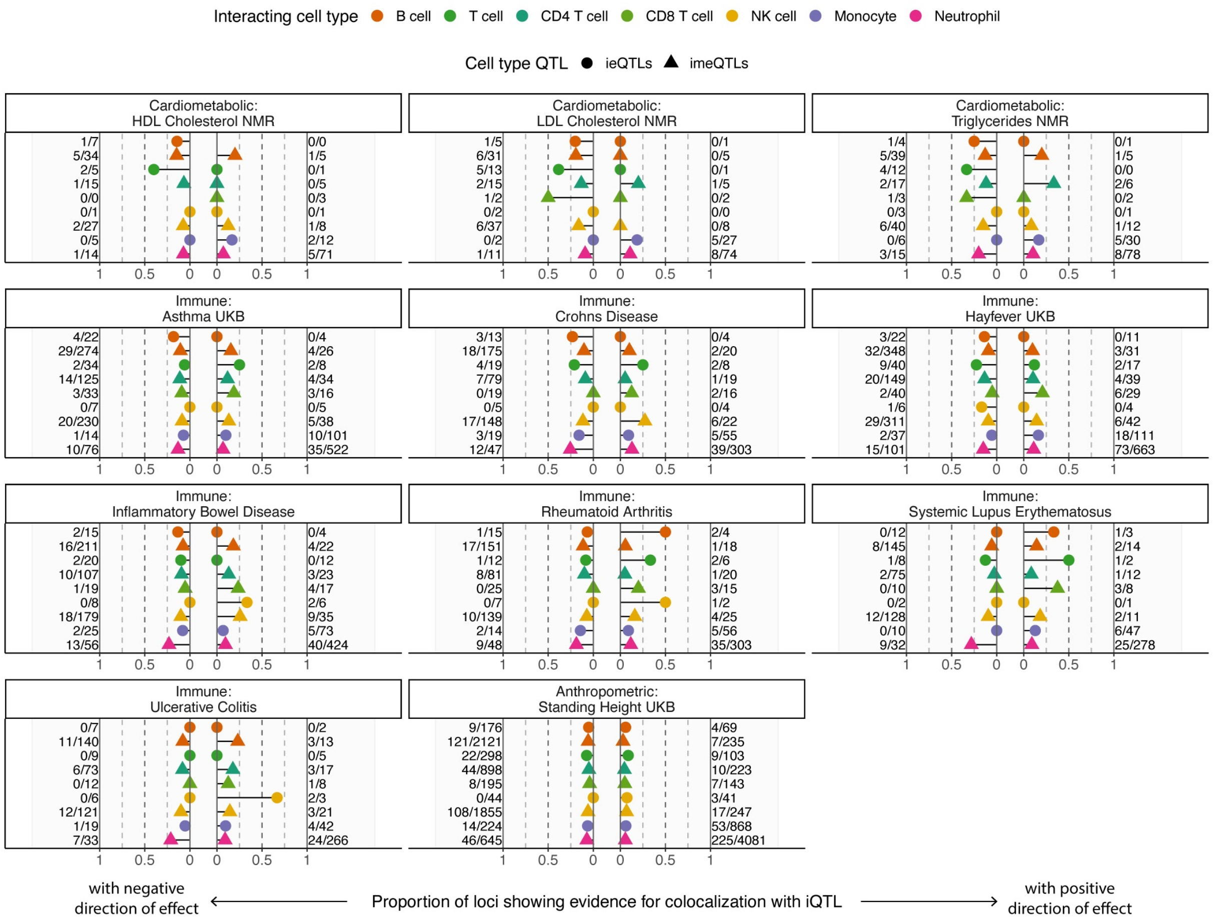


**Figure S18. Colocalization between cell type iQTLs and selected GWAS traits.** Summary of colocalization analysis between cell type iQTLs and selected GWAS traits by the direction of the cell type iQTL effect. We included cell type ieQTLs with FDR < 0.25 in exam 1 or exam 5 and sentinel cell type imeQTLs with FDR < 0.05 in exam 1 or exam 5 to this analysis. Support for colocalization is defined as PP4 > 0.5.

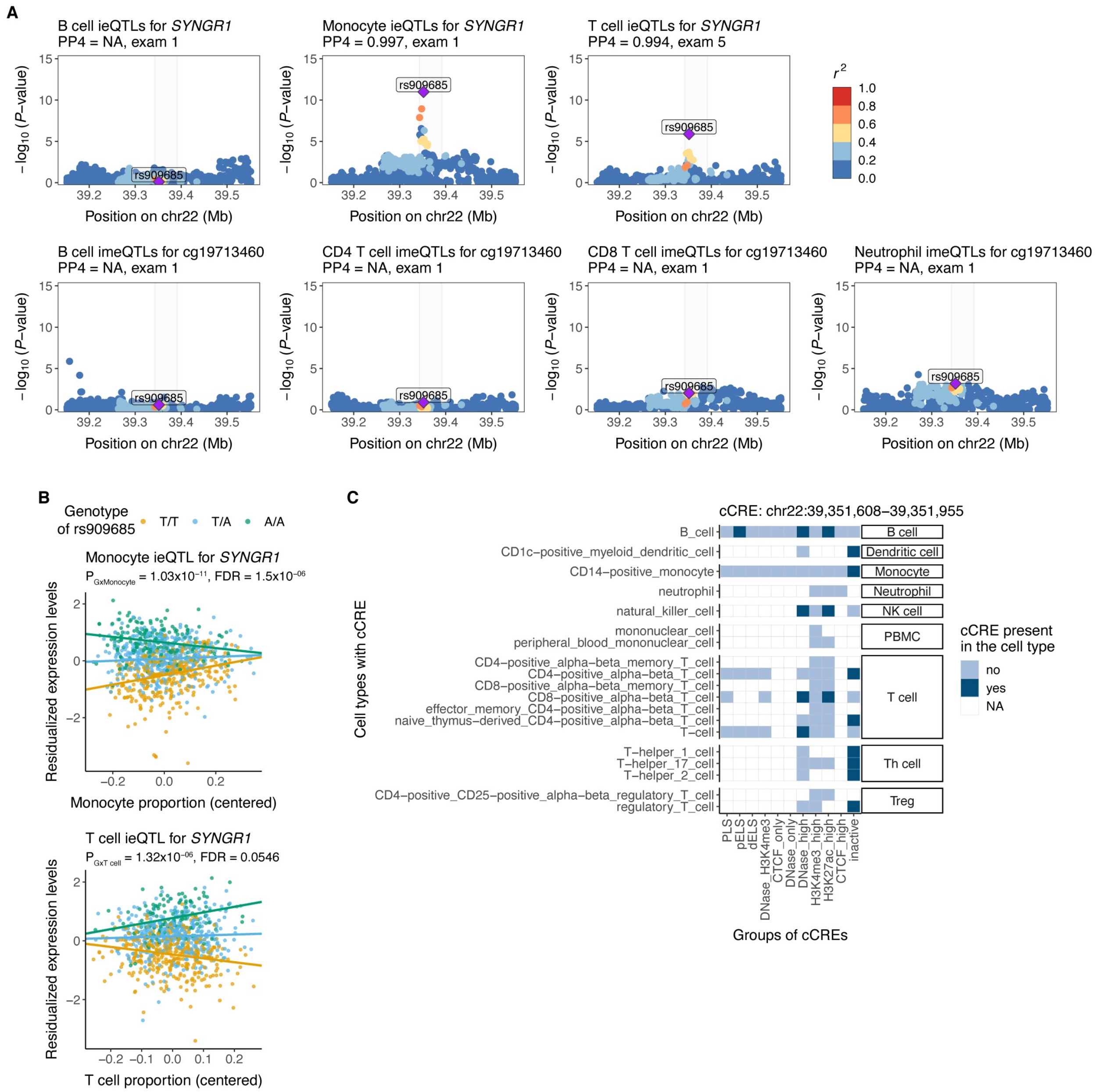


**Figure S19. Cell type iQTLs for *SYNGR1* and association with rheumatoid arthritis. A)** Colocalization between GWAS for rheumatoid arthritis (RA) and cell type iQTLs other than NK cell iQTLs for *SYNGR1* or a nearby CpG site cg19713460 shown as regional association plots. Highlighted region is depicted as in **Fig. 5B** and shows the location of the lead GWAS variant for RA, rs909685, and the CpG site relative to the *SYNGR1* gene. Posterior probability for colocalization (PP4) is set to NA, if the cell type iQTL was not included to the colocalization analysis (FDR < 0.25 for ieQTLs and FDR < 0.05 for imeQTLs). For cell type iQTLs included to the colocalization analysis, showing data from the datapoint with the lowest interaction *P*-value. Otherwise, exam 1 is used, where we observed the association with NK cell iQTLs. **B)** Association plot for monocyte ieQTL and T cell ieQTL for *SYNGR1*. Dots are colored based on the genotype of rs909685. **C**) Overview of the candidate *cis*-regulatory element, where the lead iQTL rs909685 is located. This cCRE is characterized by high DNase and H3K27ac in B cells, NK cells, and CD8 T cells.

### Supplemental Tables

Supplemental tables are provided as separate Excel files.

**Table S1. Cell type iQTLs affecting both expression levels of a gene and DNA methylation levels of a nearby CpG site.** Pairs of significant cell type ieQTLs and sentinel cell type imeQTLs in LD ($r^2 \geq 0.5$ in MESA) with FDR < 0.05 in exam 1 and exam 5 are listed.

**Table S2. Replication of cell type ieQTLs in the eQTL Catalogue.** Based on cell type ieQTLs with FDR < 0.05 in exam 1 or exam 5. Replication is quantified as the proportion of true positives, absolute median effect size in eQTL data, and proportion of variants showing concordance in allelic direction in eQTL effect.

**Table S3**. **Functional enrichment analysis of cell type iQTLs**. Based on cell type ieQTLs and sentinel cell type imeQTLs with FDR < 0.05 in exam 1 or exam 5.

**Table S4**. **Results of the mediation analysis of age imeQTLs by neutrophil imQTL effect based on exam 5 data**.

**Table S5**. **Results of the colocalization analysis between cell type iQTLs and immune or cardiometabolic traits**. We tested for colocalization with (**A**) cell type ieQTLs with FDR < 0.25 in exam 1 or exam 5 and (**B**) sentinel cell type imeQTLs with FDR < 0.05 in exam 1 or exam 5. Support for colocalization is defined as PP4 > 0.5.

### Supplemental Material and Methods

#### Genotypes from Whole Genome Sequencing

DNA samples from MESA participants were sequenced from whole blood to 38x average coverage on Illumina HiSeq X-Ten with 2x151 paired-end reads using a PCR-free library preparation protocol. Sequence data were mapped to the 1000 Genomes GRCh38 human genome reference sequence with decoy contigs using bwa-mem 0.7.15. Duplicates were marked and base call quality scores were recalibrated and binned to 4 bins using bamUtils dedup_LowMem. SNP and short indel variants were discovered and genotyped with vt_discover2. Genotyping used an adjusted genotype likelihood to allow for estimated per-sample DNA sample contamination. After merging genotypes across all TOPMed studies, variant sites were filtered using an SVM classifier to assign "PASS" and "FAIL" labels. The classifier was trained using a positive set of variants that are polymorphic in Illumina Omni 2.5 genotyping of the 2,504 1000 Genomes samples and a negative set of variants that showed multiple discordant genotypes and/or Mendelian inconsistencies in a set of 7,000+ TOPMed duplicate samples and trio nuclear families. Genotyping and variant filtering were performed jointly across all TOPMed studies. The final phased genotype calls for the 1,319 MESA participants included in this study were obtained from the TOPMed freeze 8 genotypes from dbGaP (accession phs001416).

#### Methylation

DNA methylation was quantified in bisulfite-converted genomic DNA from whole blood using the Illumina EPIC array for 1,894 samples (including replicate samples per individual and related individuals). For quality control and normalization, we used the *meffil* R package (Min et al. 2018). First, IDAT files were parsed and probe intensities were dye-bias and background corrected using the *noob* method (Triche et al. 2013). Second, sample quality control was performed and 4 samples were excluded due to: 1) low quality, i.e., samples with >5% of CpGs with detection P-value > 0.01 or methylated/unmethylated ratio outliers; 2) sex mismatches; or 3) genotype mismatches, based on the explicit SNP probes on the array (concordance threshold = 0.8). Third, methylation samples that passed quality filters were normalized using the functional normalization method (Fortin et al. 2014) as implemented in *meffil*. To reduce unwanted technical variation in the data, the top 11 principal components of the control probes were used, which were determined to minimize the residual variance under a 10-fold cross-validation scheme. Fourth, for analyses of the data, we kept the higher-quality sample among replicates, and for a pair of related individuals, we kept the sample of the individual with the most data types in MESA. As a result, the final sample size was n = 900 and n = 899 for exam 1 and exam 5, respectively. Additionally, we performed filtering of CpG sites as follows: 1) probes with >10% of samples with detection P-value > 0.01; 2) masking of probes as recommended in (Zhou et al. 2017), including excluding probes with mapping problems to GRCh38, cross-reactive probes, Type I probes with putative CCS SNPs; 3) non-CpG targeting probes; 4) probes on Y chromosome; and 5) polymorphic probes (i.e., probes with SNVs within 5bp from the 3’-end of the probe (including the extension base for type II probes)) of MAF over 1% across all unrelated individuals in MESA Freeze 8. The final number of CpG sites retained for analyses was n = 740,291 and n = 747,771 for exam 1 and exam 5, respectively, with n = 747,868 CpG sites present in at least one of the exams. Fifth, for meQTL analyses, beta-values of CpG sites with detection P-value > 0.01 were set to missing. Then, missing values were imputed to the mean followed by inverse normal transformation.

#### RNA

PolyA+ mRNA libraries (Illumina TruSeq) were prepared from 1,163 PBMC samples from exam 1 and 966 PBMC samples from exam 5, and sequenced on Illumina HiSeq 4000 with target coverage of >40M reads at two sequencing centers (Broad Institute of MIT and Harvard and the Northwest Genomics Center at the University of Washington). Additionally, RNA-seq was performed on cell-sorted CD19+ monocytes and CD4+ T cells from 401 and 501 exam 5 samples, respectively. The assays were performed and the resulting data processed at the two centers using harmonized protocols and computational pipelines. RNA-seq data were aligned to the human reference genome GRCh38 (excluding ALT, HLA, and decoy contigs) with STAR v2.6.1d (Dobin et al. 2013). To minimize allelic mapping bias in allelic expression and sQTL analyses, the --waspOutputMode option (van de Geijn et al. 2015) was used with a VCF containing the genotypes for each sample. Quality control of the samples was performed following guidelines described in (GTEx Consortium 2020), using metrics computed with RNA-SeQC v2.4.2 (Graubert et al. 2021). Briefly, low-quality samples were identified and removed based on the following alignment metrics: <10 million mapped reads; read mapping rate <0.2; intergenic mapping rate >0.3; base mismatch rate (mismatched bases divided by total aligned bases) >0.01 for read mate 1 or >0.02 for read mate 2; rRNA mapping rate >0.3. Additionally, outlier samples were identified based on gene expression profiles with a correlation-based statistic and sex incompatibility checks, following methods described in (Wright et al. 2014). Among technical replicates (i.e., the same aliquot sequenced multiple times for QC purposes), the sample with the highest number of reads was retained for inclusion in the analysis freeze set. After applying these filters and excluding samples from related participants, 931 and 864 PBMC samples from exams 1 and 5, respectively, and 355 monocyte and 362 T cell samples from exam 5 remained and were used for all downstream analyses. eQTL phenotypes were quantified as previously described (GTEx Consortium 2020).

#### molQTL mapping

molQTLs were mapped using TensorQTL (Taylor-Weiner et al. 2019). For methylation, the *cis*-window was ±500kb from the CpG; for RNA, the *cis*-window was ±1Mb from the gene's TSS. For all molecular phenotypes, the covariates included PEER factors (see below), sex, and the top 11 genotype PCs (Taliun et al. 2021) to account for population structure. For *cis*-molQTLs, 13,294,020 common variants with MAF ≥ 0.01 across all 1,319 samples in the cohort were used for mapping all molecular phenotypes. For *trans*-QTLs, to minimize potential false discoveries at low allele frequencies, an in-sample threshold of MAF ≥ 0.05 was used.

##### PEER factor selection

The optimal number of PEER factors to include was selected for each molecular phenotype based on the number of features identified with at least one molQTL (e.g., eGenes for gene expression), as described previously (GTEx Consortium et al. 2017).

#### molQTL fine-mapping

Fine-mapping of all molQTLs was performed with the TensorQTL (Taylor-Weiner et al. 2019) implementation of SuSiE (Wang et al. 2020), which is functionally equivalent to the R version, with the parameters L=10 and estimate_prior_method='EM' and defaults otherwise. Fine-mapping was performed on individual-level data, with the same set of covariates as used for QTL mapping. For *cis*-QTLs, the *cis*-window was the same as used for QTL mapping; for *trans*-QTLs, a window of ±1Mb around the lead variant was used.

#### Estimation of PBMC cell type proportions from RNA-seq

Cell type proportions were estimated using the R implementation (v1.0.4) of CIBERSORT (Newman et al. 2015) with default settings and the LM22 gene signature matrix provided with the software. CIBERSORT was run on the gene expression matrix (in TPM, from RNA-SeQC) containing the 2,648 analysis freeze samples, which included samples with sorted CD19+ monocytes and CD4+ T cells. For duplicate gene symbols, the gene with the highest mean expression across all samples was retained and others removed from the input. The proportions of cell type subcategories were added up, resulting in proportions for the following broad cell type categories: B cells, Dendritic cells, Eosinophils, Macrophages, Mast cells, Monocytes, NK cells, Neutrophils, Plasma cells, and T cells.

### Supplemental Note

#### Interaction QTLs and confounders

Commonly in GWAS analysis of cell type composition, cell type estimates are adjusted for relevant confounders, and the inverse normalized residuals are used in the association analysis as the response variable (for example, see Vuckovic *et al.*[1]).

We investigated whether for interaction molecular QTL (iQTL) mapping estimated cell type proportion should be adjusted for relevant confounders, using age as an example of a confounder.

##### 1) Age as confounder

Age affects cell type proportion and molecular phenotype.

We fit 3 different models:

- *C ~ Age, Y ~ G + C + GxC* (naive model),
- *C ~ Age, Y ~ G + C + GxC + Age* (full model - corresponds to data generation criteria),
- *C ~ Age, Y ~ G + C-adj + GxC-adj + Age* (adjusted model),

where

- *C-adj* denotes cell type proportions adjusted for the relevant confounders (age in this example), and using the inverse normalized residuals for modeling.

```
# Number of simulations
sim <- 100

# Effect of age on cell type
beta_age <- seq(-0.5, 0.5, 0.25)

# Sample size
n <- 800

# Age - mimicking age range in MESA
age <- runif(n = n, min = 40, max = 60)
age <- round(age)

# Genotype from binomial distribution, MAF = 0.35
geno <- rbinom(n = n, size = 2, prob = 0.35)
```

###### 1.1) Strong iQTL effect

```
## The effect of covariates on the outcome (molecular phenotype):
```

```
##      geno      cell genoxcell       age
##      2.00      1.20      1.50      0.25
```

#### The effect of age on cell type abundance - beta = -0.5

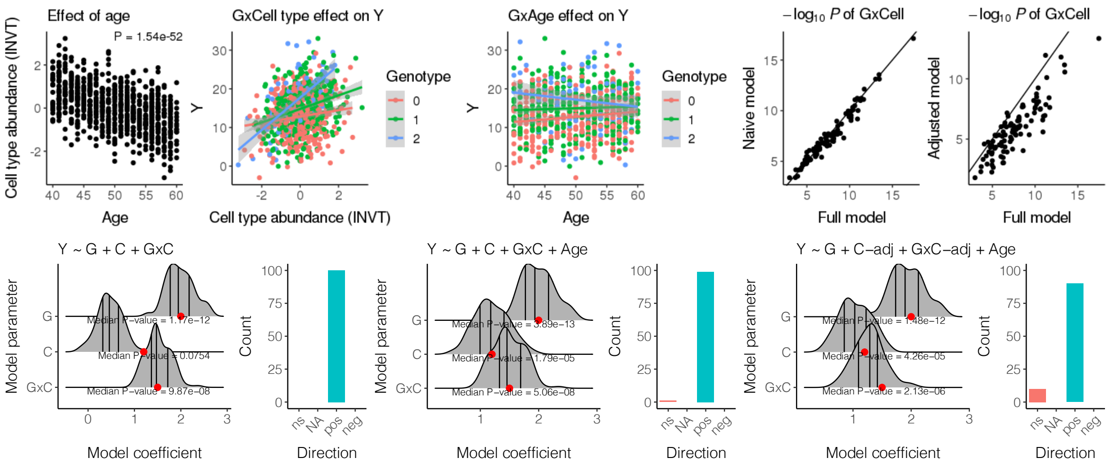


#### The effect of age on cell type abundance - beta = -0.25

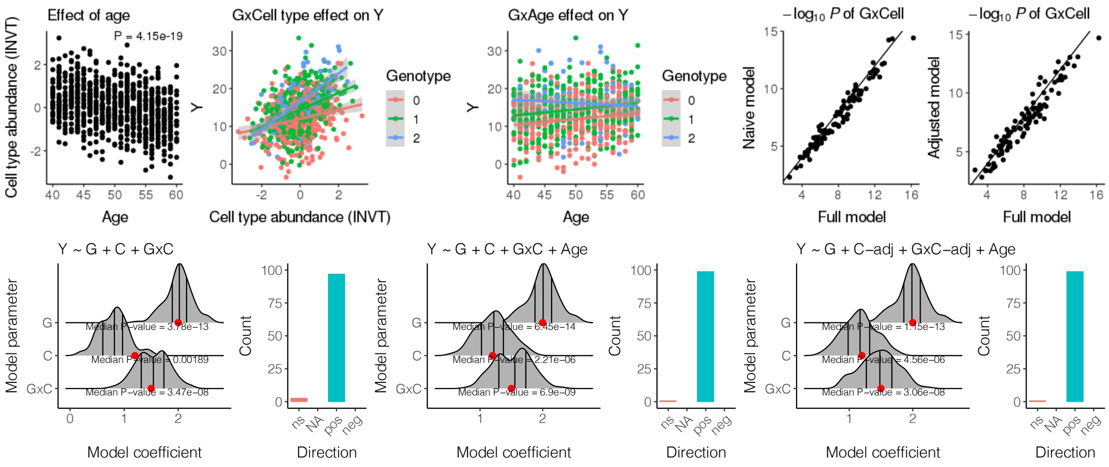

### The effect of age on cell type abundance - beta = 0

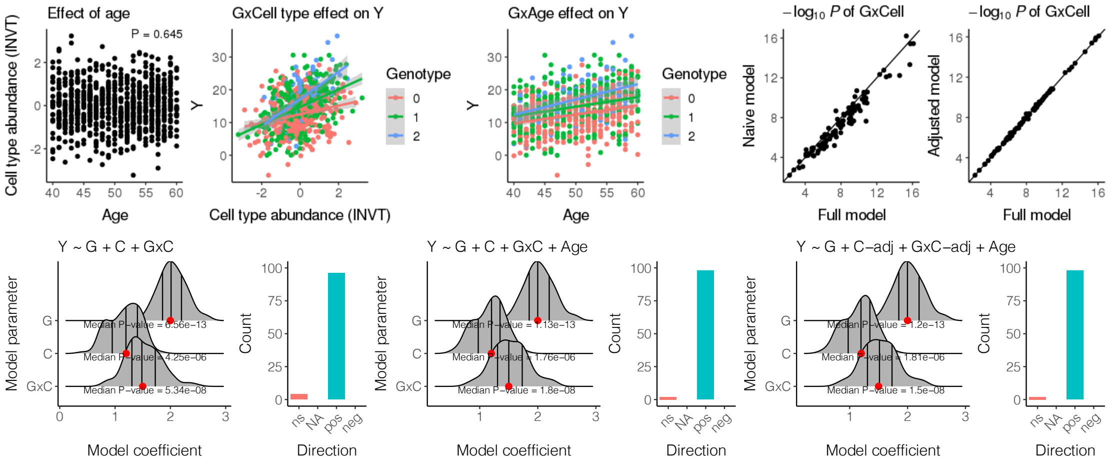


### The effect of age on cell type abundance - beta = 0.25

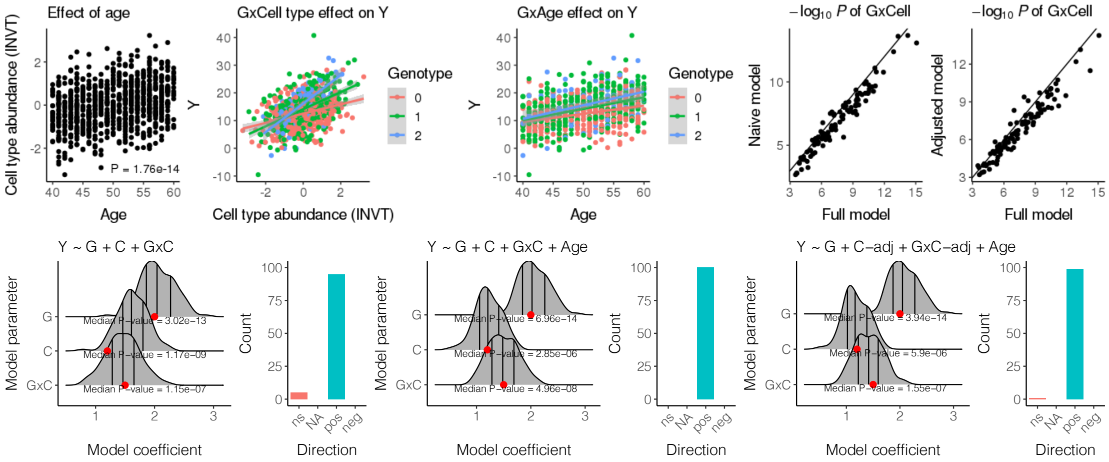


### The effect of age on cell type abundance - beta = 0.5

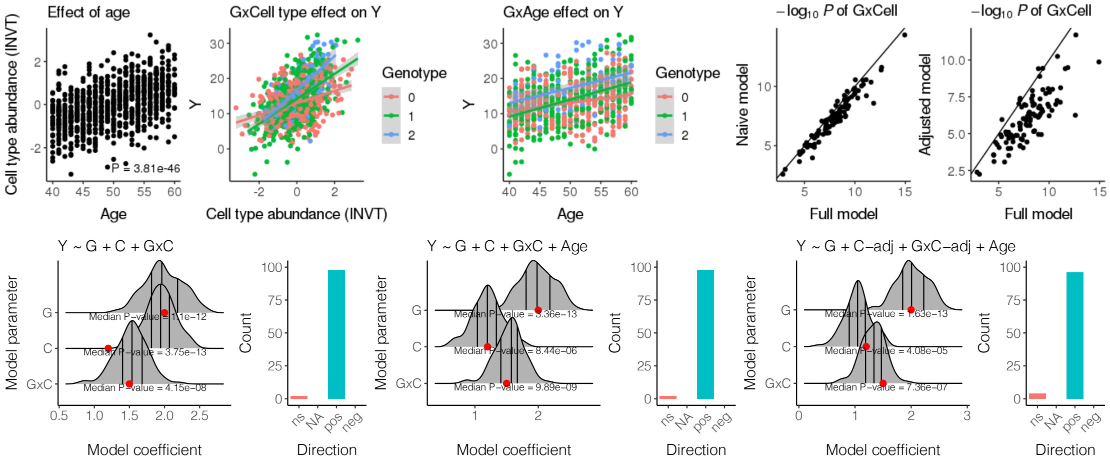

Conclusion

For detecting strong iQTL effects between genotype and cell type abundance, adjusting cell type proportions for a confounder and using inverse normalized residuals in the model (adjusted model) is less powerful as compared to using unadjusted cell type abundance (full model). The extent of loss of power depends on the correlation between the confounder and cell type abundance. Inclusion of the confounder to the iQTL model as a covariate is sufficient when the confounder also has an effect on the molecular phenotype (outcome variable).

#### 1.2) No iQTL effect

Just molQTL effect, but no interaction effect.

```
## The effect of covariates on the outcome (molecular phenotype):

##      geno      cell genoxcell       age
##      2.00      1.20      0.00      0.25
```

The effect of age on cell type abundance - beta = -0.5

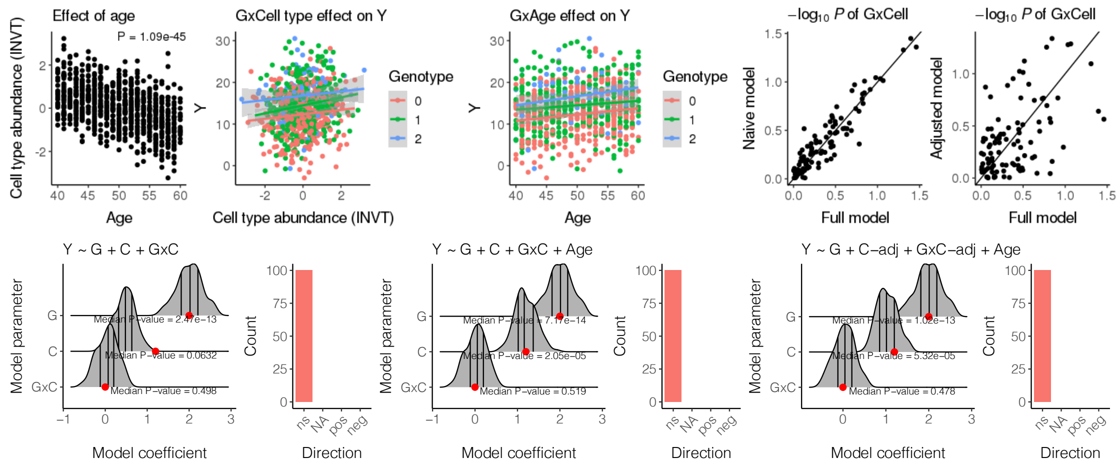

### The effect of age on cell type abundance - beta = -0.25

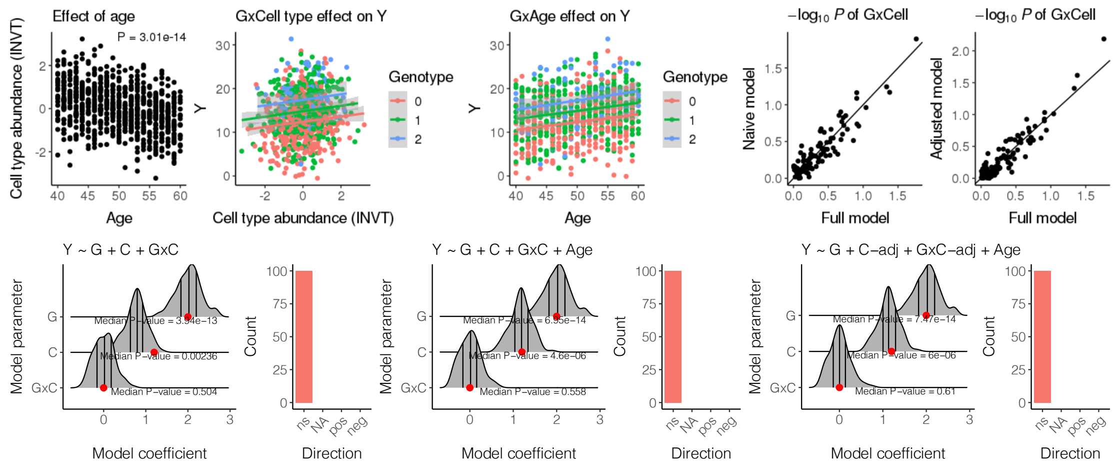


### The effect of age on cell type abundance - beta = 0

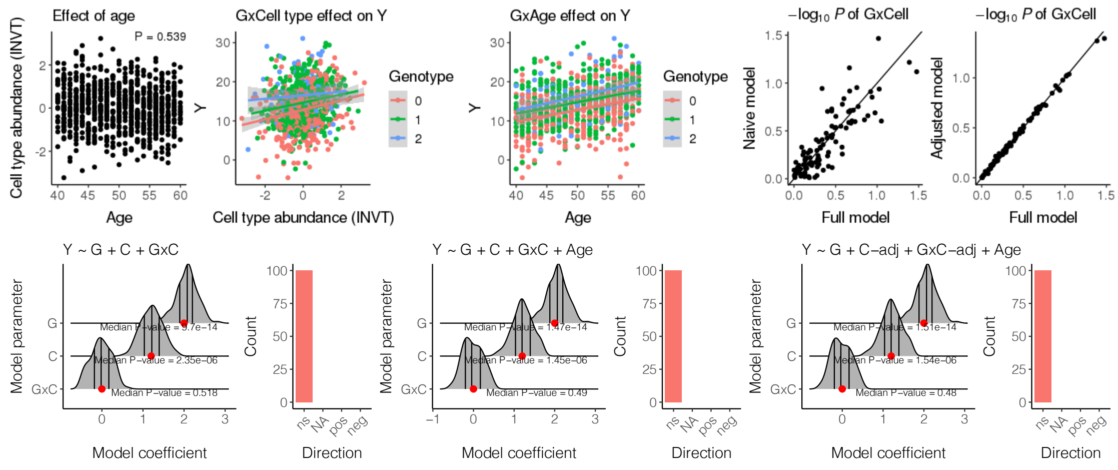


### The effect of age on cell type abundance - beta = 0.25

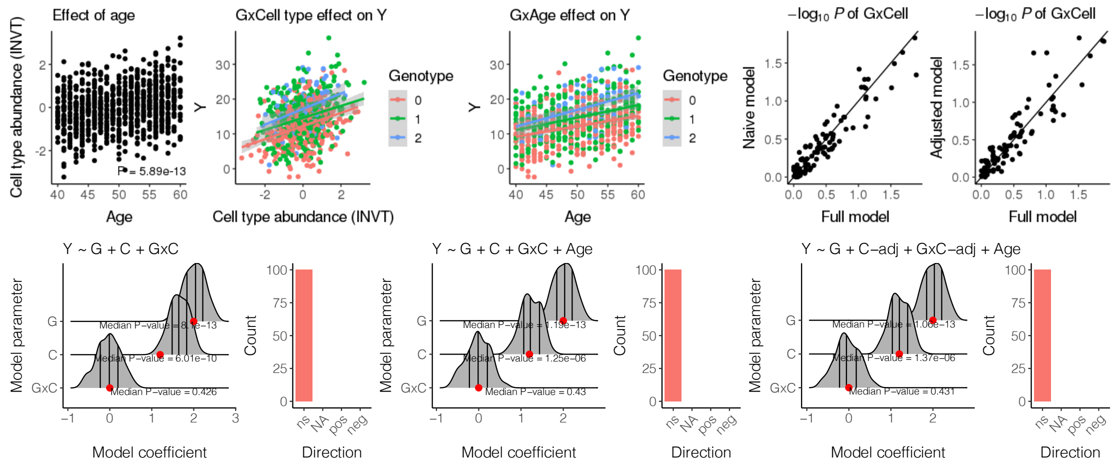

The effect of age on cell type abundance - beta = 0.5

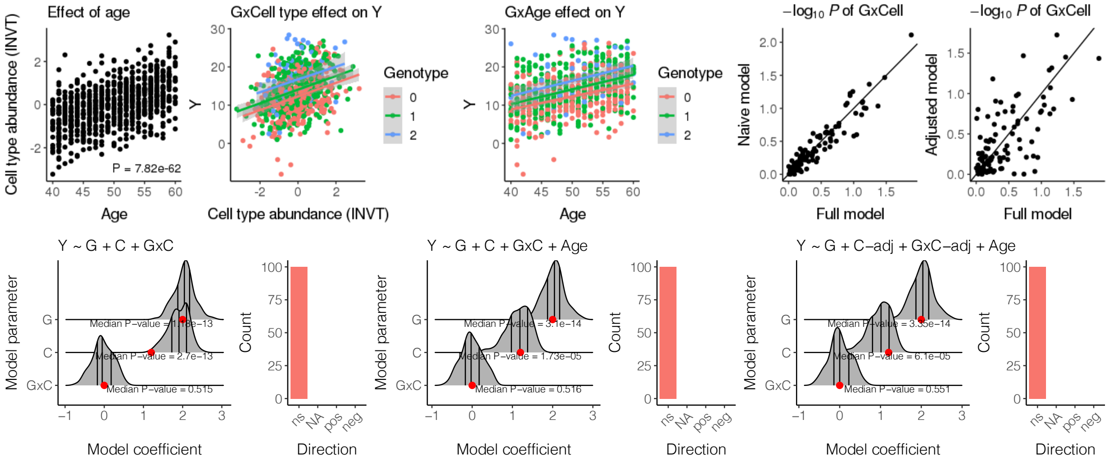


#### Conclusion

If there are no significant iQTL effects, we observe variability between *P*-values from the adjusted model and full model. The extent of variability depends on the correlation between the confounder and cell type abundance. Neither of the models show an increase in false positive rate.

### 2) Age as confounder and moderator

In addition to the effect of age on cell type proportions and molecular phenotype, age also influences the association between G and Y (there is a strong GxAge effect).

We fit 3 different models:

- *C ~ Age, Y ~ G + C + GxC + Age* (naive model),
- *C ~ Age, Y ~ G + C + GxC + Age + GxAge* (full model - corresponds to data generation criteria),
- *C ~ Age, Y ~ G + C-adj + GxC-adj + Age* (adjusted model),

where

- *C-adj* denotes cell type proportions adjusted for the relevant confounders (age in this example), and using the inverse normalized residuals for modeling.

```
# Number of simulations
sim <- 100

# Effect of age on cell type
beta_age <- seq(-0.5, 0.5, 0.25)

# Sample size
```

```r
n <- 800

# Age - mimicking age range in MESA
age <- runif(n = n, min = 40, max = 60)
age <- round(age)

# Genotype from binomial distribution, MAF = 0.35
geno <- rbinom(n = n, size = 2, prob = 0.35)
```

#### 2.1) Strong iQTL effect

```
## The effect of covariates on the outcome (molecular phenotype):

##       geno      cell genoxcell       age  genoxage
##       2.00      1.20      1.50      0.25      0.40
```

##### The effect of age on cell type abundance - beta = -0.5

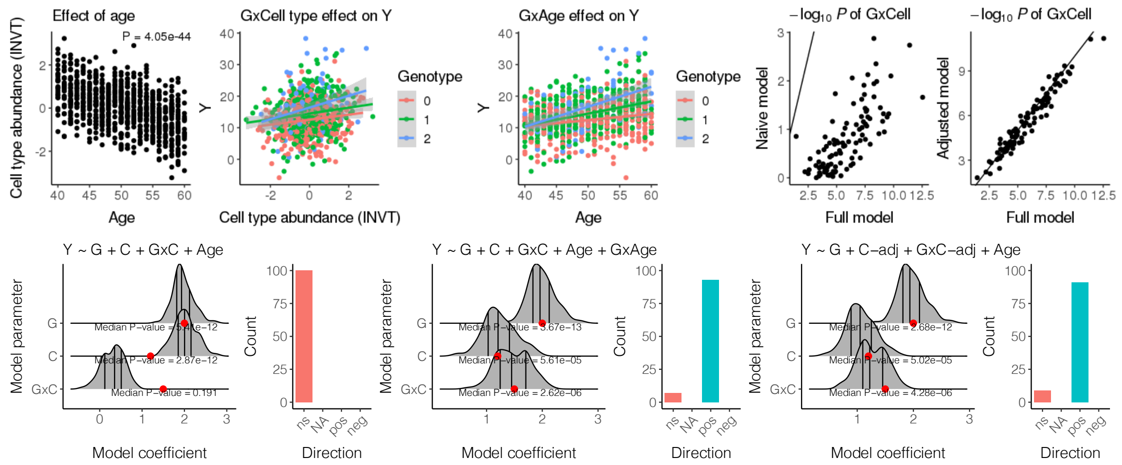


##### The effect of age on cell type abundance - beta = -0.25

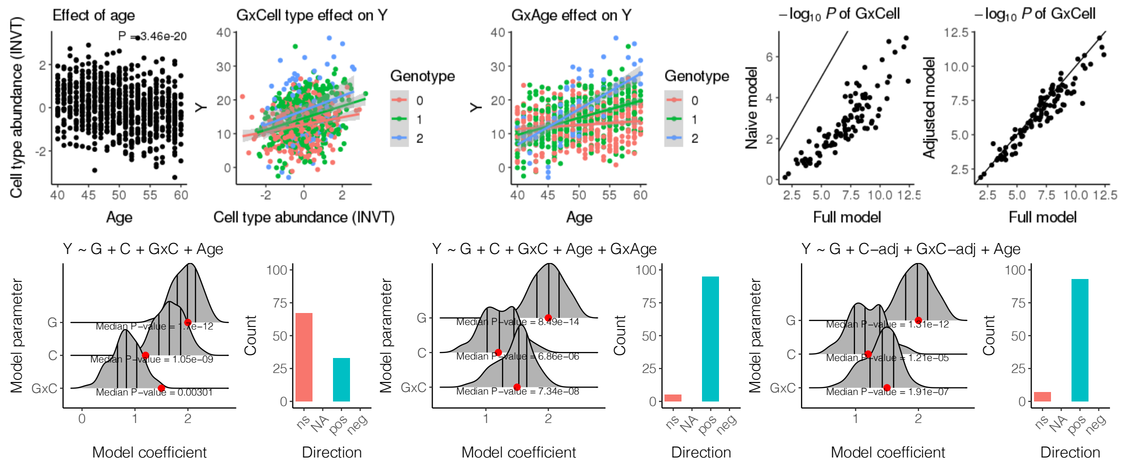

### The effect of age on cell type abundance - beta = 0

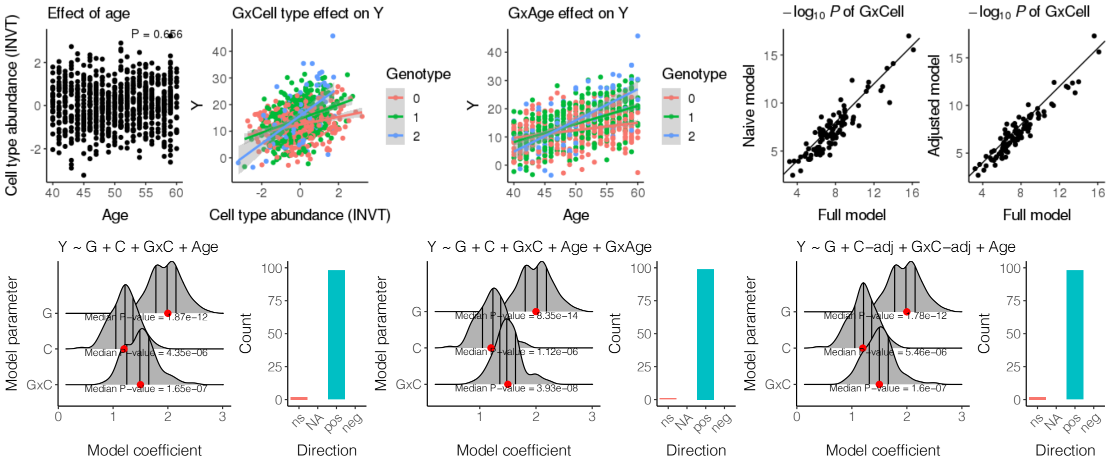


### The effect of age on cell type abundance - beta = 0.25

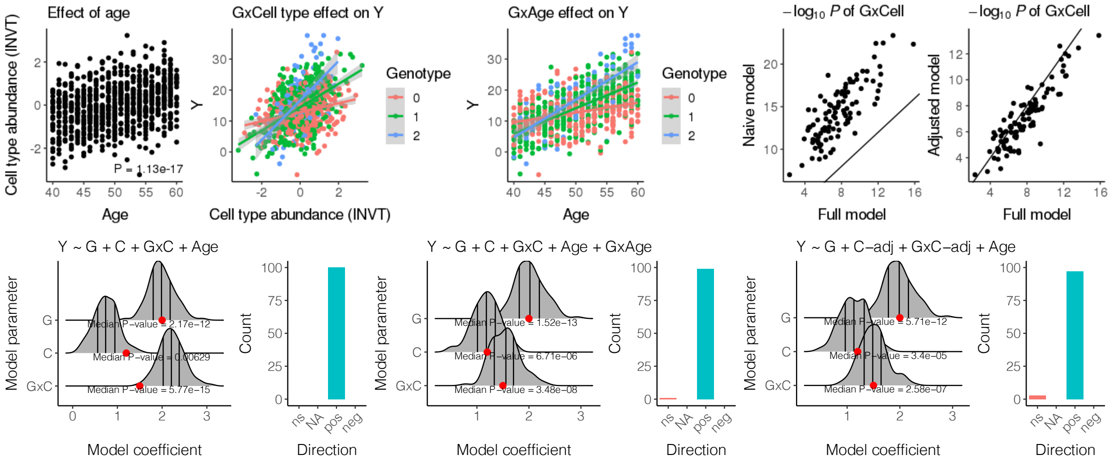


### The effect of age on cell type abundance - beta = 0.5

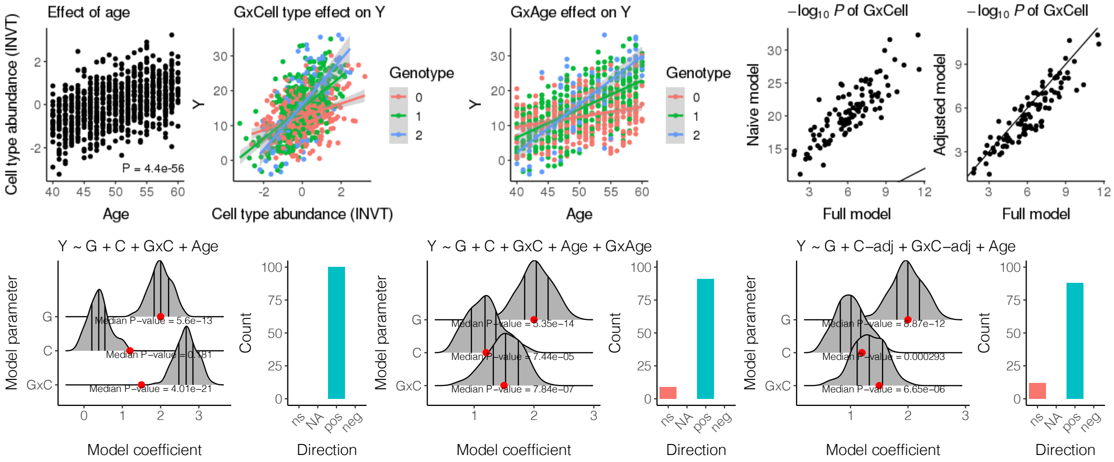

Conclusion

For detecting strong iQTL effects between genotype and cell type abundance in the presence of a strong GxConfounder effect, adjusted model is less powerful compared to full model. However, the current implementation of tensorQTL does not support the inclusion of additional interactions. The naive model, which is currently used for iQTL mapping assuming that relevant confounders are captured by the PEER factors, either overestimates or underestimates the iQTL effect depending on the correlation between the cell type abundance and confounder.

#### 2.2) No iQTL effect

Just molQTL effect, but no interaction effect.

```
## The effect of covariates on the outcome (molecular phenotype):

##      geno      cell  genoxcell       age  genoxage
##      2.00      1.20       0.00      0.25      0.40
```

The effect of age on cell type abundance - beta = -0.5

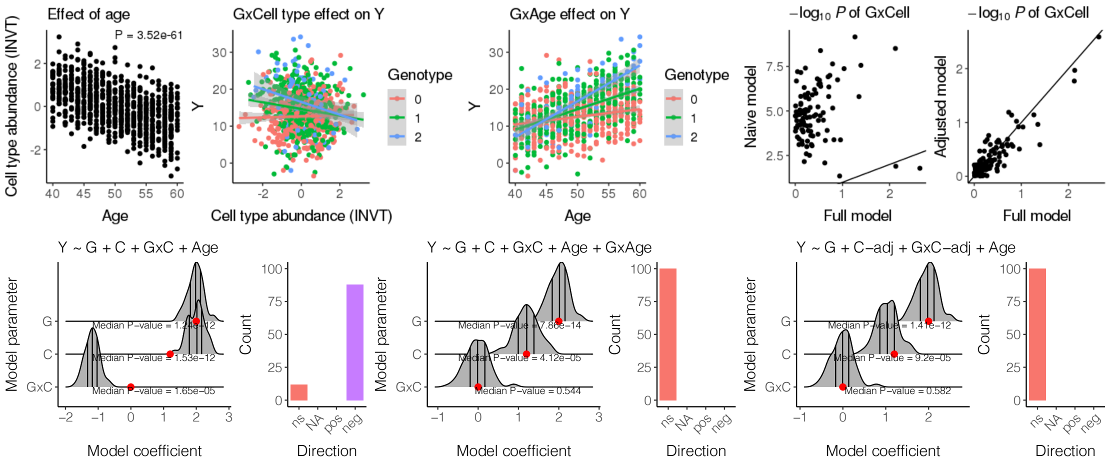

### The effect of age on cell type abundance - beta = -0.25

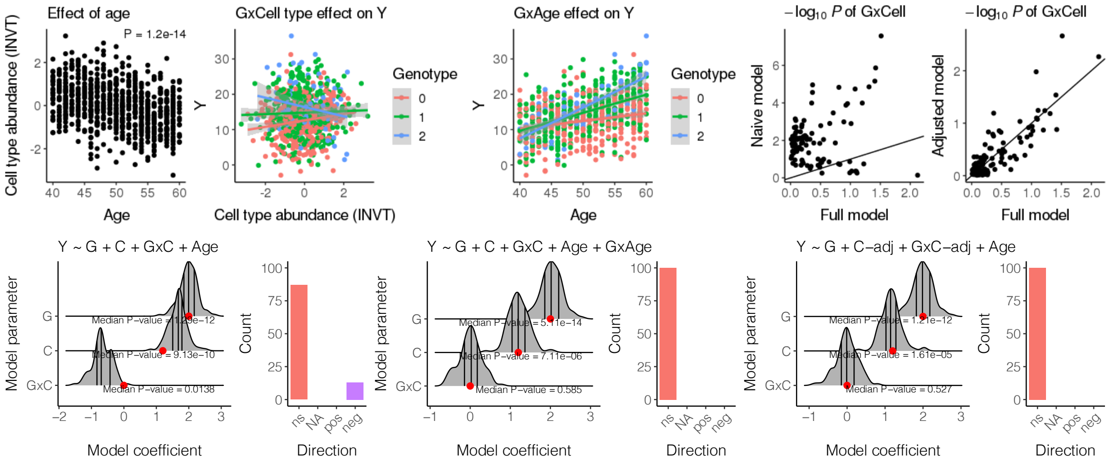


### The effect of age on cell type abundance - beta = 0

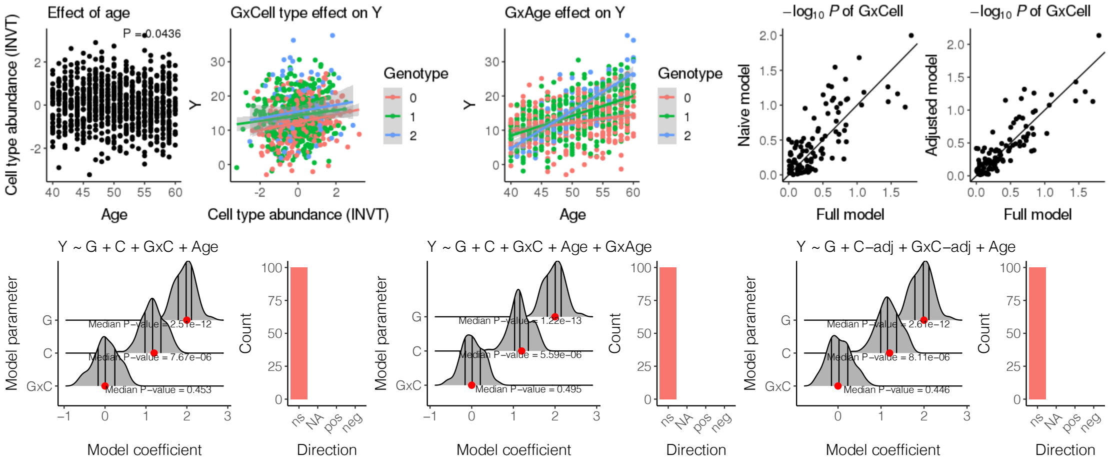


### The effect of age on cell type abundance - beta = 0.25

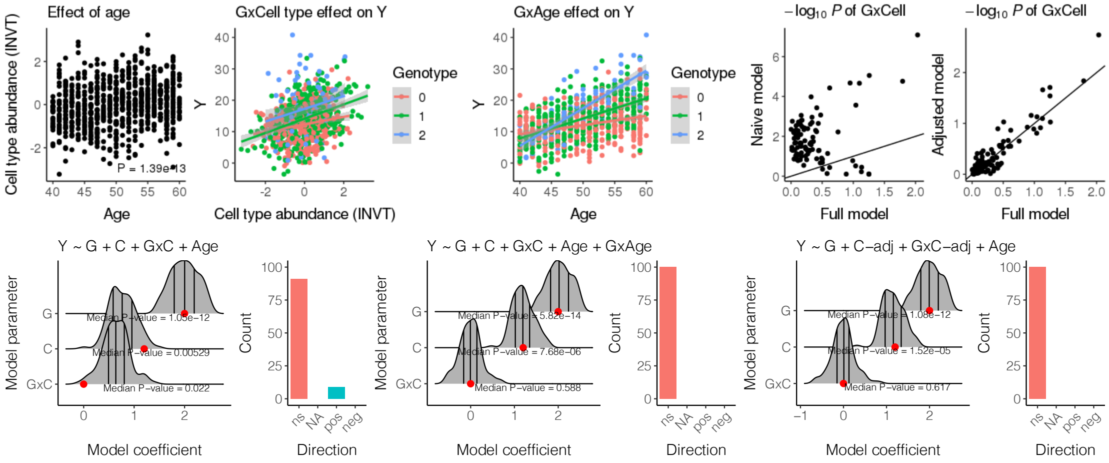

The effect of age on cell type abundance - beta = 0.5

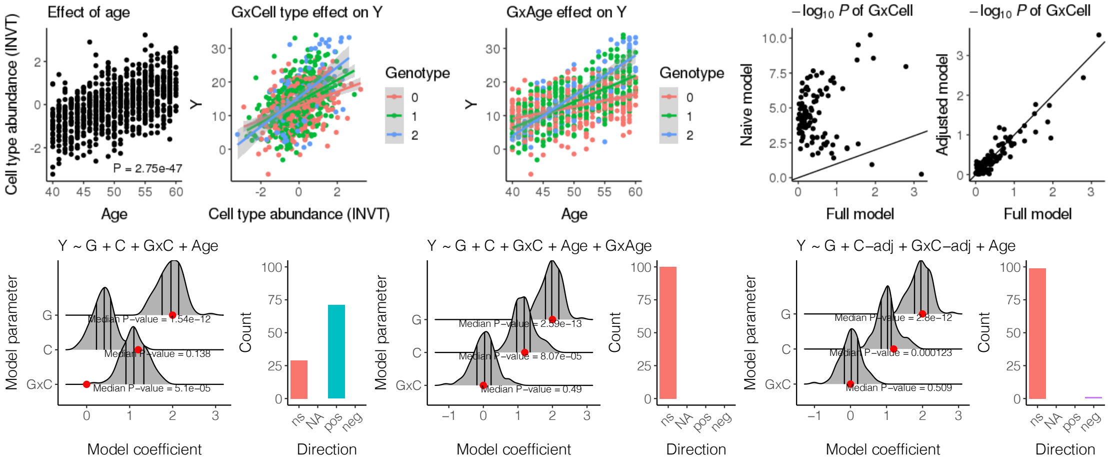


Conclusion

If there are no significant iQTL effects in the presence of a strong GxConfounder effect, we observe variability in *P*-values of the between the GxCell type effect from the full model and adjusted model. Results from the naive model are, in large, overestimated and leading to many false positives, when there is a very strong effect of a confounder on cell type abundance, which in practice is not typical.

### Summary

In practice, cell type abundance is the probably the factor with the strongest influence on the association between genotype and molecular phenotype (*i.e.,* strongest iQTLs). Thus, scenarios in this simulation leading to large scale over- or underestimation in the presence of other GxConfounder effects, where the confounder also affects cell type abundance, are not typical. Correcting cell type abundances for confounders and using normalized residuals in the model rather seems to reduce the study power in most typical scenarios.
